# Nonenzymatic, prebiotic aminoacylation couples chirality of RNA and protein

**DOI:** 10.1101/2024.07.29.605638

**Authors:** Joshua A. Davisson, Evan M. Kalb, Isaac J. Knudson, Alanna Schepartz, Aaron E. Engelhart, Katarzyna P. Adamala

**Affiliations:** University of Minnesota Department of Genetics, Cell Biology and Development, Minneapolis, MN, USA; Department of Chemistry, University of California, Berkeley, CA, USA; Center for Genetically Encoded Materials, University of California, Berkeley, CA, USA

## Abstract

Life as we know it depends on the homochirality of nucleic acids and proteins. However, there is no widely accepted explanation for why life uses only D-sugars for nucleic acids and L-amino acids for proteins. Here we demonstrate a prebiotically plausible method of nonenzymatic aminoacylation in a water ice-eutectic phase. These reactions produce high yields of aminoacyl-tRNAs, which are active in translation. Surprisingly, we discovered these nonenzymatic aminoacylation conditions were stereoselective, favoring coupling of amino acids and RNA of “opposite” L- and D- configurations. D-RNA shows greater aminoacylation yields for L-amino acids. The opposite was true for L-RNA, which had greater yields with D-amino acids. Nucleic acid backbone chirality influencing stereoselectivity of aminoacylation presents the missing link in the origin of modern biochemistry. This phenomenon provides insight into the chirality of the RNA world, and helps to explain the “opposite” stereochemistry of modern biomolecules.

## Introduction

Stereoselectivity is one of the hallmarks of biological life^1,2^. Interestingly, the two major polymers used by extant life are of the “opposite” stereochemistry: all chiral proteinogenic amino acids have an L configuration, while all sugar backbones of RNA and DNA possess the D configuration. While it is easy to understand why enantioselectivity is necessary to life in general, there has been no widely accepted explanation for why the two major biopolymers possess opposite configurations. The theories to explain the origin of homochirality invoke polarized light^3^, surfaces^4^, or reaction conditions^5^ as a source of the original symmetry breaking. However, none of these theories provide evidence to explain why the two major classes of molecules ended up with the opposite configurations^6^. One would expect that any physical method of symmetry breaking would influence all substrates similarly, resulting in all-L or all-D biopolymers.

Another crucial question is how the RNA-templated synthesis of proteins began. To utilize codons of the genetic code, some form of prebiotic tRNA would need to be charged by specific amino acids. In modern biology, this process is catalyzed by highly conserved, complex enzymes, known as acyltransferases. A simple, non-specific method for enzymatically charging tRNA with amino acids has been developed – a short RNA enzyme called flexizyme, offering a possible intermediate step in the origin of biological aminoacylation^7–9^. Flexizyme-promoted reactions exhibit little stereoselectivity for the chirality of the amino acid substrate^10^. While flexizymes provides a simplified route for charging tRNAs, it is still unclear how aminoacylation might have occurred before the origin of enzymes.

Previous work relied on interstrand transfer of amino acids to demonstrate nonenzymatic aminoacylation^11–13^ (aminoacylation under high pressure being an exception^14^). In these studies, one oligonucleotide is synthesized with an aminoacyl phosphate on the 5’ end. When two strands are annealed, or supported by a bridging oligonucleotide, there is an interstrand transfer of the amino acid from the 5’ end of one strand to the 3’ hydroxyl of the other strand (Figure **1b**). While these studies show stereoselectivity, a truly intermolecular aminoacylation reaction, using a separate amino acid not linked to annealed oligonucleotides, has never been shown.

**Figure 1.**
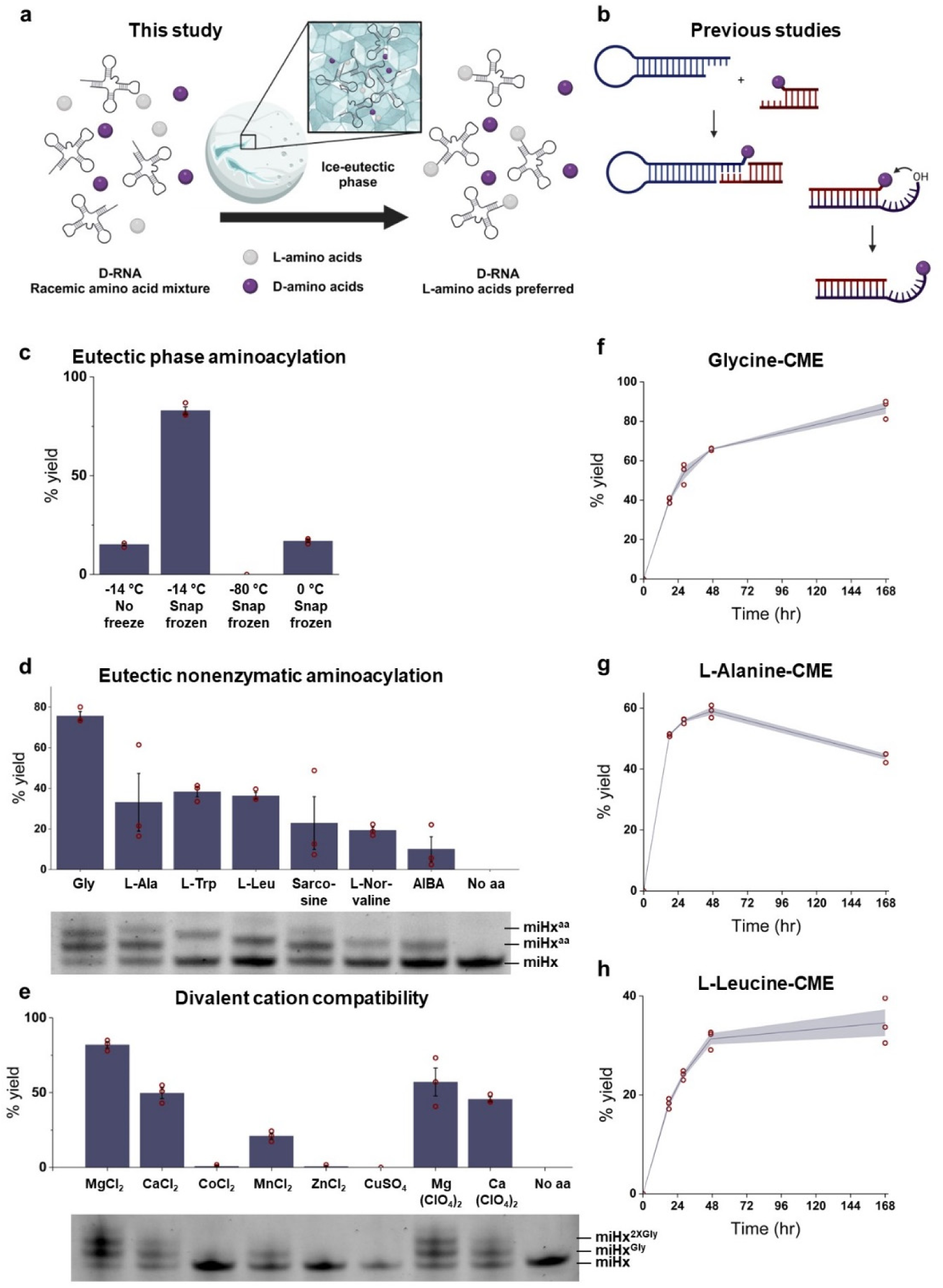
Prebiotically plausible eutectic reaction conditions enable nonenzymatic aminoacylation. **a.** A schematic showing how the Snowball Earth period could facilitate eutectic reaction conditions, including nonenzymatic stereoselective aminoacylation. **b.** A schematic showing previous studies’ methods from achieving nonenzymatic aminoacylation. **c.** Nonenzymatic acylation of glycine-CME to a microhelix (miHx) was performed with both snap freezing, and at different temperatures. Eutectic reaction conditions significantly increased the product yield. Gel analysis in supplementary figures S24-25. **d.** A chemically diverse set of amino acids (all CME esters) were aminoacylated onto a microhelix (miHx) under eutectic conditions. Gel analysis in supplementary figures S6-8. **e.** Nonenzymatic acylation in a eutectic phase is compatible with some, but not all divalent cations. Acylation of miHx with glycine-CME yielded measurable product with Mg(II), Ca(II), and Mn(II), but no acylation is observed in the presence of Zn(II), Cu(II), and Co(II), all cations used at 20 mM. Gel analysis in supplementary figures S11-13. **f-h**. Time courses of nonenzymatic aminoacylation of miHx for glycine-CME (panel **f**), L-alanine-CME (panel **g**), and L-leucine-CME(panel **h**). Purple lines represent the average yield of n=3 replicates. Individual data points are shown using maroon circles. The purple shaded areas show the standard error of the mean for the data set. Gel analysis in supplementary figures S40-44. For panels **c-e**: Purple bars represent the average of n=3 replicates. Maroon circles represent individual data points. Error bars show the standard error of the mean.

Additionally, previous work has relied on aminoacylating oligonucleotides that are not compatible with the translation machinery. There has been no demonstration of charging of full length tRNAs that can be functional in translation, thus no linking of modern biomolecules with prebiotically plausible aminoacylation.

Here we present a scenario that bridges the gap between prebiotically plausible aminoacylation, and the homochiral life based on polymers of the “opposite” chirality. The nonenzymatic aminoacylation conditions that allow for symmetry breaking are compatible with physicochemical conditions of the early Earth, utilizing a water-ice eutectic phase (Figure **1a**).

We demonstrate a crucial step in the origin of codon-based translation, non-enzymatically charging tRNA with amino acids of the “opposite” stereochemistry, and we demonstrate the product is translationally active. We have also shown that those stereoselective aminoacylation conditions are compatible with protocell vesicles, with a variety of divalent cations, and with perchlorate.

## Results and Discussion

### Nonenzymatic aminoacylation in an ice-eutectic phase

During the process of investigating flexizyme-catalyzed aminoacylation reactions in a water ice-eutectic phase^15^, we discovered examples where rapid tRNA aminoacylation occurred in the absence of added flexizyme. All examples of flexizyme-free acylation involved amino acids activated as cyanomethyl esters (CME) and not dinitrobenzyl esters (DBE). To determine if this flexizyme-free aminoacylation was dependent on eutectic conditions, we used miHx, a 22 nt tRNA acceptor stem mimic that can easily establish aminoacylation yields by gel separation. We used miHx and glycine-CME in a flexizyme-free aminoacylation reaction with and without snap freezing and at different temperatures. We found a ∼5 fold increase in acylation yield in a water-ice eutectic phase (snap frozen and then incubated at -14 °C) (Figure **1c**). The -14 °C no freeze samples remained a liquid for the duration of the reaction. The reaction performed at 0 °C with snap freezing quickly melted and also was a liquid for the duration of the experiment. Only the reaction that was snap frozen and then incubated at -14 °C substantially increased acylation yield (Figure **1c**).

We next explored the substrate scope of these nonenzymatic reactions, using cyanomethyl esters of proteogenic and nonproteogenic amino acids. We found miHx was acylated in high yield when reacted with the cyanomethyl esters of a variety of α-amino (Figure **1d**). When nonenzymatic aminoacylation was tested with different divalent cations, we found Mg(II) to produce the highest acylation yields, followed by Ca(II) and Mn(II) (Figure **1e**). Co(II), Zn(II), and Cu(II) did not produce measurable aminoacylation (Figure **1e**). Aminoacylated miHx was readily formed for glycine-CME, L-alanine-CME, and L-leucine-CME (reaching substantial yields with 24 to 48 hours), and were stable in the eutectic phase, persisting beyond a week at -14 °C, pH 7.5 (Figure **1f-h**).

These experiments show that nonenzymatic aminoacylation is dependent on a water ice-eutectic phase to achieve high yields. Nonenzymatic aminoacylation is compatible with a variety of different divalent cations. The kinetics of the reaction are compatible with both the presence of eutectic conditions during periodic (seasonal) freeze thaw cycles, and with the long-term freeze conditions of the snowball Earth period. Overall, we have demonstrated that acceptor stem RNA can be aminoacylated with a variety of amino acids, both proteinogenic and non-proteogenic, under eutectic conditions via a nonenzymatic reaction. This reactivity provides a plausible route to explain the origin of coupling of between an amino acid and RNA - a necessary step in the origin of Central Dogma. The activating group on the amino acid, the cyanomethyl ester, is prebiotically plausible^16^, and it has been previously used in RNA catalyzed aminoacylation scenarios^7,17^.

### Nonenzymatic-acylated tRNAs are functional in translation

Next, we wanted to assess if the nonenzymatic aminoacylation only occurs with miHx RNA, or if full length tRNAs could also be acylated under the same conditions. Using intact tRNA LCMS^14^, we confirmed that nonenzymatic charging using glycine-CME and alanine-CME produced aminoacylated tRNA^fMet^ (Figure **2a-b**). To test if these acylated tRNAs were functional in translation, glycine-CME and alanine-CME were used to aminoacylate formylmethionine tRNA (tRNA^fMet^) in a nonenzymatic reaction. These acyl-tRNAs were added to a PURExpress (ΔtRNA, Δaa) *in vitro* translation reaction, with a DNA template encoding the HiBiT peptide^18^ and all necessary amino acids and tRNA except for tRNA^fMet^ and methionine.

**Figure 2.**
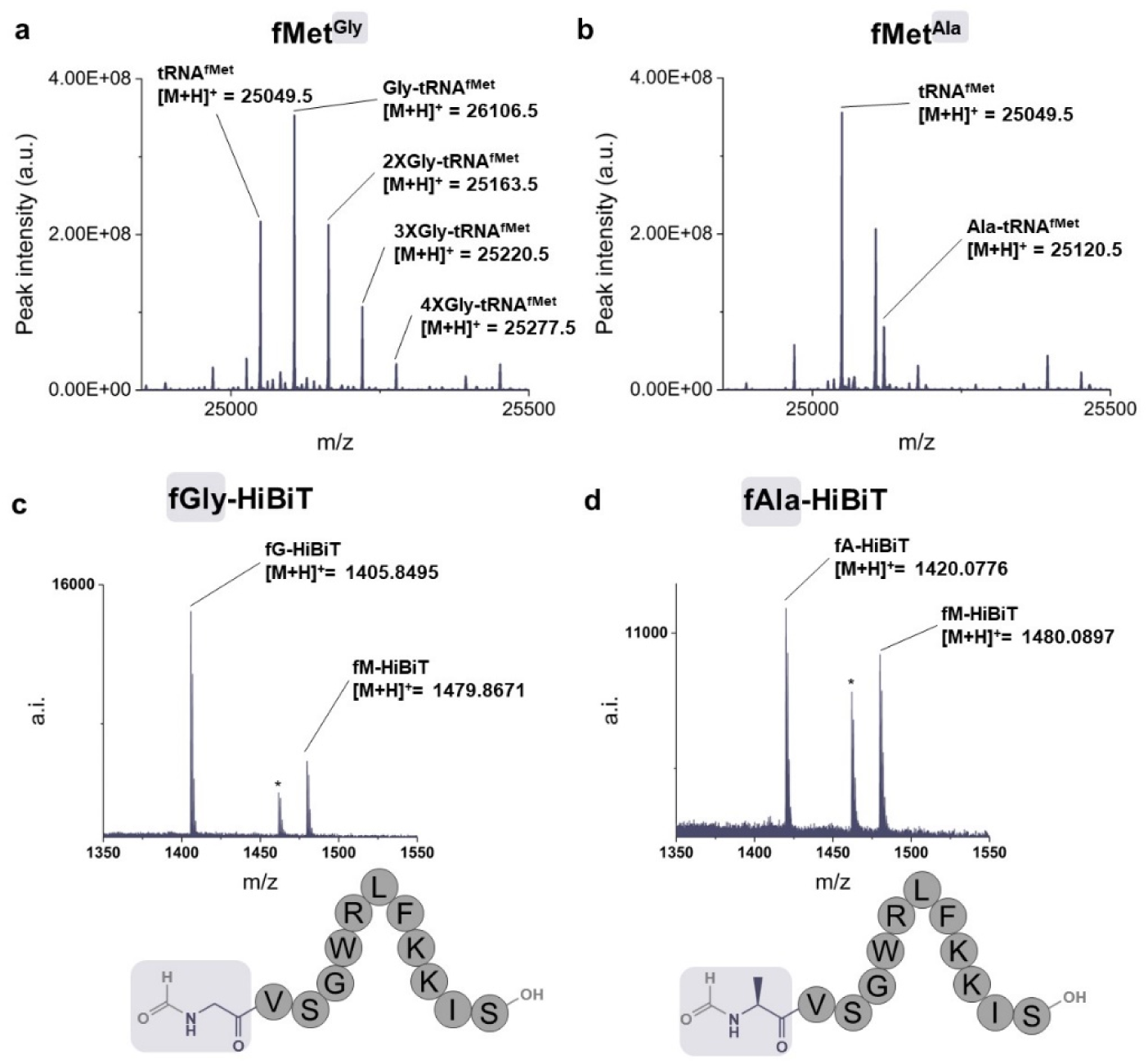
Full length tRNA aminoacylated under nonenzymatic eutectic conditions is active in translation. **a-b.** Intact tRNA LCMS confirms full length tRNA fMet is aminoacylated with glycine-CME and L-alanine-CME using eutectic phase conditions. **c-d.** tRNA fMet was acylated with glycine-CME or L-alanine-CME and used in an *in vitro* translation reaction to initiate translation of a HiBiT peptide. Mass spectrometry was used to detect peptide products matching glycine- or alanine-initiated peptides in the translation reaction. Labels indicate peaks for gly/L-ala-initiated HiBiT and fMet-initiated HiBiT. Asterisks show -18 Da dehydration product from fMet-HiBiT. HiBiT luminescence data shown in supplementary figures 64-65. Panels **a**, **c** show experiments with glycine, and panels **b**, **d** show experiments with L-alanine.

Using MALDI mass spectrometry, we directly detected peptides with masses matching glycine- and alanine-initiated HiBiT peptides among the *in vitro* translation reaction products (Figure **2c-d**). Nonenzymatic charging reactions produce aminoacyl tRNA active in translation. These data demonstrate a plausible route for aminoacylation of both full size modern tRNA, as well as shorter possible prebiotic precursors to the tRNA (as demonstrated in the miHx experiments).

### Origins of an L-amino acid world

To investigate if nonenzymatic acylation had a chiral preference for amino acids, we activated both the L and D configurations of leucine. To our surprise, we discovered that the L-α-amino acid acylation yields with miHx RNA were generally higher than yields of the analogous D-α-amino acid (performed using “natural” D-ribose miHx) (Figure **3c**). Under identical conditions, average yields of acyl-miHx (determined by gel estimation) with L-α-amino acids exceeded those of D-α-amino acids. This presented an opportunity to investigate the possible mechanisms of prebiotic symmetry breaking,

**Figure 3.**
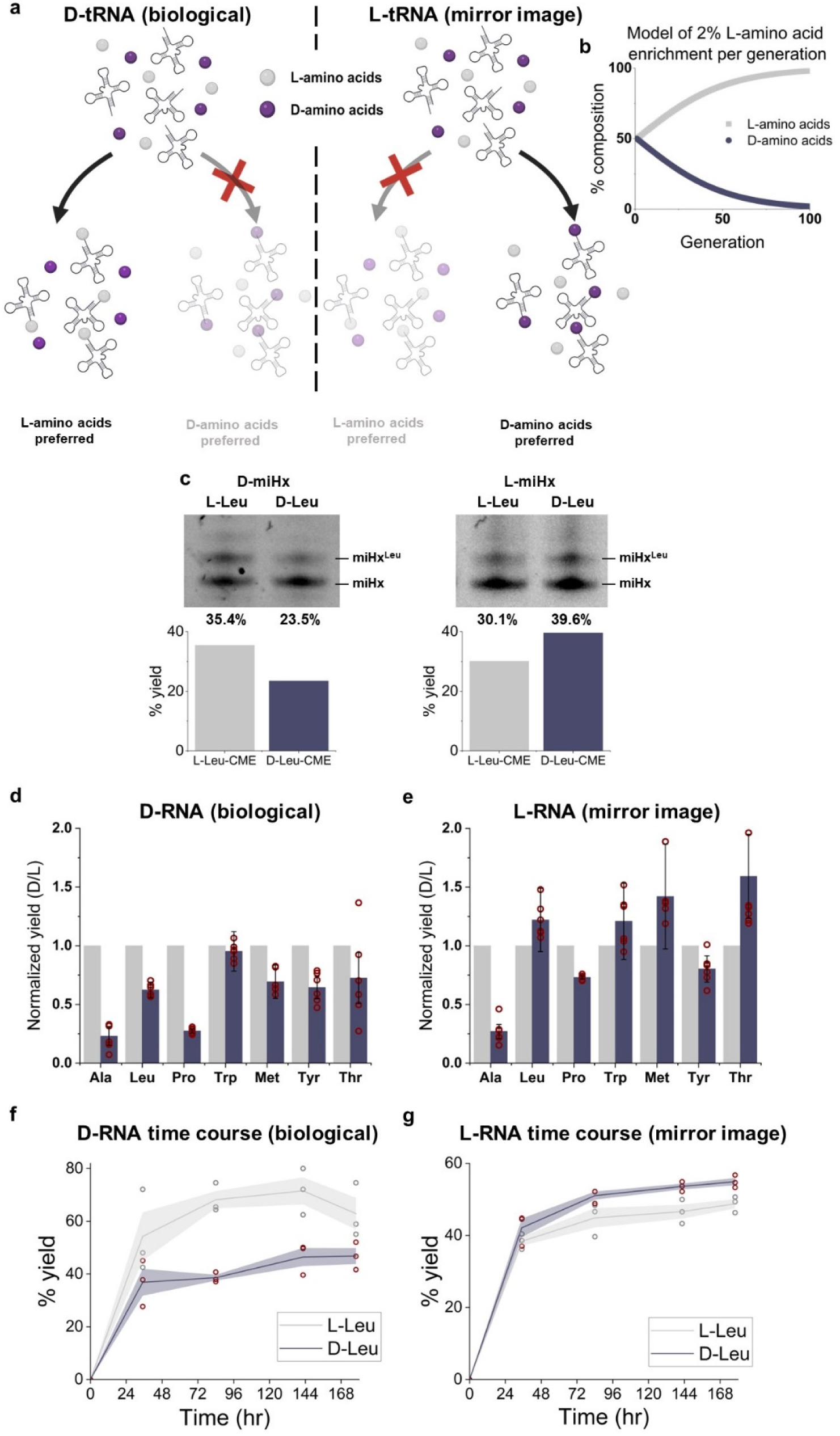
Nonenzymatic aminoacylation is stereoselective between L and D-amino acids and stereoselectivity is dependent on the nucleic acid chirality. **a.** Schematic showing nonenzymatic eutectic phase reactions exhibiting stereoselectivity. The schematic shows biologically relevant D-RNA preferentially being acylated with L-amino acids, while L-RNA (mirror image life) being preferentially acylated with D-amino acids. **b.** A model showing that 2% enrichment of L-amino acids per generation, assuming a racemic starting mixture, results in completely enantiopure composition in 100 generations. **c.** A representative gel showing L and D-leucine-CME being acylated onto either D-RNA or L-RNA miHx. The gel shows D-miHx has increased aminoacylation yields for L-leucine over D-leucine. L-miHx has increased aminoacylation yields for D-leucine over L-leucine. Bar graphs show yield calculated from the intensity of gel bands. **d-e.** D-amino acid acylation yields (in purple) relative to normalized L-amino acid acylation yields (in gray) for both D- and L-RNA. For the biologically relevant D-RNA, D-amino acid acylation yields are lower than L-amino acid acylation yields. For the mirror image L-RNA, D-amino acid acylation yields are generally greater than L-amino acid acylation yields. Purple and gray bars represent the average of n=6 replicates. Maroon circles represent individual data points. Error bars show the standard error of the mean. Gel analysis in supplementary figures S14-22, 28-39. **f-g.** Aminoacylation time courses charging D-miHx or L-miHx using L-Leu-CME and D-Leu-CME. Maroon circles represent individual data points. Shaded areas show the standard error of the mean. Gel analysis in supplementary figures S65-71.

To test if eutectic nonenzymatic aminoacylation was generally stereoselective, we activated a total of seven L and D amino acids with cyanomethyl esters. When aminoacylating D-ribose RNA (the biologically relevant enantiomer), we observed that L-α-amino acids achieved higher acylation yields compared to D-α-amino acids (Figure **3d**). We wondered if the stereoselectivity of nonenzymatic aminoacylation was dependent on the chirality of the RNA backbone. To test this, we synthesized the L-ribose enantiomer of miHx and performed an analogous set of acylation reactions.

We tested the same L- and D-amino acids with L-ribose RNA (mirror image to extant terrestrial life). We found that the opposite was true: with L-RNA, D-α-amino acids generally achieved higher yields compared to L-α-amino acids (Figure **3e**). We also assessed formation over time of L and D-aminoacyl L and D-miHx (Figure **3f-g**). To discern if stereoselectivity was imparted by eutectic phase conditions, we also performed experiments in a liquid, noneutectic phase at - 14 °C. Although aminoacylation yields were much lower, stereoselective aminoacylation persisted without a water ice-eutectic phase (Supplementary figures S72-76).

The experiments elucidate the possible origin of the relationship between two key polymers used in life: nucleic acids and proteins. All modern terrestrial life uses D-ribose/deoxyribose for nucleic acid chains, and L-amino acids for proteins. The stereoselectivity of nonenzymatic charging reactions presented here possibly represents an early link between nucleic acid and protein chirality that has formed the foundation of all extant life. Our data supports the possibility of “opposite chirality” of those two major polymers. Whether L-amino acids with D-sugars, or D-amino acids with L-sugars, the nonenzymatic aminoacylation favors coupling between the amino acids and RNA of opposite configurations. The prebiotic reactions do not need to be completely stereoselective^5,19^. Even two percent enrichment (less than we have observed with most tested amino acids) will result in enantiopure products over only 100 generations. (**Figure 3b**).

### Nonenzymatic charging is compatible with early protocells and prebiotic conditions

The prevailing hypothesis is that the origin of life was a multi-step process, with several elements with different chemical constraints having to come together to give rise to the Last Universal Common Ancestor. Particularly, the emergence of DNA replication, transcription, and translation had to be compatible with the protocell lipid membranes, as encapsulation supports individuality as a fundamental element of Darwinian selection. To test whether the nonenzymatic RNA aminoacylation reported here is compatible with this fundamental element of prebiotic evolution, we tested whether the reaction would proceed in the presence of liposomes composed of fatty acids. Fatty acids are prebiotically plausible membrane components; however, these membranes are unstable in the presence of higher amounts of unchelated divalent cations^20,21^. High cation concentrations are required in many prebiotically plausible nucleic acid replication and catalysis. It has been previously demonstrated that the high divalent cation concentrations required for nonenzymatic RNA replication could be combined with fragile fatty acid liposomes using citrate, a common and prebiotically plausible metabolite^20^. Inspired by this result, we tested if nonenzymatic RNA aminoacylation could be compatible with fatty acid liposomes using citrate.

Nonenzymatic aminoacylation acylation in the presence of oleic acid liposomes and citrate moderately decreased acylation yield (Figure **4a**), but importantly liposomes were stable and had expected encapsulation rates despite a freeze-thaw cycle included in a eutectic phase reaction (Figure **4b, e**). We next tested the effect of diluting reagent concentrations on the reaction. We found that even diluting the reaction to 20 times its original volume before initiating the reaction, acylation yields were practically unchanged. This reduced RNA concentrations to 1.25 μM and MgCl_2_ concentrations to 1 mM. Intrigued with how well nonenzymatic charging worked at low concentrations of magnesium, we further reduced the MgCl_2_ concentration down to adding no additional MgCl_2_ yet observed no effect on acylation yield for glycine-CME and alanine-CME (Supplementary figure S26-27).

**Figure 4.**
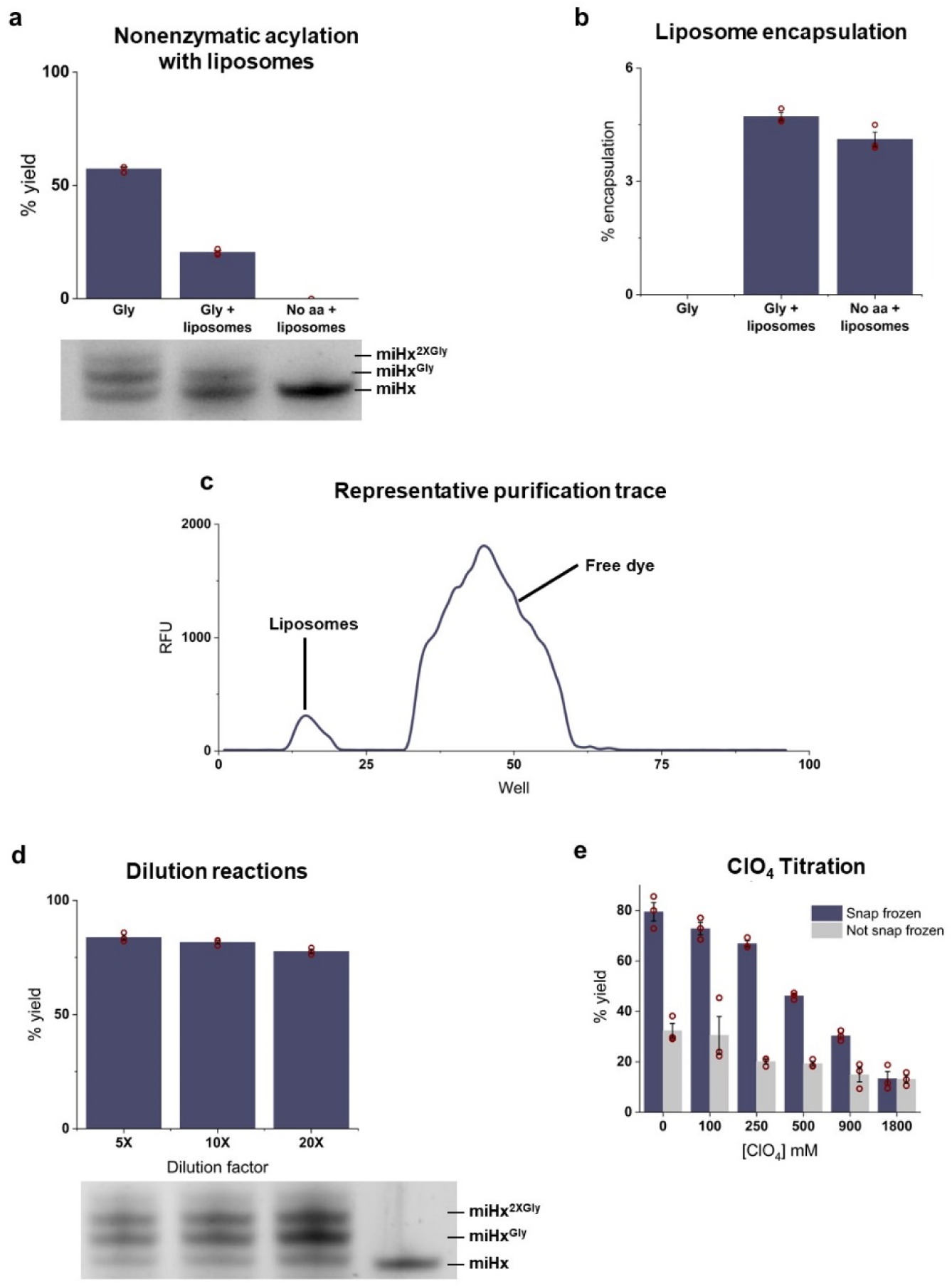
Nonenzymatic aminoacylation is compatible with a variety of prebiotic conditions, including the presence of protocells, low magnesium, and high perchlorate concentrations. **a.** Nonenzymatic acylation reactions are compatible with protocells and citrate chelation. Aminoacylation still occurred when nonenzymatic eutectic acylation reactions of glycine-CME onto miHx were performed in the presence of oleic acid liposomes, with citrate chelation to stabilize liposomes in the presence of MgCl_2_. Gel analysis in supplementary figure S23. **b.** Liposomes remain intact after a freeze and thaw cycle during the eutectic reaction, in presence of citrate and magnesium. Bar graphs show encapsulation of small molecule calcein dye in liposomes from reactions shown on panel **a**. **e.** Representative trace of size exclusion liposome purification. **c.** Low concentrations of reaction components do not decrease efficiency of nonenzymatic charging. Eutectic nonenzymatic acylation reactions were diluted with water up to 20 times in volume (from 10 μL to 200 μL) before being snap frozen and incubated at -14 °C. Gel analysis in supplementary figures S9-10. **f.** NaClO_4_ concentration was titrated to test the resistance of nonenzymatic charging to increasing perchlorate concentration. Higher concentrations of perchlorate prevented the reactions from staying frozen, so the acylation was performed both with and without snap freezing as a control. Purple bars represent reactions snap frozen before incubation at -14 °C while gray bars indicate the reaction was not snap frozen and remained liquid with no eutectic phase. Gel analysis in supplementary figures S1-5. For panels **a-d, f**: Purple bars represent the average of n=3 replicates. Maroon circles represent individual data points. Error bars show the standard error of the mean.

Inspired by RNA and ribozymes’ inherent ability to resist strong chaotropic ions such as ClO_4_ ^-^ present on Mars^22^, we tested nonenzymatic acylation in the presence of increasing concentrations of NaClO_4_. We found nonenzymatic aminoacylation was still observable at 1.8 M NaClO_4_, far too concentrated for protein enzymes to remain functional (Figure **4f**). However, the highest ClO_4_ ^-^ concentrations inhibited freezing of the reaction. We ran reactions that were not snap frozen and remained liquid to control for a liquid vs an ice-eutectic phase. In both cases, acylation was still present, although at reduced yields, at 1.8 M ClO_4_ ^-^. These experiments show nonenzymatic aminoacylation could occur in environments too denaturing for protein enzymes.

## Conclusions

Here we demonstrated how a prebiotically plausible method of aminoacylation, without the use of enzymes, can produce translationally active acyl-tRNAs. This nonenzymatic charging works with a wide variety of amino acids. The eutectic phase reaction results in rapid product formation, and the delicate acyl bond exhibits long-term stability under the reaction conditions. We showed how the ice-eutectic phase is critical for these substantial acylation yields, even at low reagent concentrations or minimal divalent cation concentrations. We demonstrated how nonenzymatic charging is compatible with prebiotically relevant fatty acid protocells. We also demonstrated that nonenzymatic acylation reactions are stereoselective, depending on the chirality of the nucleic acid, and amino acid being charged. D-RNA favors L-amino acid acylation, the biologically relevant stereoisomers of both polymers. This highlights a potentially ancient relationship between nucleic acid and protein chirality, where a D-RNA world might have produced the L-amino acid world of extant biology. This stereochemical relationship also raises possibilities of alternative starts to life, where an L-RNA world selects for a D-amino acid protein world. This is the first experimental evidence pointing to simultaneous, interdependent origin of homochirality of both major classes of biopolymers.

Our work demonstrated experimental evidence for the origin of the “opposite” configuration of major biopolymers of life, demonstrating plausible scenario for evolution of L-sugars & D-peptides biochemistry.

## Materials and Methods

### Synthesis of activated amino acids

All amino acids used were activated with a cyanomethyl ester and were synthesized using previously published method^23^. 0.83 mmol of *N*-butoxycarbonyl (Boc)-protected amino acid was dissolved in 1 mL of dimethyl formamide (DMF), 0.3 mL of chloroacetonitrile, and 0.26 mL of diisopropylethylamine. This mixture was stirred at room temperature for 16 hours. To work up the reaction, the reaction mixture was diluted with 15 mL of ethyl acetate before being washed twice with 1 M HCl, twice with saturated NaHCO_3_, and five times with saturated NaCl. The organic layer was dried using MgSO_4_. The organic layer was concentrated by rotary evaporation. Removal of the boc group was accomplished by adding 2 mL of 4 M HCl in dioxane and stirred at room temperature for 30 minutes. This mixture was then concentrated by rotary evaporation and precipitated using cold diethyl ether. The solvent was removed using rotary evaporation and the precipitate was washed an additional two times with cold diethyl ether. Any remaining solvent was removed under vacuum overnight. Reaction products were confirmed using ^1^H NMR (Supplementary figures S45-62).

### Preparation of RNA

DNA templates for D-microhelix (miHx) were prepared via annealing of oligonucleotides purchased from Integrated DNA Technologies (IDT). DNA template for tRNA^fMet^ was made using three successive PCR reactions^24^. The PCR reaction consisted of 1X Q5 DNA polymerase buffer (New England Biolabs), 0.75 mM dNTPs, and 0.02 U/μL Q5 DNA polymerase. First fMet-F and fMetR2 were used in the first round of PCR at 5 μM final concentrations with 3 cycles.

The extension product was used in the next round of PCR using 5 μM of T7-F and fMetR2 with 3 cycles. The extension product of this reaction was used in the next round of PCR using 0.5 μM of T7-F and fMetR3 with 15 cycles. Thermal cycles were 1. 95 °C for 1 min, 2. 95 °C for 40 seconds, 3. 50 °C for 40 sec, 4. 72 °C for 40 sec, 5. 4 °C infinite. Steps 2-4 were repeated for each cycle.

For the D-miHx and tRNA^fMet^ were transcribed using a T7 transcription reaction as previously described^25^. Briefly, the reaction consisted of 1X template, 1X Homemade NEB Buffer, 8 mM GTP, 4 mM A/C/UTP, 0.005X phosphatase 25 ng/μL, 1 μM T7 RNAP, RNAse inhibitor 0.4 U/μL. The transcripts were digested with Turbo DNAse (New England Biolabs) at 37 °C for 20 minutes before being purified using a Monarch RNA purification kit using 2X volume ethanol (New England Biolabs). Concentration determined using a Nanodrop ND-1000, diluted to 250 μM, and stored in aliquots at -80 °C. Sequence table is included in supplementary table S1. L-miHx was purchased from Biosynthesis (Lewisville, TX)

### Ice eutectic reactions

Reaction setup was adapted from flexizyme reaction conditions and tC19z reaction conditions as described previously^26,27^. Reaction conditions: 5mM HEPES-KOH pH 7.5, 25 μM miHx or tRNA, 20 mM MgCl2, 2.5% DMSO. A mixture of 1 μL of HEPES-KOH pH 7.5, 1 μL of 250 μM miHx, and 4 μL of water was annealed at 95 °C for 2 minutes then slowly cooled to 25 °C (in 10 °C intervals for 1 minute each). Next, 2 μL of 100 mM MgCl2 was added and the reaction was placed on ice. Then, 2 μL of 25 mM activated amino acid (in a 12.5% DMSO-water solution) was added and mixed thoroughly. The reactions were snap frozen in liquid nitrogen (unless otherwise specified) and incubated in an ethylene glycol-water bath at -14 °C for 84 hours (unless otherwise specified). To quench, the reaction mixture was diluted with 40 μL of 300 mM NaOAc pH 5.2 and 100 μL of 100% ethanol and mixed. The quenched reaction was centrifuged at 15,000 x g at room temperature for 15 minutes.

For the oleic acid liposomes experiments, the MgCl_2_ was reduced to 10 mM and the reaction contained 40 mM citrate added after the MgCl_2_, but before the amino acid.

For the ClO_4_ reactions, less water was added to the initial HEPES/miHx and the ClO_4_ solution was added after the MgCl_2_, but before the amino acid.

### Acid PAGE gel shift assay

The precipitated RNA pellet was resuspended in 4 μL of 10 mM NaOAc and 12 μL of acid PAGE loading buffer as described previously^26^. 3.5 μL of this mixture was loaded per lane. The reactions were run at 125 V for 2.75 hours on a denaturing (6 M urea) 20% (w/v) polyacrylamide gel in 50 mM NaOAc pH 5.2 running buffer (using the Biorad Mini-PROTEAN tetra gel, 1 mm gel thickness). The gels were stained with Sybr Gold (Invitrogen) and visualized using an Omega Lum gel imager (Aplegen). MiHx bands were quantified using ImageJ.

We observed gel shift assays and intact tRNA LCMS show miHx and tRNAs with multiple acylations for some amino acids tested (e.g. Gly, Leu, and Met). While previous work has shown that bis-acyl-tRNA on the 2’ and 3’ terminal hydroxyls are active substrates for translation, besides the 3’-OH, it is unknown where on the miHx/tRNA these amino acids are being appended^28^.

### Analysis of tRNA acylation products using intact tRNA LC-HRMS

Using a nonenzymatic reaction with 5 μL of 250 μM tRNA, glycine, alanine, or a negative control with only DMSO was acylated for 84 hours. The reactions were precipitated by adding 200 μL of 300 mM NaOAc pH 5.2 and 500 μL of 100% ethanol. These tubes were mixed and centrifuged at 15,000 x g for 15 minutes. The supernatant was removed and 250 μL of 70% ethanol was added to each tube and centrifuged at 15,000 x g for 10 minutes. The supernatant was removed and this step was repeated. The supernatant was removed and the pellet was allowed to dry on the benchtop with the cap open covered with a kimwipe for 5 minutes. The tube was snap frozen in liquid nitrogen and stored at -80 °C.

Intact tRNA LC-MS was performed as previously reported^29^. Samples were resuspended in water before analysis using intact tRNA LC-MS. In general, 20 pmol of tRNA was injected to ensure adequate signal in the UV and ion chromatograms. Samples were resolved on a ACQUITY UPLC BEH C18 Column (130 Å, 1.7 μm, 2.1 mm X 50 mm, Waters part # 186002350, 60 °C) using an ACQUITY UPLC I-Class PLUS (Waters part # 186015082). The mobile phases used were (A) 8 mM TEA, 80 mM HFIP, 5 μM EDTA (free acid) in 100% MilliQ water; and (B) 4 mM TEA, 40 mM HFIP, 5 μM EDTA (free acid) in 50% MilliQ water/50% methanol. Samples were eluted using a flow rate of 0.3 mL/min. The elution method began with Mobile Phase B at 22%; the fraction of Mobile Phase B increased linearly to 40% B over 10 min and from 40 to 60% B over 1 min and was then held at 60% B for 1 min. The fraction of Mobile Phase B was then decreased linearly from 60 to 22% B over 0.1 min and held at 22% B for 2.9 min to equilibrate the column. For more difficult separations, the method was extended as needed. For tRNA acylation, the linear gradient of Mobile Phase B from 22% to 40% was extended to 20 min. The mass of the RNA was analyzed using LC-MS with a Waters Xevo G2-XS Tof (Waters part #186010532) in negative ion mode with the following parameters: capillary voltage: 2000 V, sampling cone: 40, source offset: 40, source temperature: 140 °C, desolvation temperature: 20 °C, cone gas flow: 10 L/h, desolvation gas flow: 800 L/h, 1 spectrum/s.

Expected masses of oligonucleotide products were calculated using https://www.cusabio.com/m-299.html#a02 or OligoCalc (for OligoCalc, the mass of H2O must be added to accurately account for the full molecular weight of the triphosphate).8 Deconvoluted mass spectra were obtained using the MaxEnt software (Waters Corporation). Acylation yields were estimated in two ways: In most cases, the relative absorbance of the chromatographically resolved peaks representing acyl- and non-acylated tRNA at 260 nm was adequate to estimate acylation yield. In some cases, low acylation obscured clear A260 signals or multiple UV-active species eluted simultaneously. In these cases, extraction of the expected major ion for acylated-MH, over the range of a single ionization envelope, was compared to an equivalent extracted ion chromatogram for the unacylated-MH. A detailed description of these methods is found in previous reports.6,7

### *In vitro* translation reactions

Using a nonenzymatic reaction with 5 μL of 250 μM tRNA, glycine, alanine, or a negative control with only DMSO was acylated for 84 hours. The reactions were precipitated by adding 200 μL of 300 mM NaOAc pH 5.2 and 500 μL of 100% ethanol. These tubes were mixed and centrifuged at 15,000 x g for 15 minutes. The supernatant was removed and 250 μL of 70% ethanol with 100 mM NaOAc pH 5.2 was added to each tube and vortexed. The tubes were centrifuged again at 15,000 x g for 5 minutes. This step was repeated once more. Finally, 250 μL of 70% ethanol per tube was added then centrifuged at 15,000 x g for 3 minutes. The supernatant was removed and the pellet was allowed to dry on the benchtop with the cap open covered with a kimwipe for 5 minutes. The pellet was resuspended in 3 μL of 1 mM NaOAc pH 5.2 and immediately added to a PURExpress (ΔtRNA, Δaa) (New England Biolabs) reaction containing 2.5 μL solution A, 3.75 μL solution B, and 3 μL resuspended aminoacyl fMet tRNA (12.5 μL total volume).

### MALDI mass spectrometry

The *in vitro* translation reactions were diluted with water (up to 30 μL) and acidified to 0.1% trifluoroacetic acid (TFA). The translated peptides were desalted using millipore C18 zip tips using solutions 1, 2, and 3a (solution 1: 50% ACN in water 0.1% TFA, 2: water 0.1% TFA, 3a: 60% ACN in water 0.1% TFA). The elutions were mixed 1:1 with saturated α-cyano-4-hydroxycinnamic acid and spotted onto the target plate. The Bruker autoflex speed MALDI-TOF was calibrated using an external peptide calibrant.

## Acknowledgements

We would like to thank the Seelig lab for the generous gift of the T7 RNAP expression plasmid. We are grateful to Dr. Jason Heier for many helpful discussions about amino acid syntheses and help interpreting NMRs, and Todd Markowski at the Center for Mass Spectrometry and Proteomics for always being willing to advise on, discuss, and help with mass spectrometry. This work was supported by the NSF Center for Genetically Encoded Materials (CHE-2002182).

## Supplementary information

**Figure S1.**
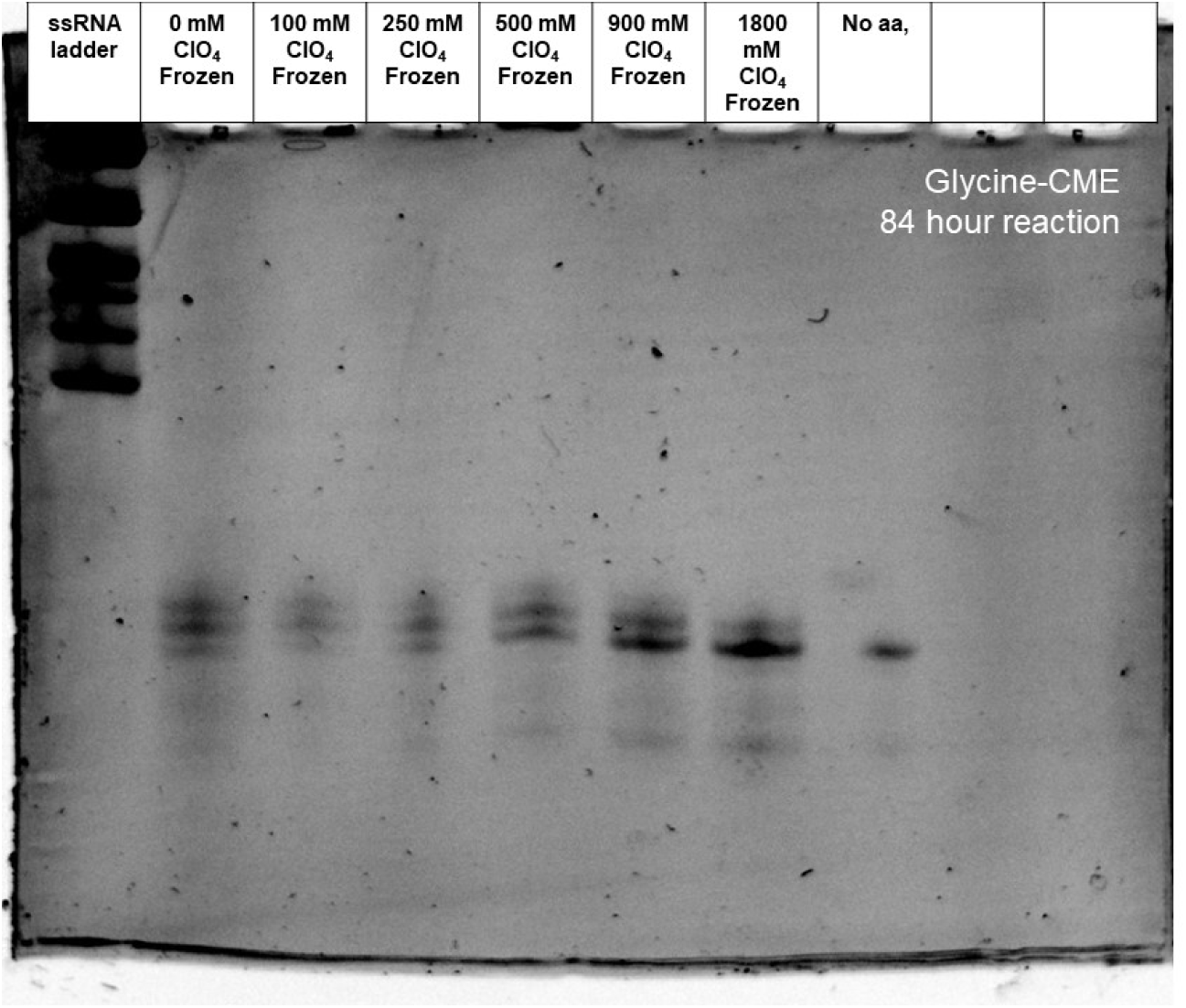
Gel of eutectic nonenzymatic charging reactions with ClO_4_ titration from 0-1800 mM NaClO_4_ charging glycine-CME at pH 7.5 at -14 °C for 84 hours.

**Figure S2.**
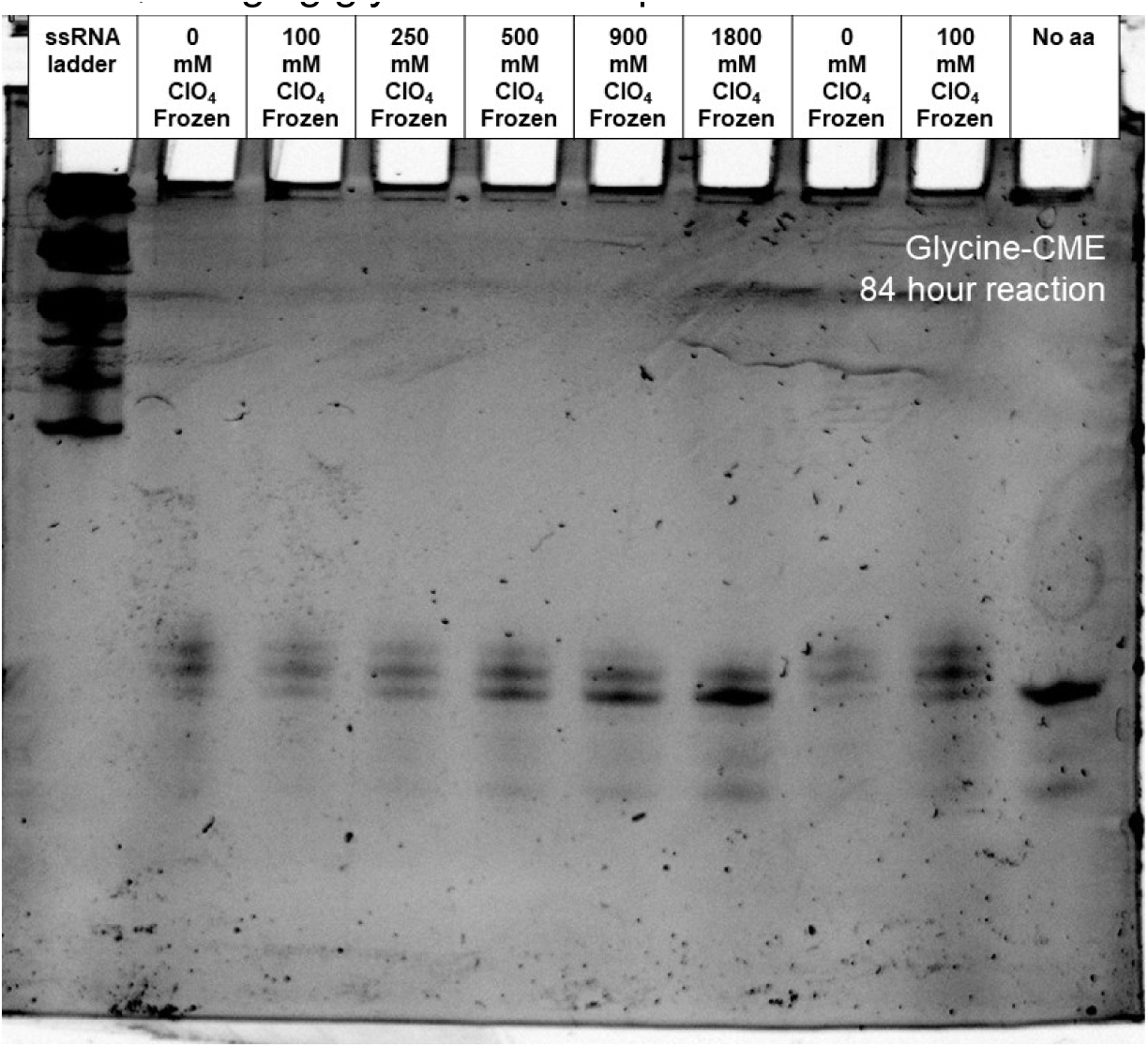
Gel of eutectic nonenzymatic charging reactions with ClO_4_ titration from 0-1800 mM NaClO_4_ charging glycine-CME at pH 7.5 at -14 °C for 84 hours.

**Figure S3.**
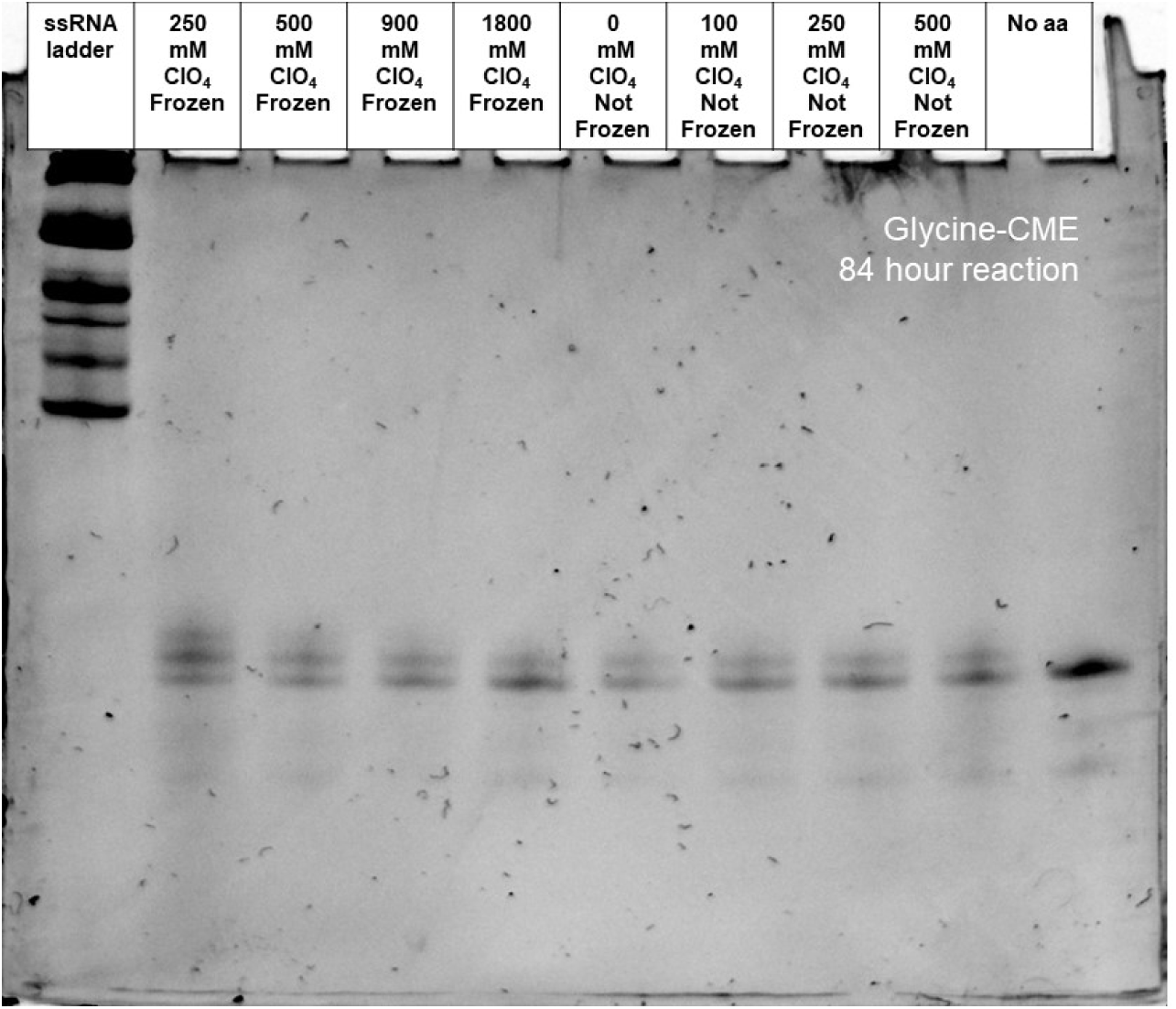
Gel of eutectic and not frozen nonenzymatic charging reactions with ClO_4_ titration from 0-1800 mM NaClO_4_ charging glycine-CME at pH 7.5 at -14 °C for 84 hours.

**Figure S4.**
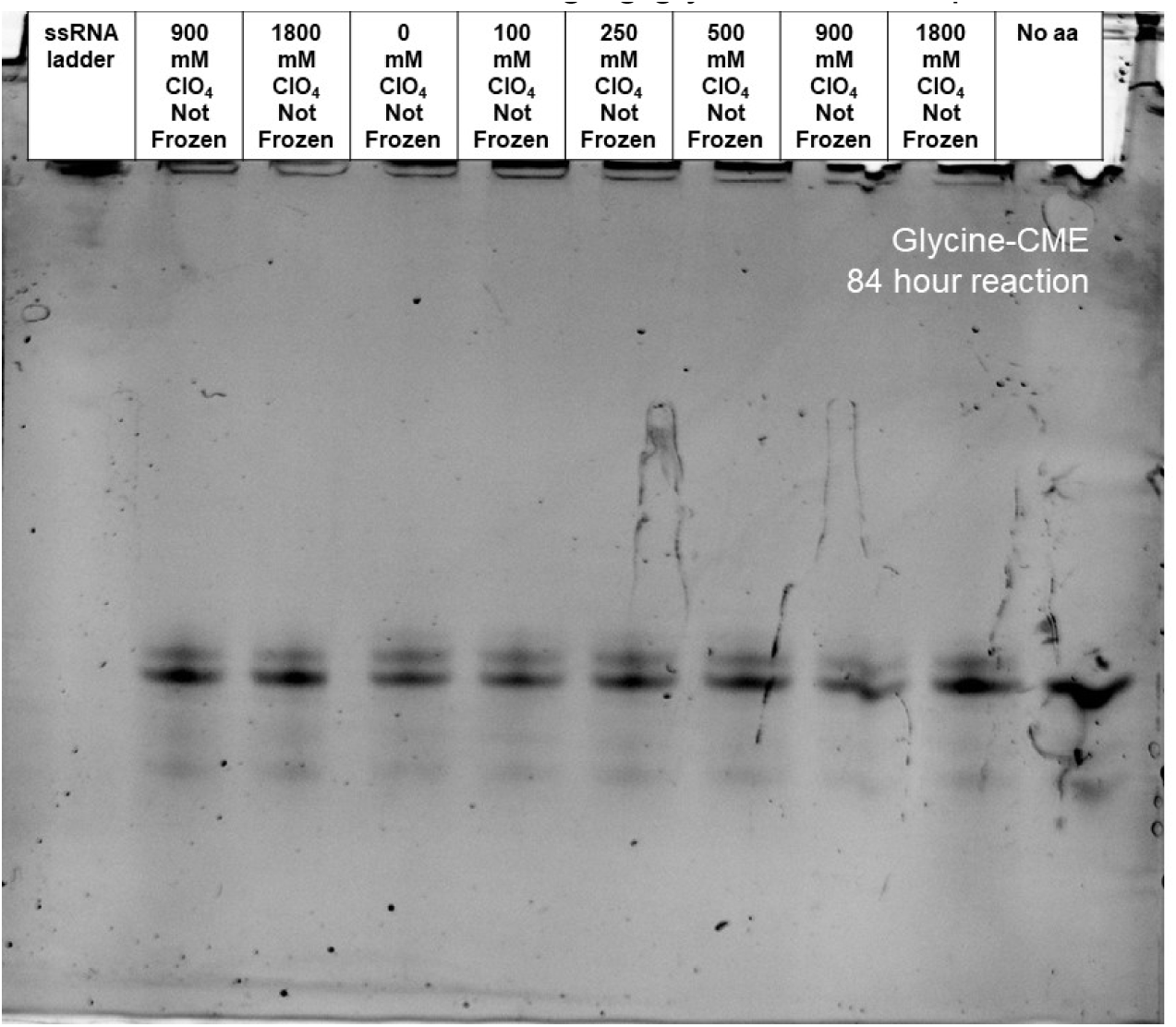
Gel of not frozen nonenzymatic charging reactions with ClO_4_ titration from 0-1800 mM NaClO_4_ charging glycine-CME at pH 7.5 at -14 °C for 84 hours.

**Figure S5.**
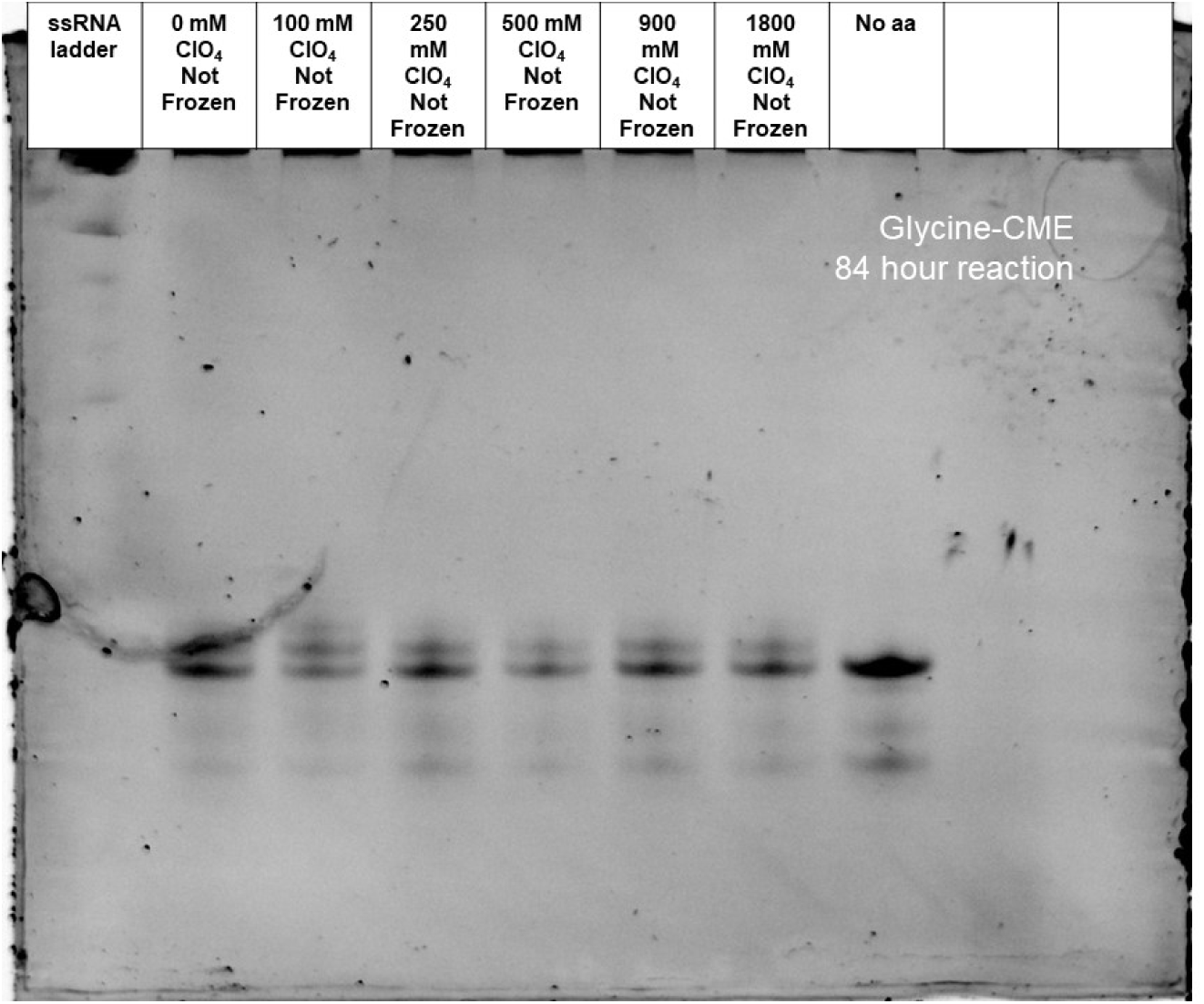
Gel of not frozen nonenzymatic charging reactions with ClO_4_ titration from 0-1800 mM NaClO_4_ charging glycine-CME at pH 7.5 at -14 °C for 84 hours.

**Figure S6.**
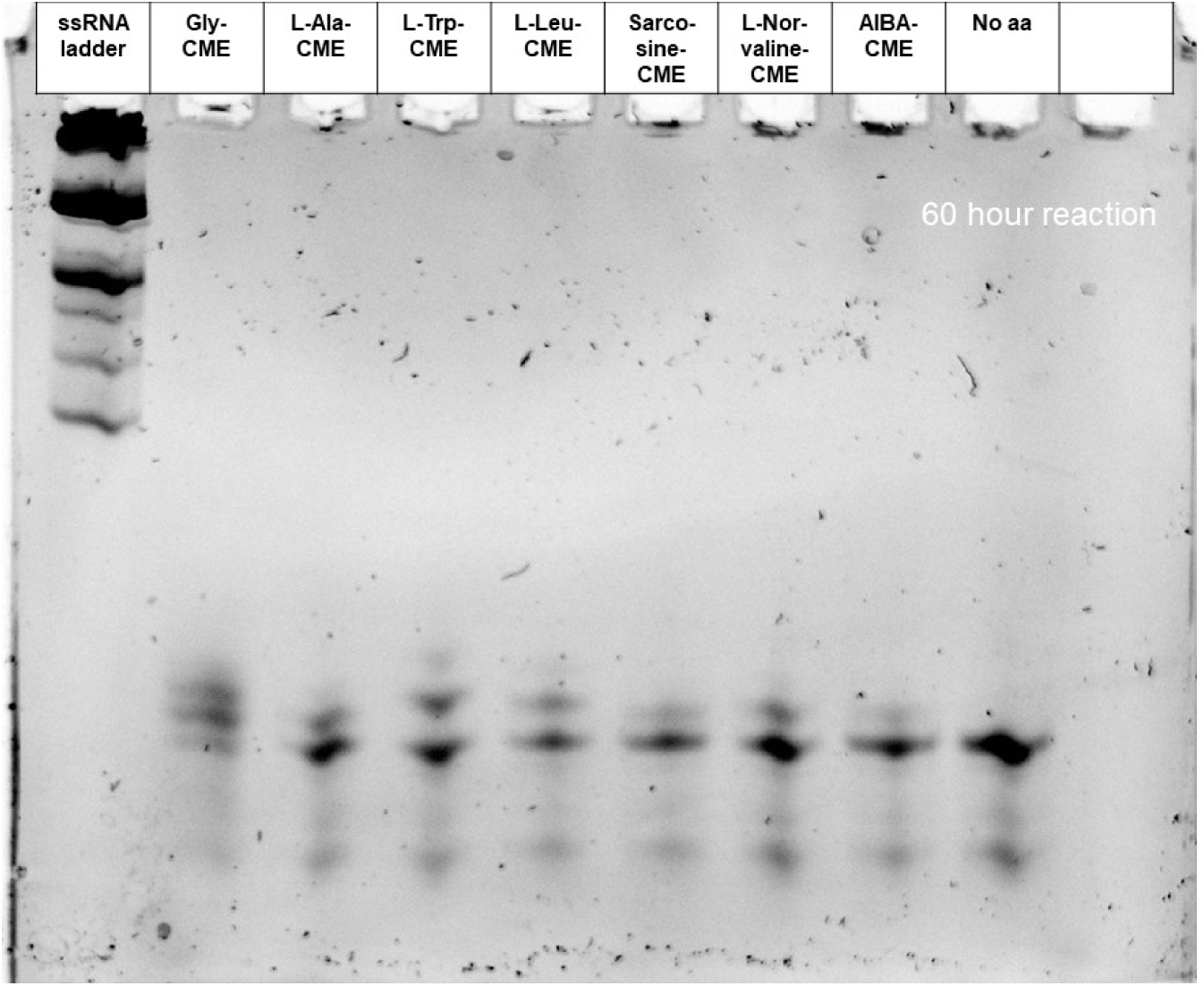
Gel of eutectic nonenzymatic acylation reactions with a variety of chemically disparate amino acids. 60 hour reaction at pH of 7.5.

**Figure S7.**
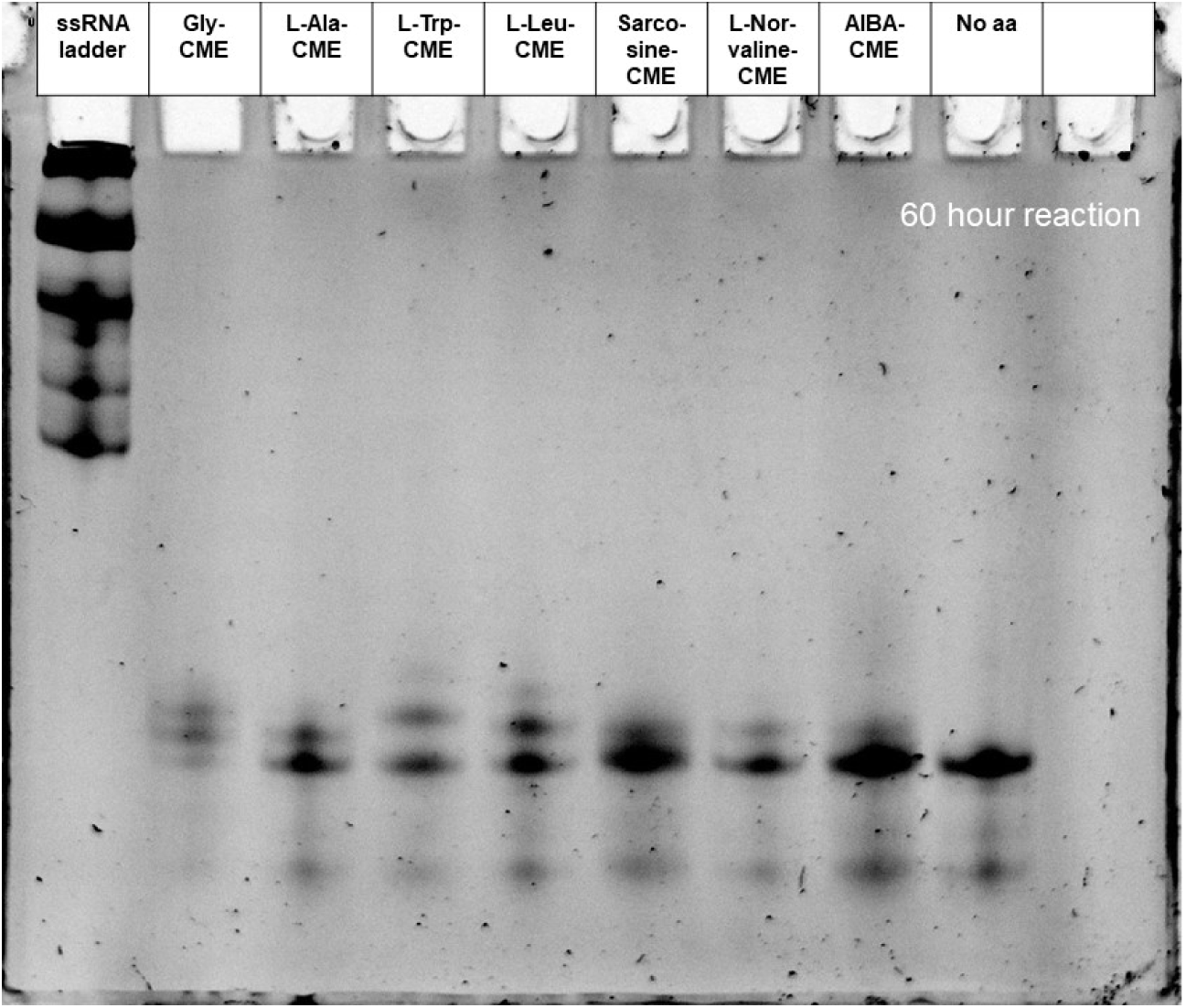
Gel of eutectic nonenzymatic acylation reactions with a variety of chemically disparate amino acids. 60 hour reaction at pH of 7.5.

**Figure S8.**
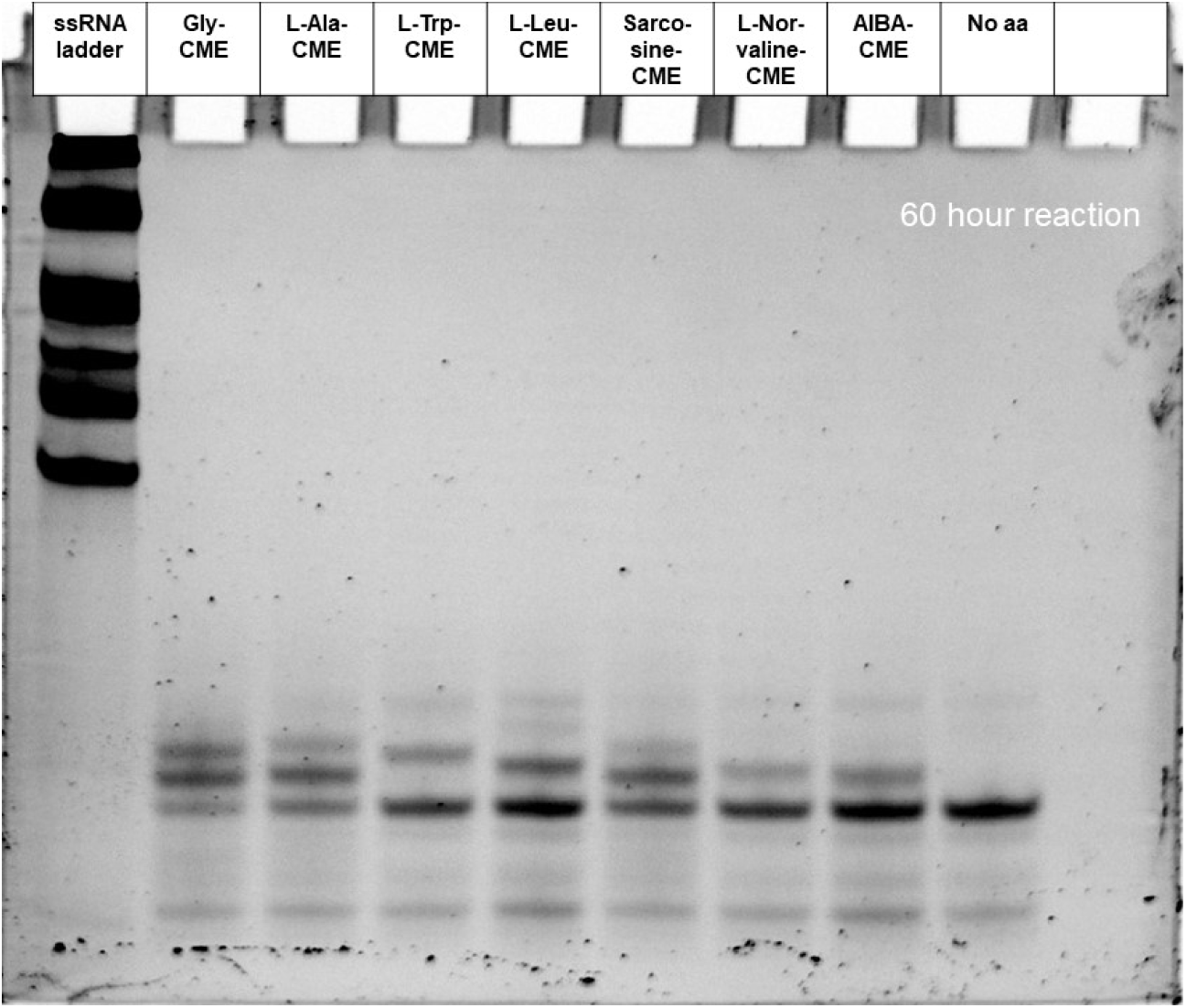
Gel of eutectic nonenzymatic acylation reactions with a variety of chemically disparate amino acids. 60 hour reaction at pH of 7.5.

**Figure S9.**
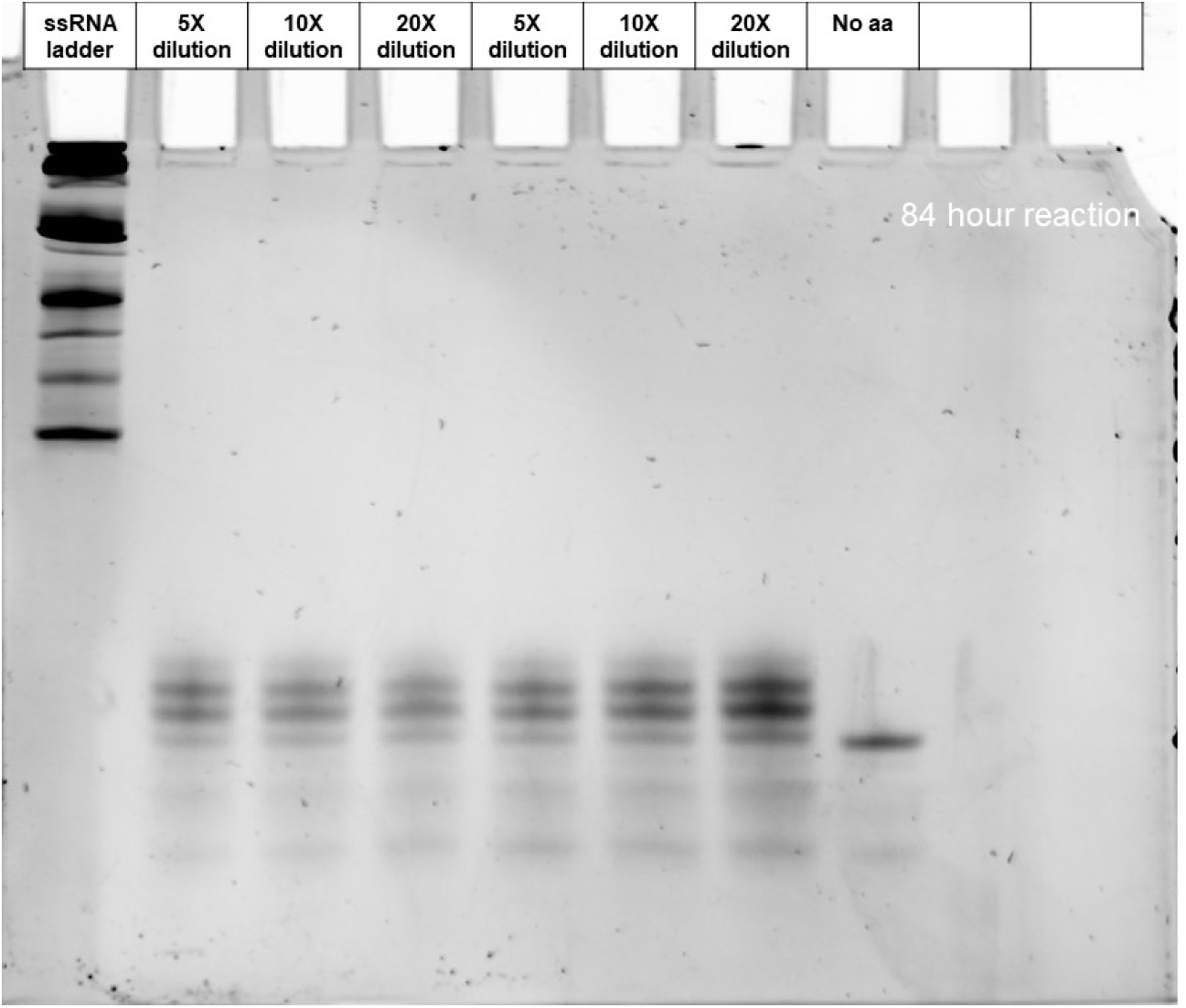
Gel of eutectic nonenzymatic acylation with 5-20X dilutions in volume before initiating the charging reaction. Glycine-CME was used for the 84 hour reaction at pH of 7.5.

**Figure S10.**
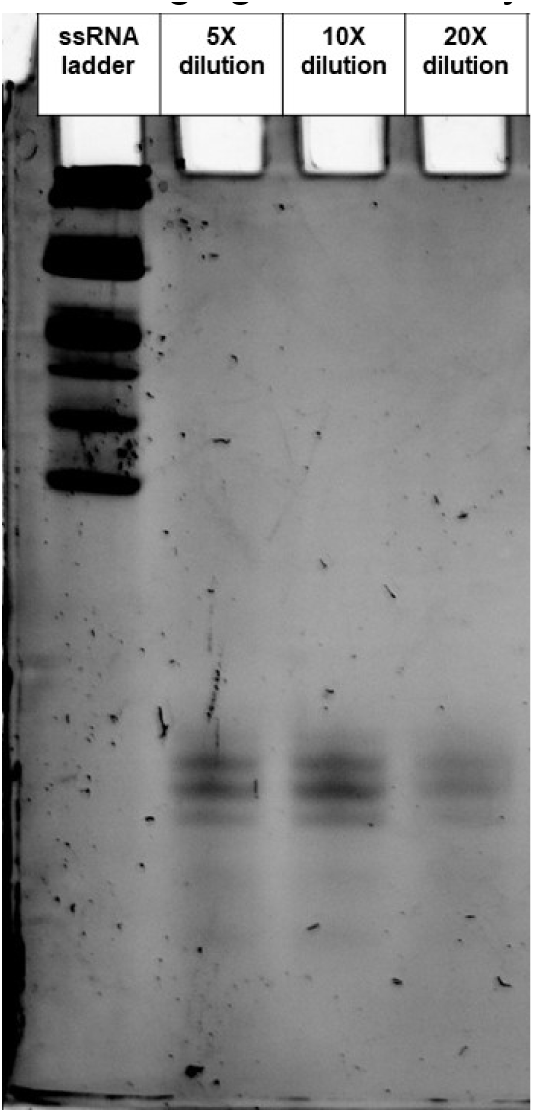
Gel of eutectic nonenzymatic acylation with 5-20X dilutions in volume before initiating the charging reaction. Glycine-CME was used for the 84 hour reaction at pH of 7.5.

**Figure S11.**
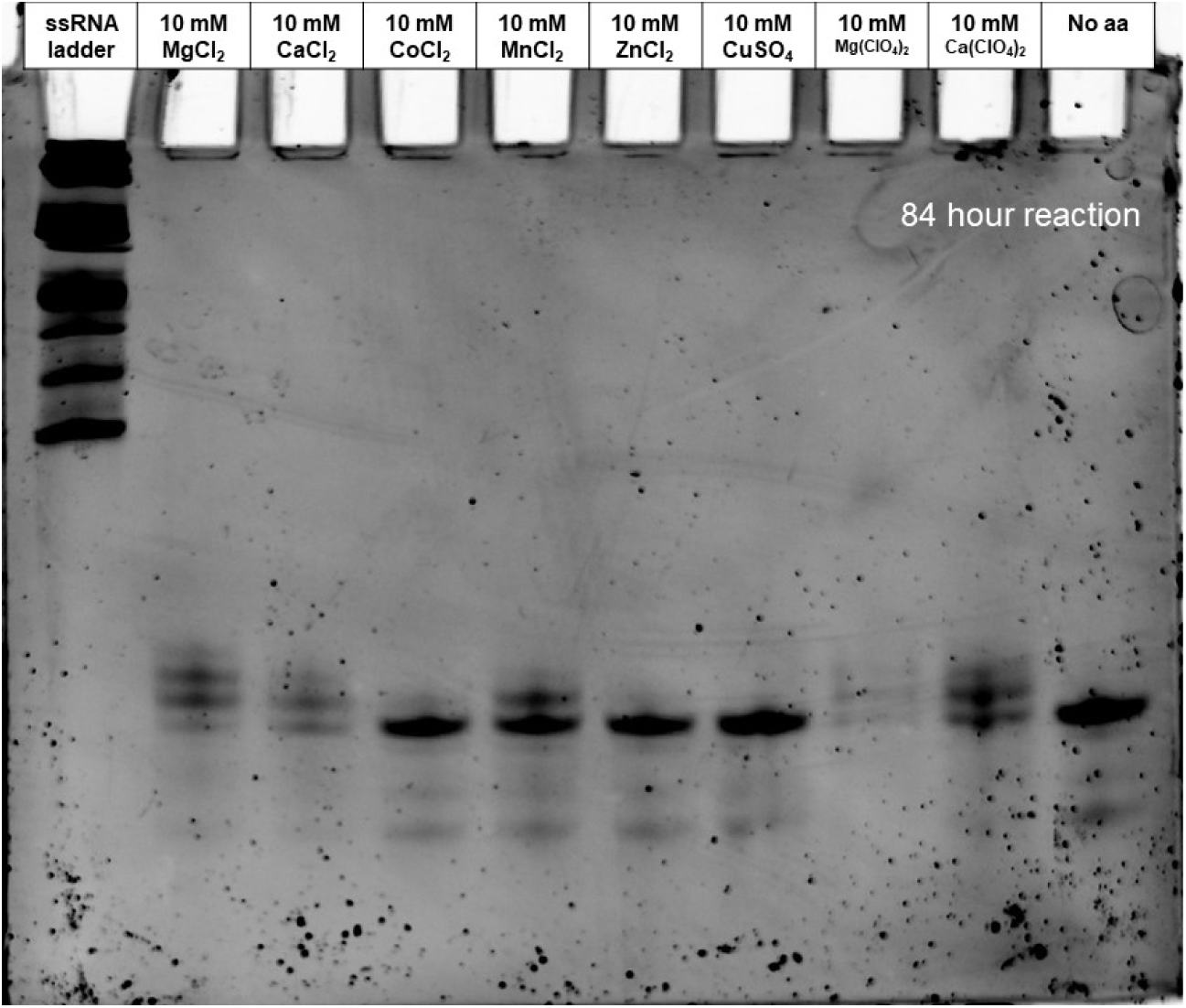
Gel of eutectic nonenzymatic acylation assessing the compatibility of the reaction with different divalent cations. All reactions used Gly-CME and were incubated for 84 hours.

**Figure S12.**
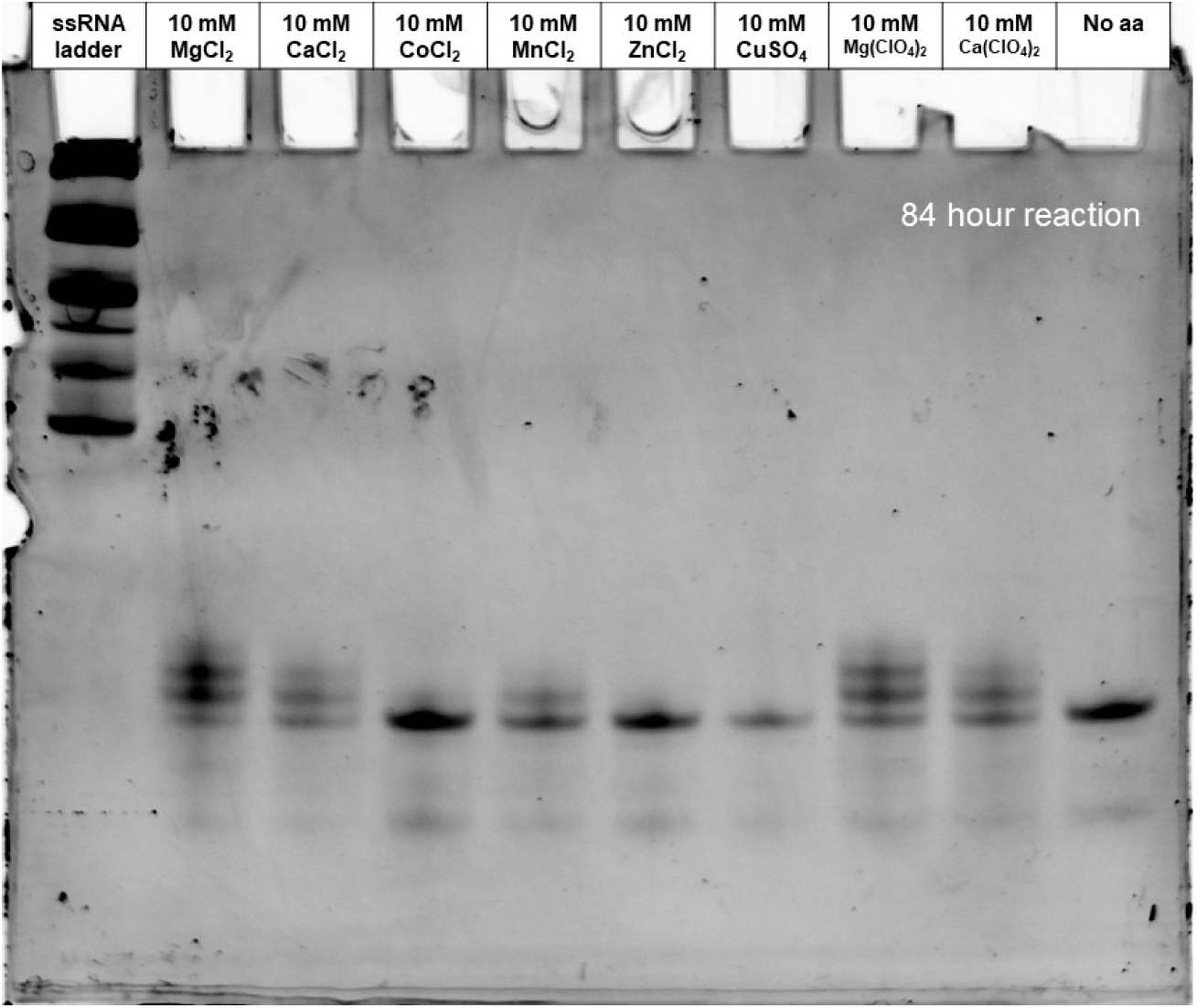
Gel of eutectic nonenzymatic acylation assessing the compatibility of the reaction with different divalent cations. All reactions used Gly-CME and were incubated for 84 hours.

**Figure S13.**
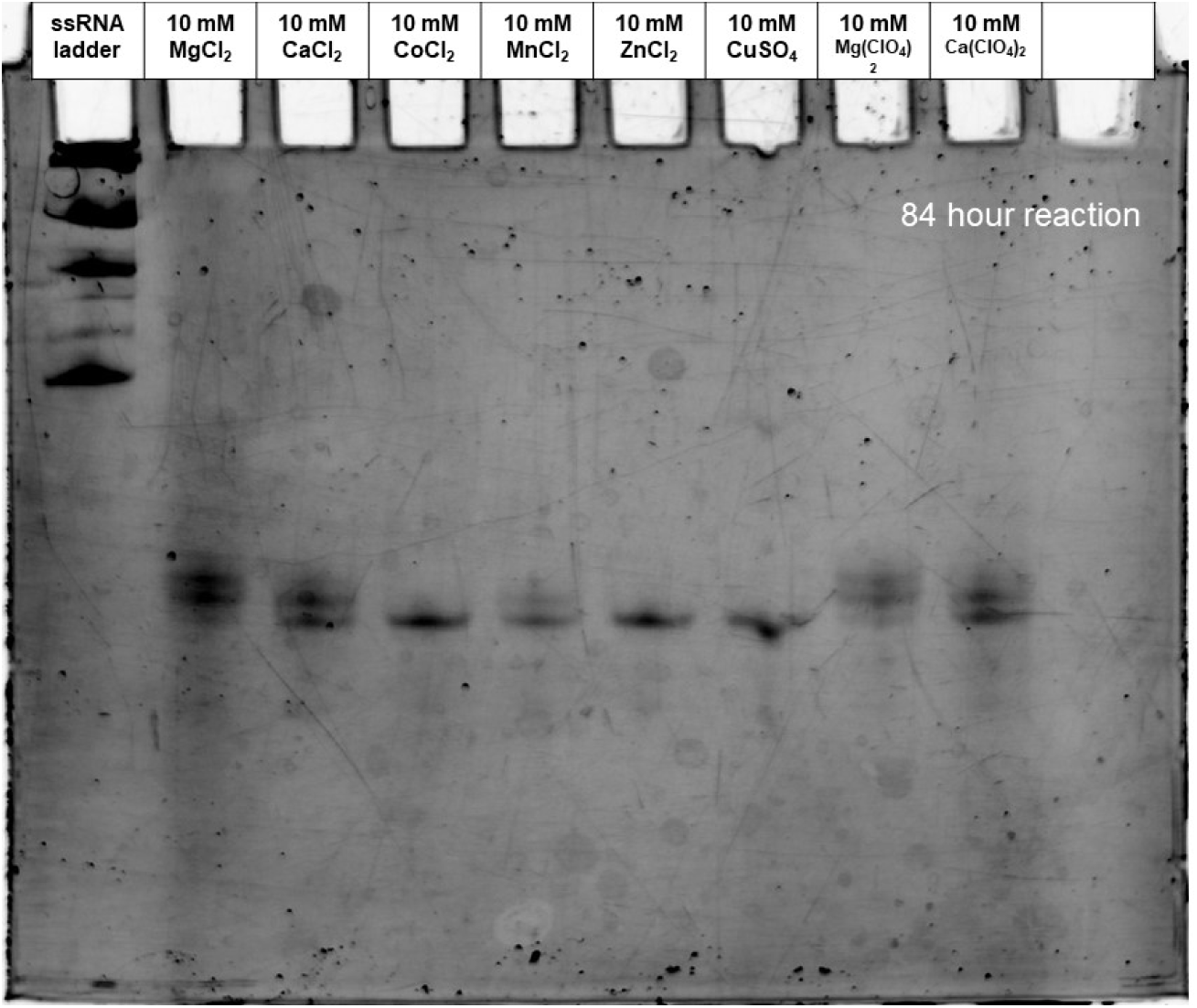
Gel of eutectic nonenzymatic acylation assessing the compatibility of the reaction with different divalent cations. All reactions used Gly-CME and were incubated for 84 hours.

**Figure S14.**
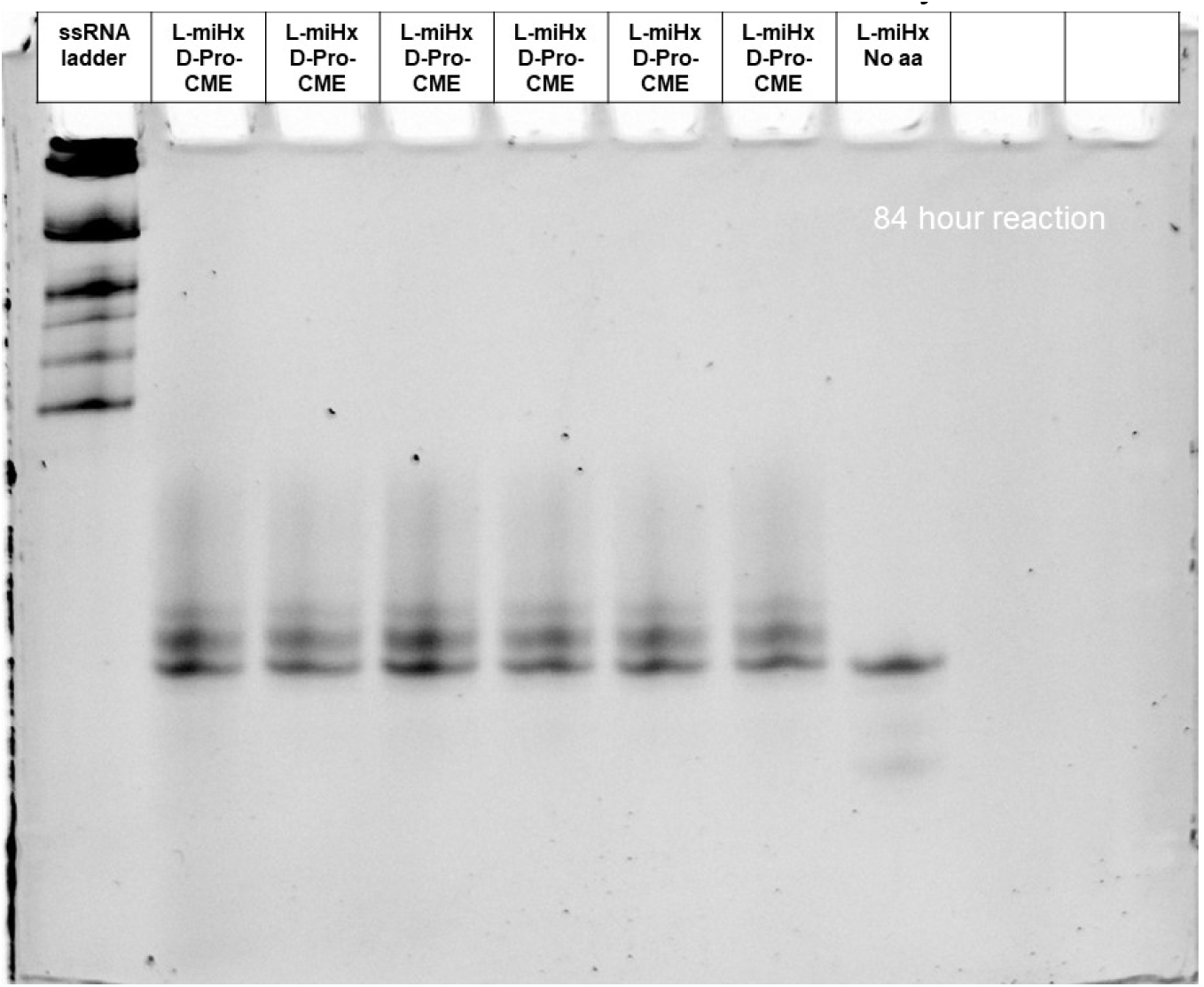
Gel of eutectic nonenzymatic acylation using L-miHx RNA to charge with D-Pro-CME for 84 hours at pH 7.5.

**Figure S15.**
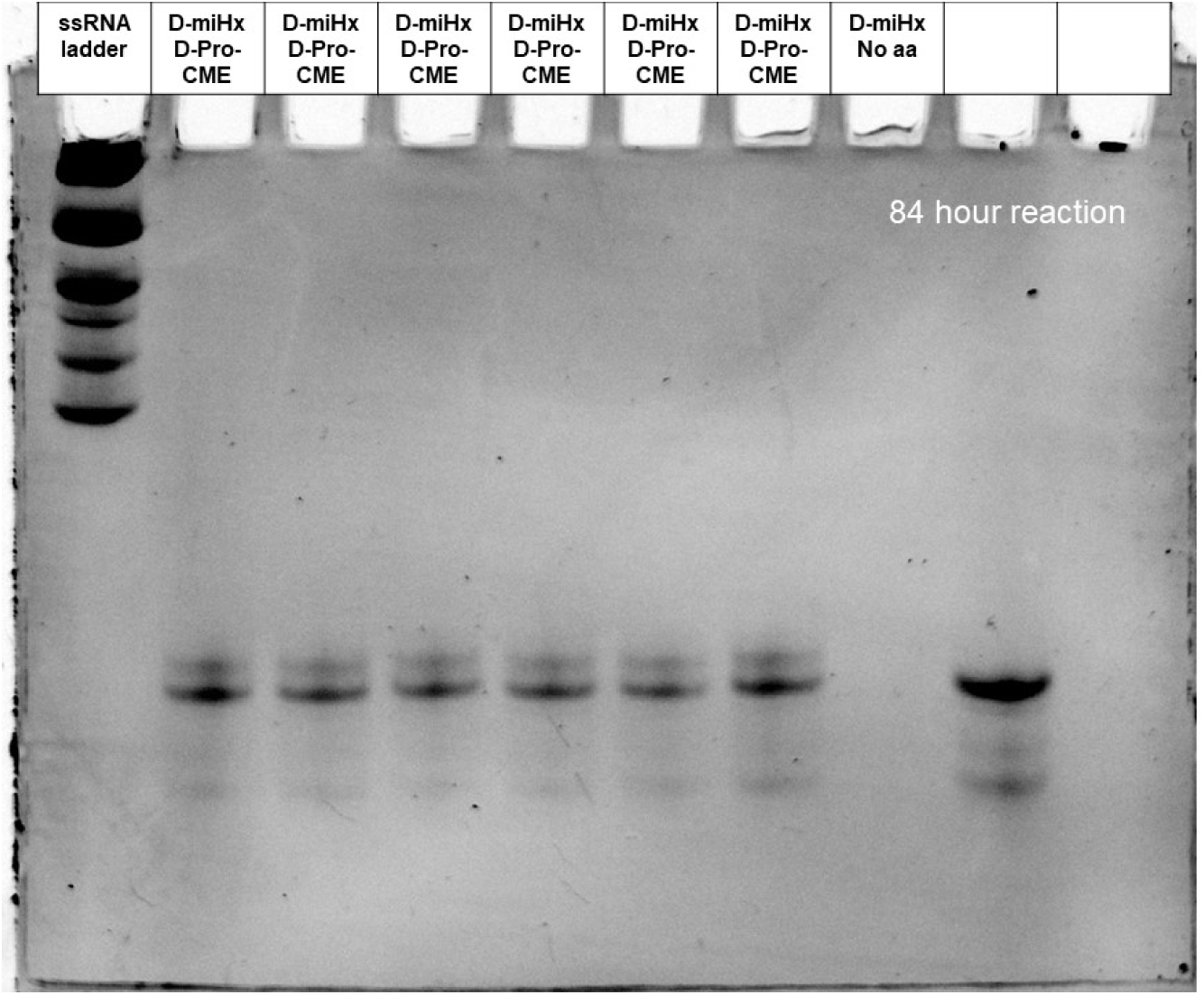
Gel of eutectic nonenzymatic acylation using D-miHx RNA to charge with D-Pro-CME for 84 hours at pH 7.5.

**Figure S16.**
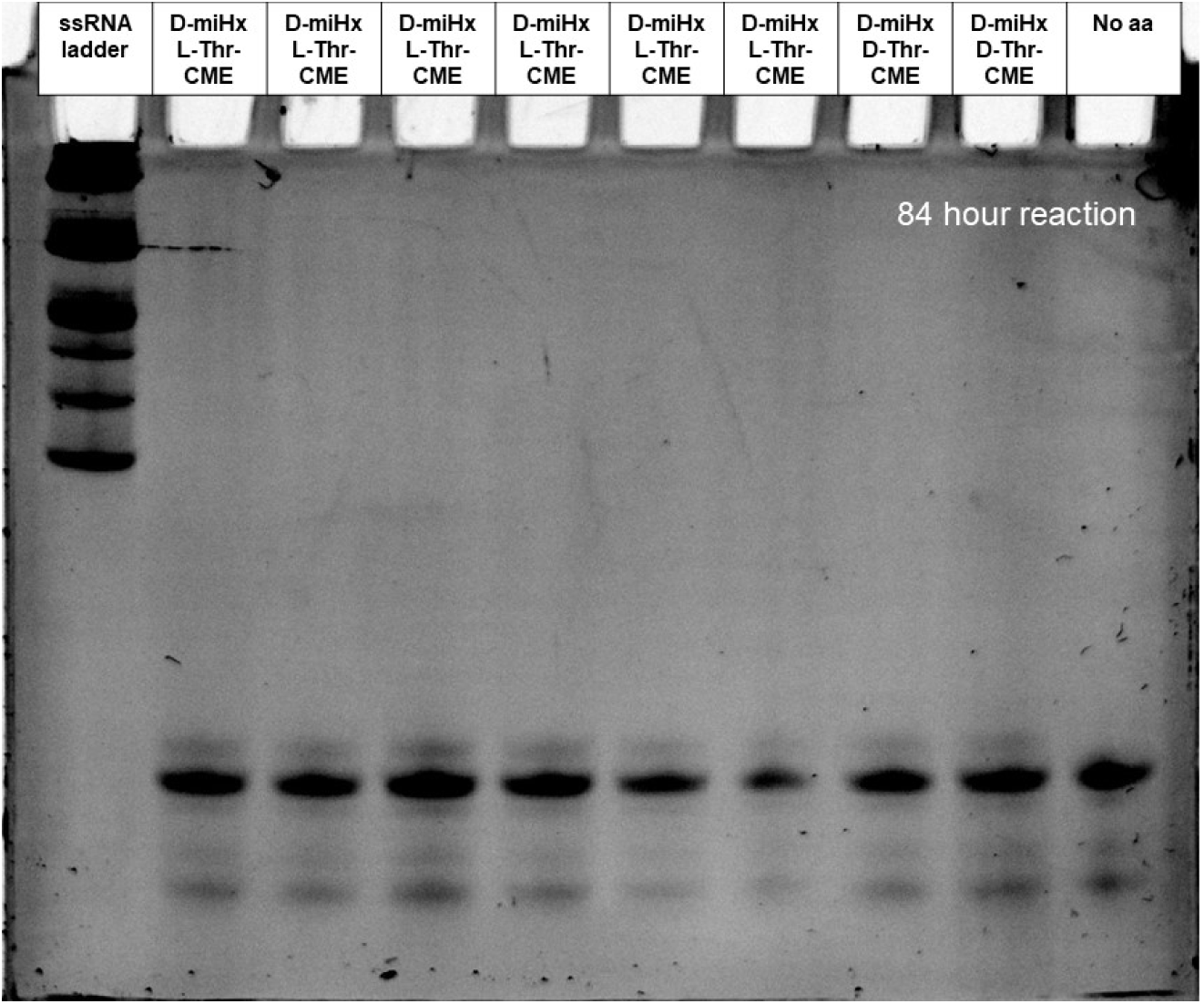
Gel of eutectic nonenzymatic acylation using D-miHx RNA to charge with L-Thr-CME or D-Thr-CME for 84 hours at pH 7.5.

**Figure S17.**
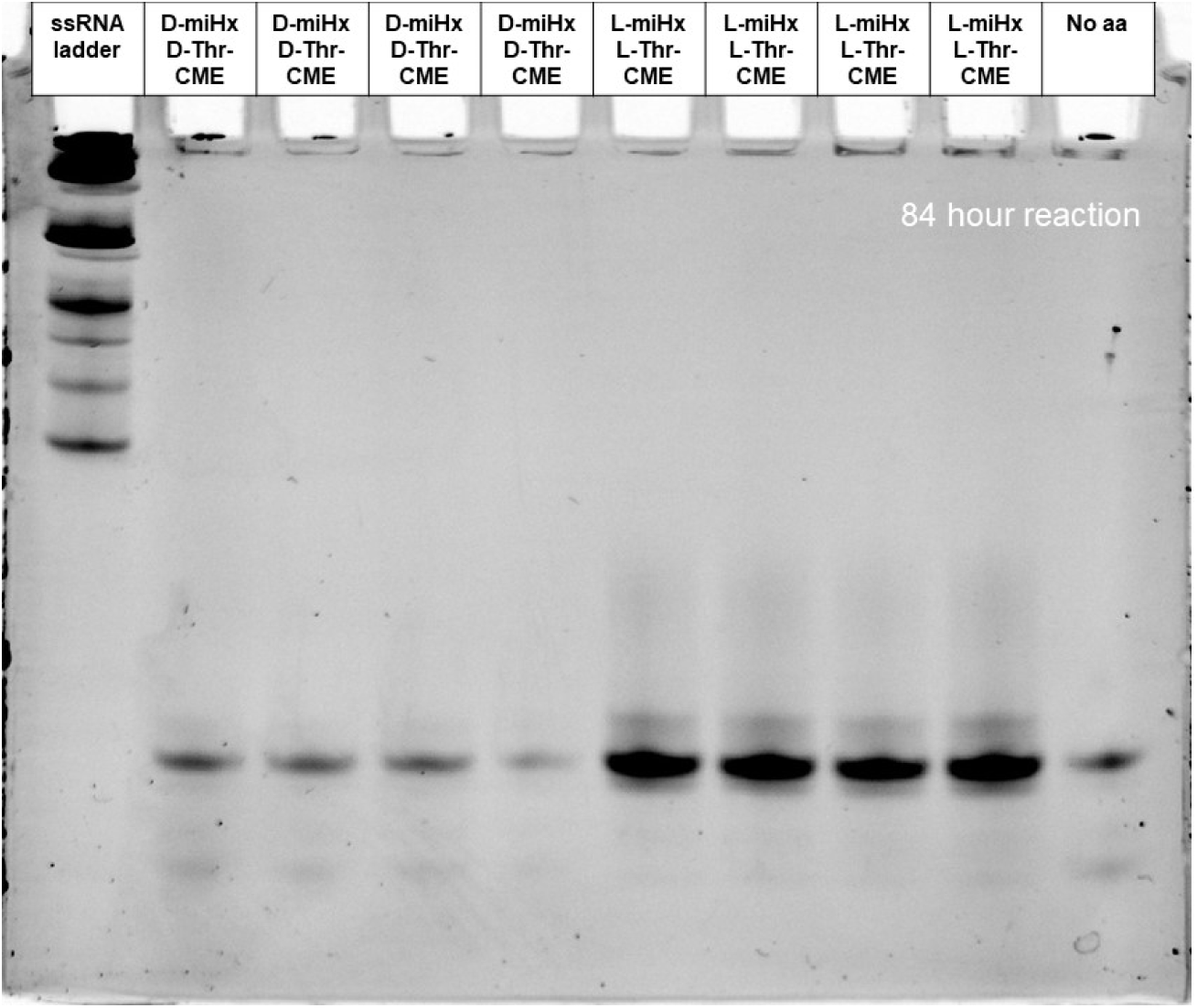
Gel of eutectic nonenzymatic acylation using D-miHx RNA or L-miHx RNA to charge with L-Thr-CME or D-Thr-CME for 84 hours at pH 7.5.

**Figure S18.**
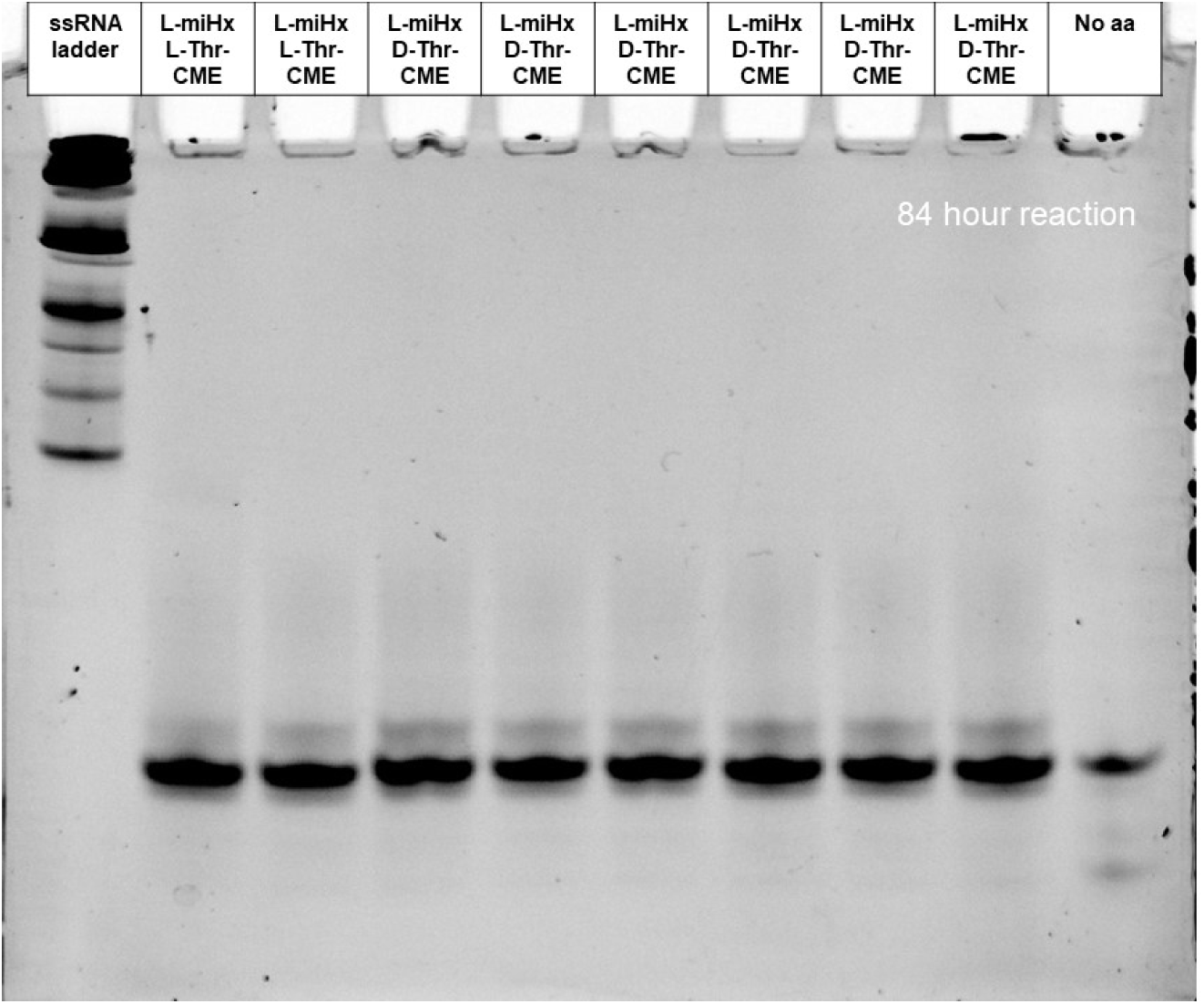
Gel of eutectic nonenzymatic acylation using L-miHx RNA to charge with L-Thr-CME or D-Thr-CME for 84 hours at pH 7.5.

**Figure S19.**
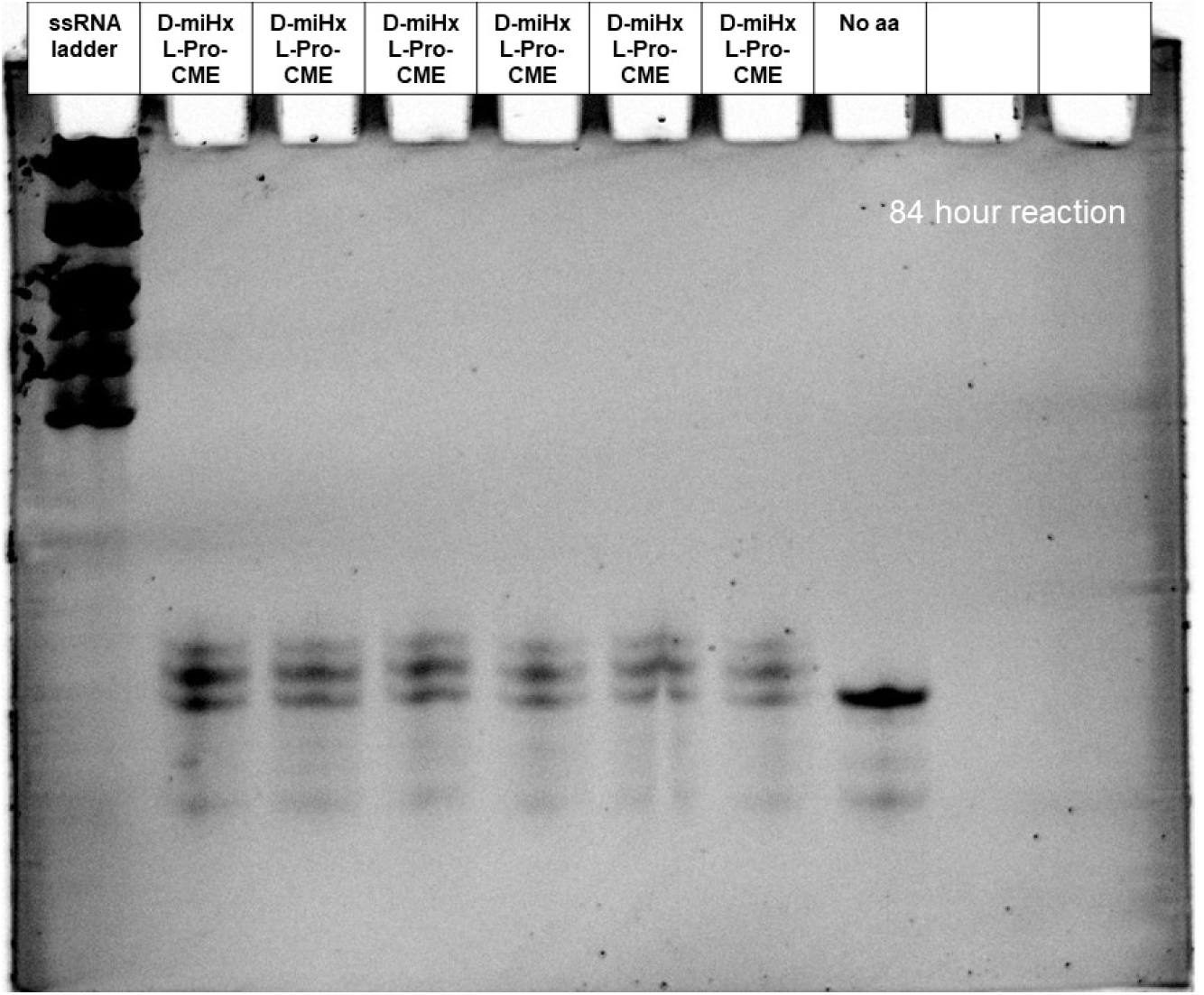
Gel of eutectic nonenzymatic acylation using D-miHx RNA to charge with L-Pro-CME for 84 hours at pH 7.5.

**Figure S20.**
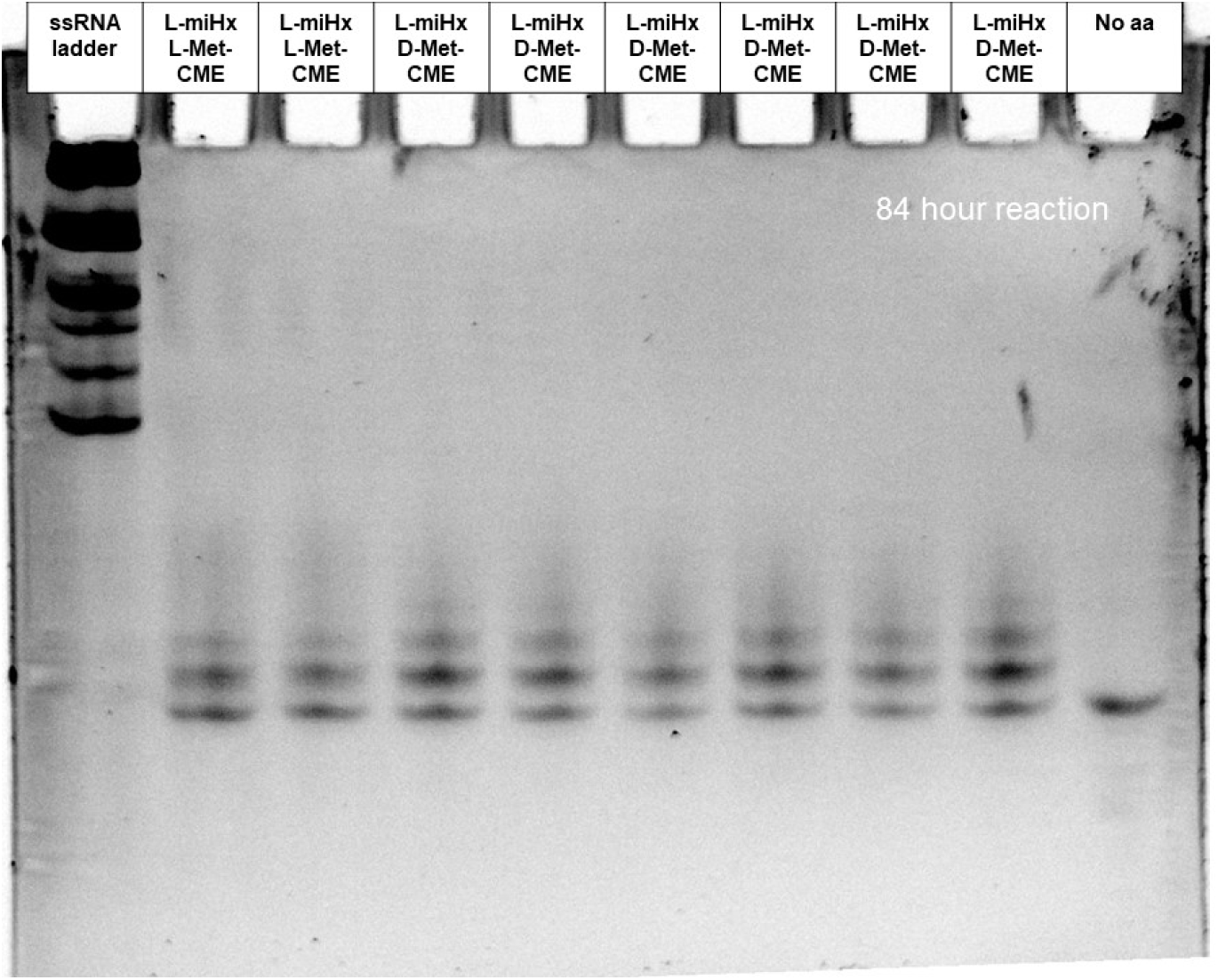
Gel of eutectic nonenzymatic acylation using L-miHx RNA to charge with L-Met-CME or D-Met-CME for 84 hours at pH 7.5.

**Figure S21.**
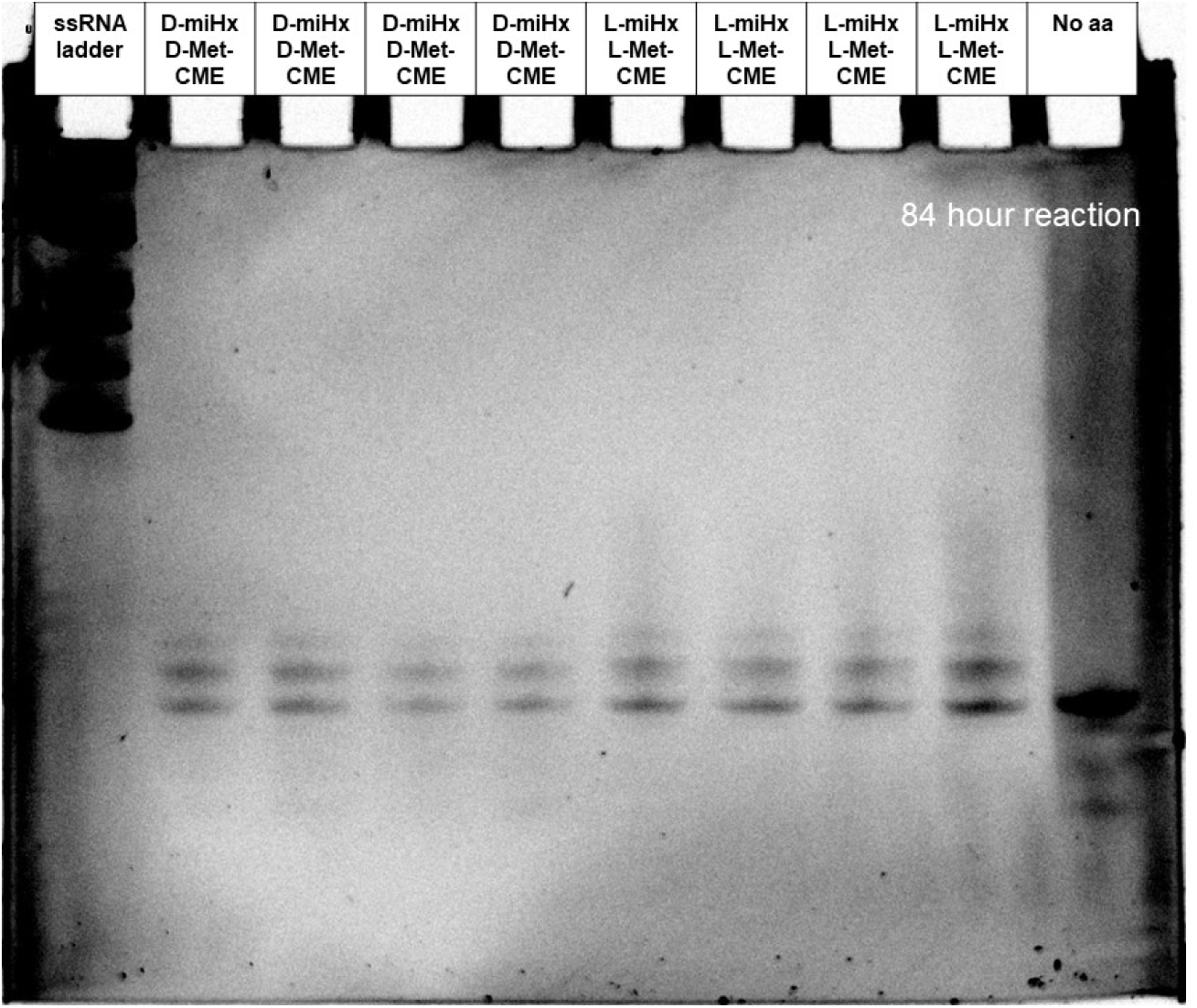
Gel of eutectic nonenzymatic acylation using D-miHx RNA or L-miHx RNA to charge with L-Met-CME or D-Met-CME for 84 hours at pH 7.5.

**Figure S22.**
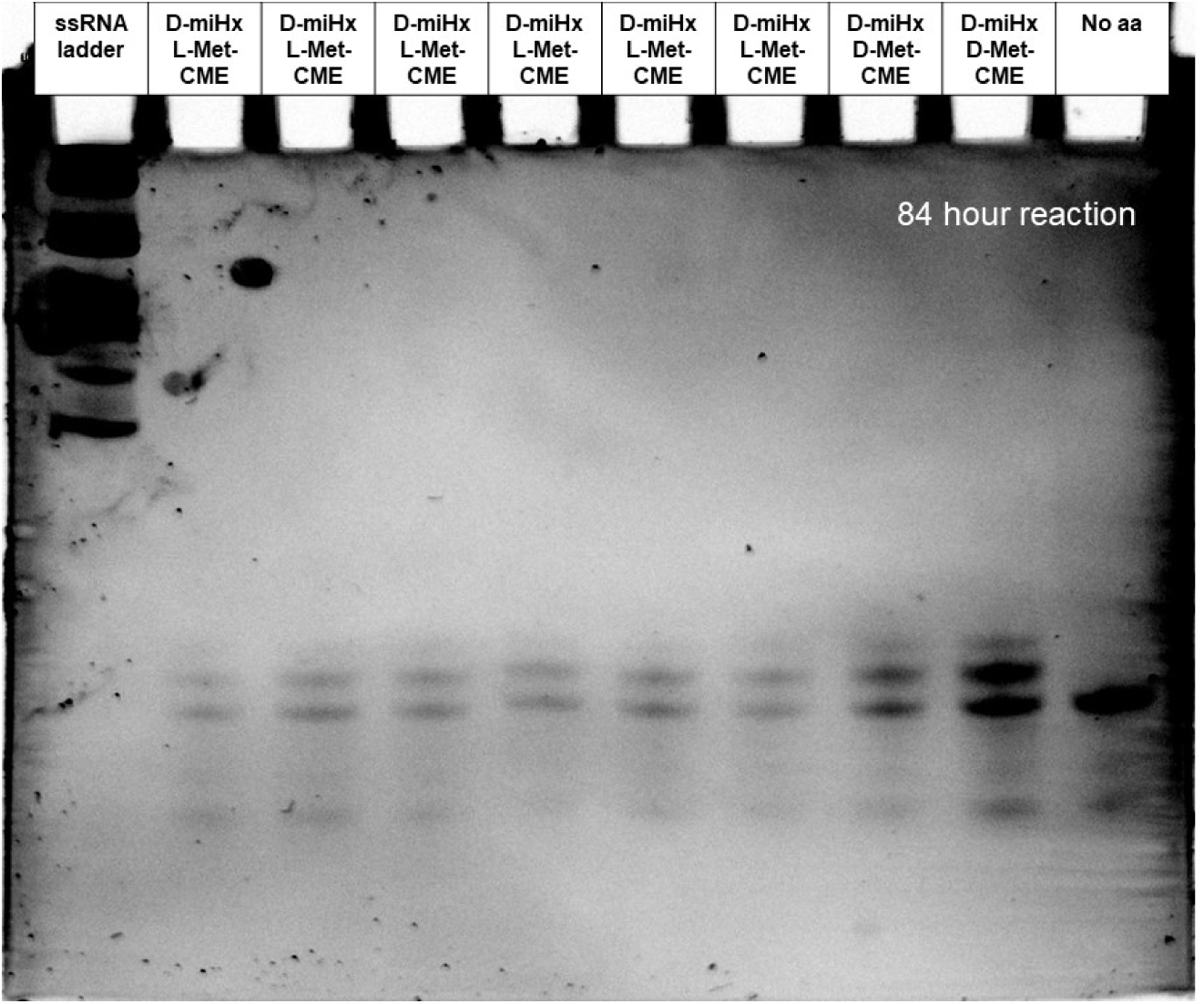
Gel of eutectic nonenzymatic acylation using D-miHx RNA to charge with L-Met-CME or D-Met-CME for 84 hours at pH 7.5.

**Figure S23.**
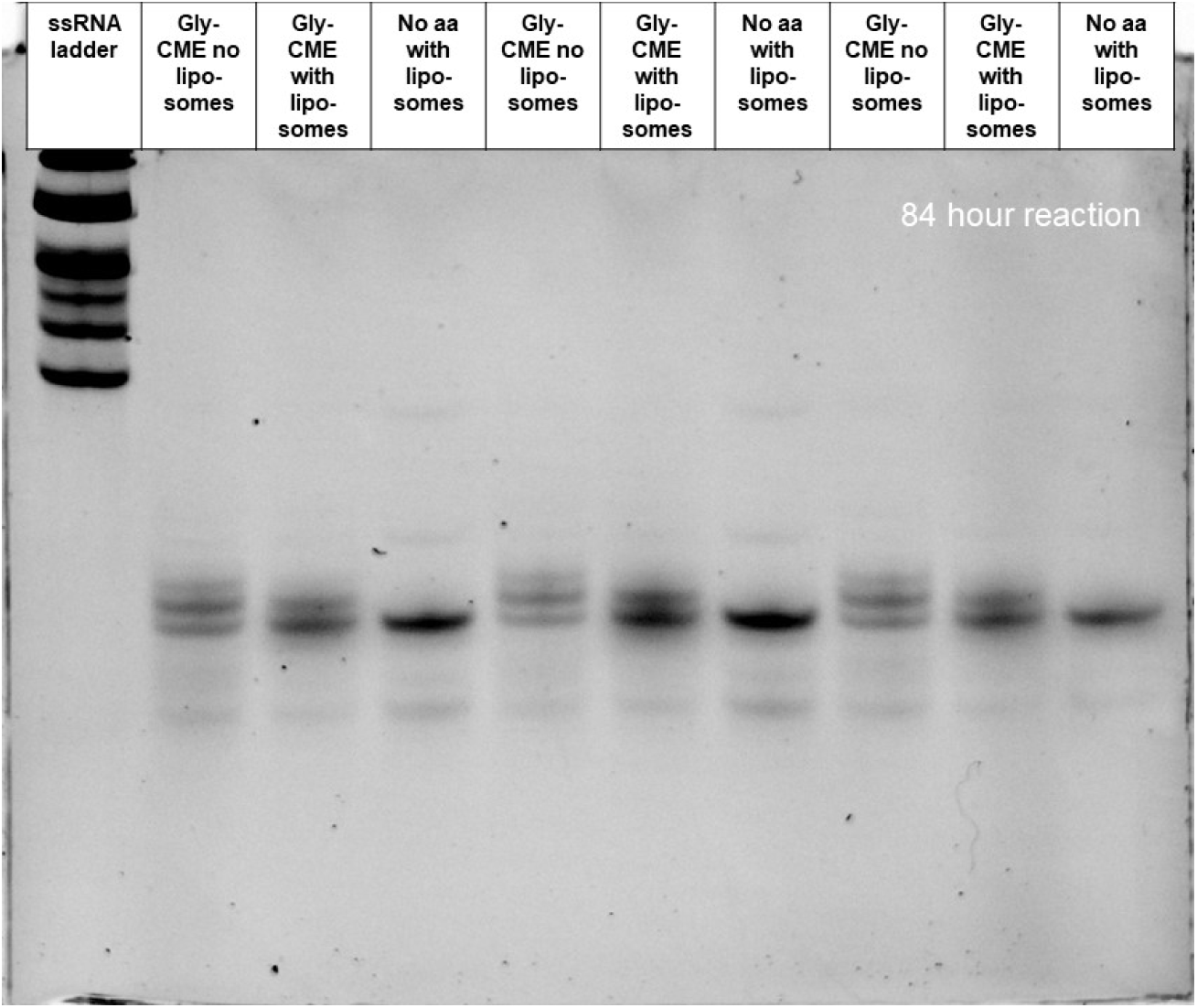
Gel of eutectic nonenzymatic acylation in the presence of oleic acid liposomes and 40 mM citrate. The reaction as performed with no liposomes + glycine-CME, with liposomes + glycine-CME, and with liposomes but without glycine-CME for 84 hours at pH 7.5.

**Figure S24.**
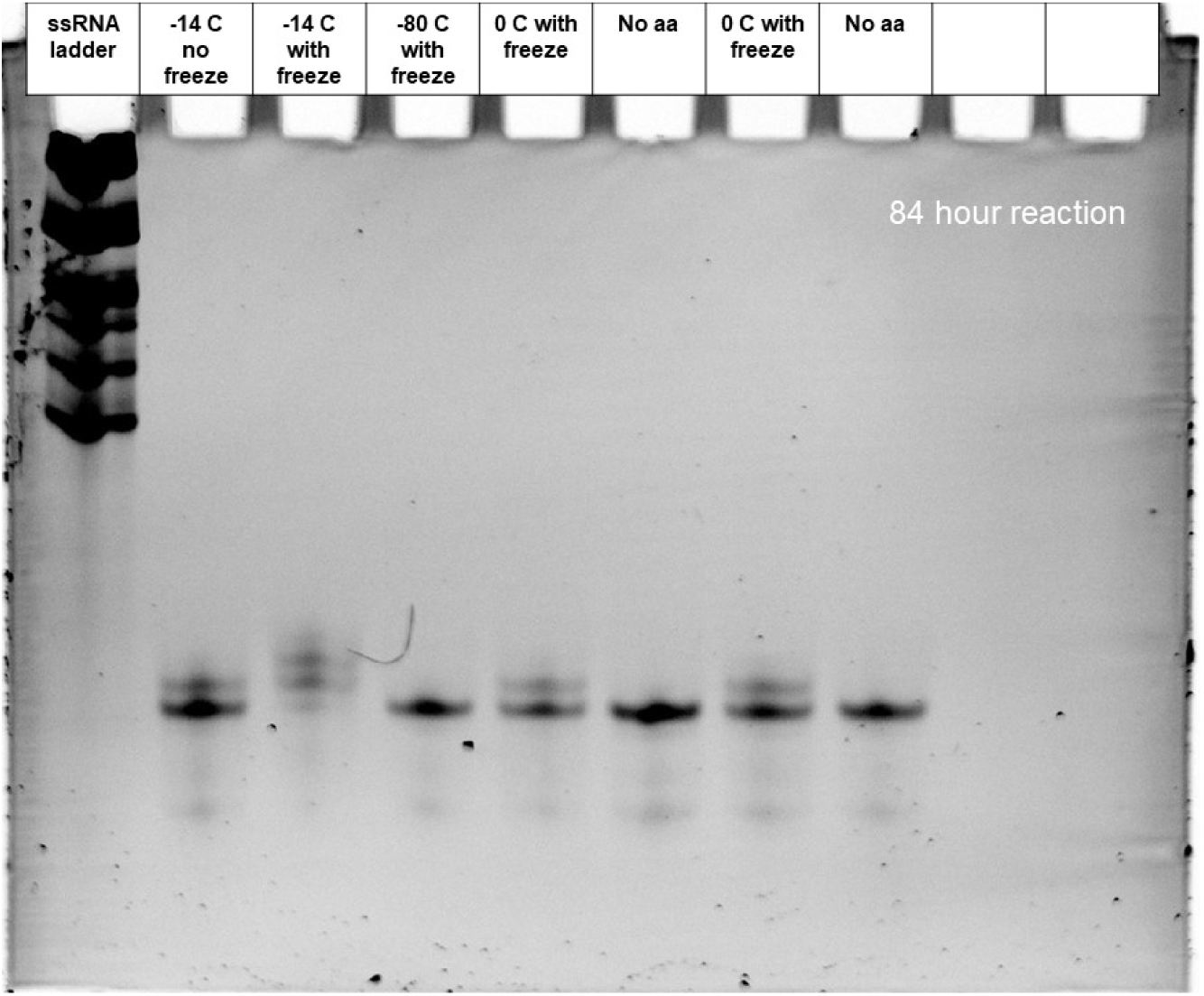
Gel of nonenzymatic acylation reactions charging glycine-CME testing the conditions necessary for acylation at different temperatures, eutectic phase vs liquid, never frozen reactions for 84 hours at a pH of 7.5.

**Figure S25.**
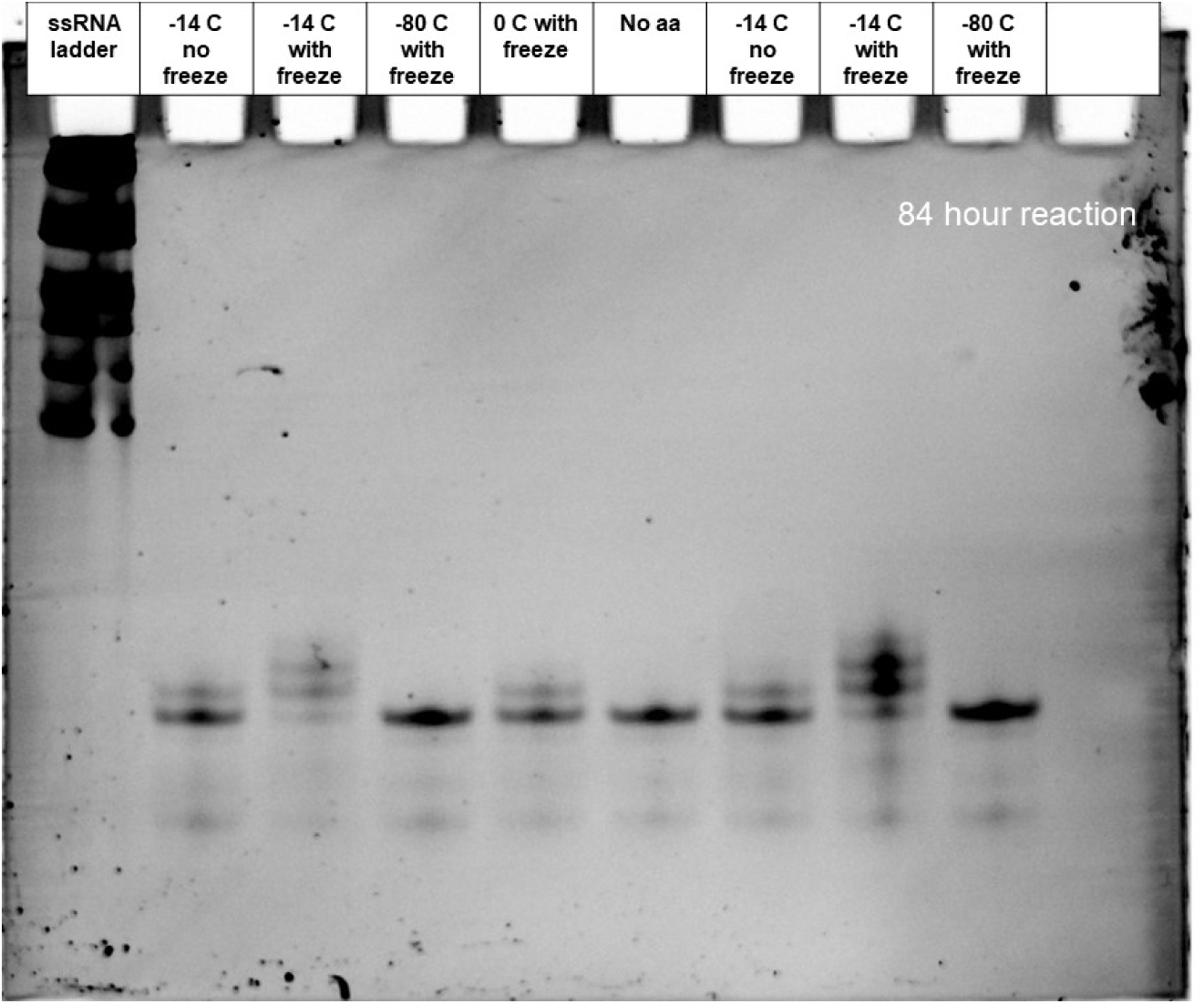
Gel of nonenzymatic acylation reactions charging glycine-CME testing the conditions necessary for acylation at different temperatures, eutectic phase vs liquid, never frozen reactions for 84 hours at a pH of 7.5.

**Figure S26.**
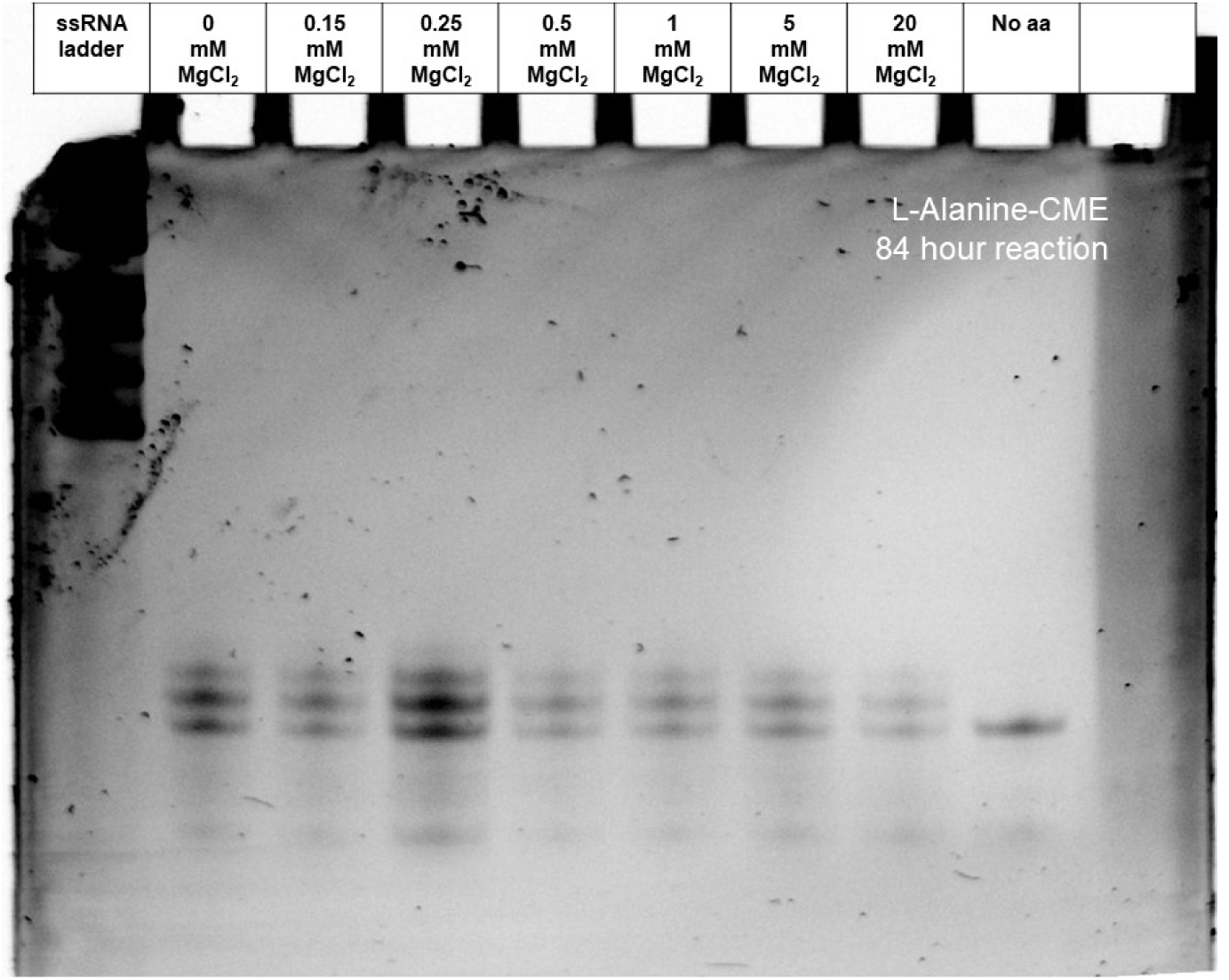
Gel of eutectic nonenzymatic acylation reactions with decreasing concentrations of MgCl_2_ without EDTA added. L-alanine-CME was charged for 84 hours at pH 7.5.

**Figure S27.**
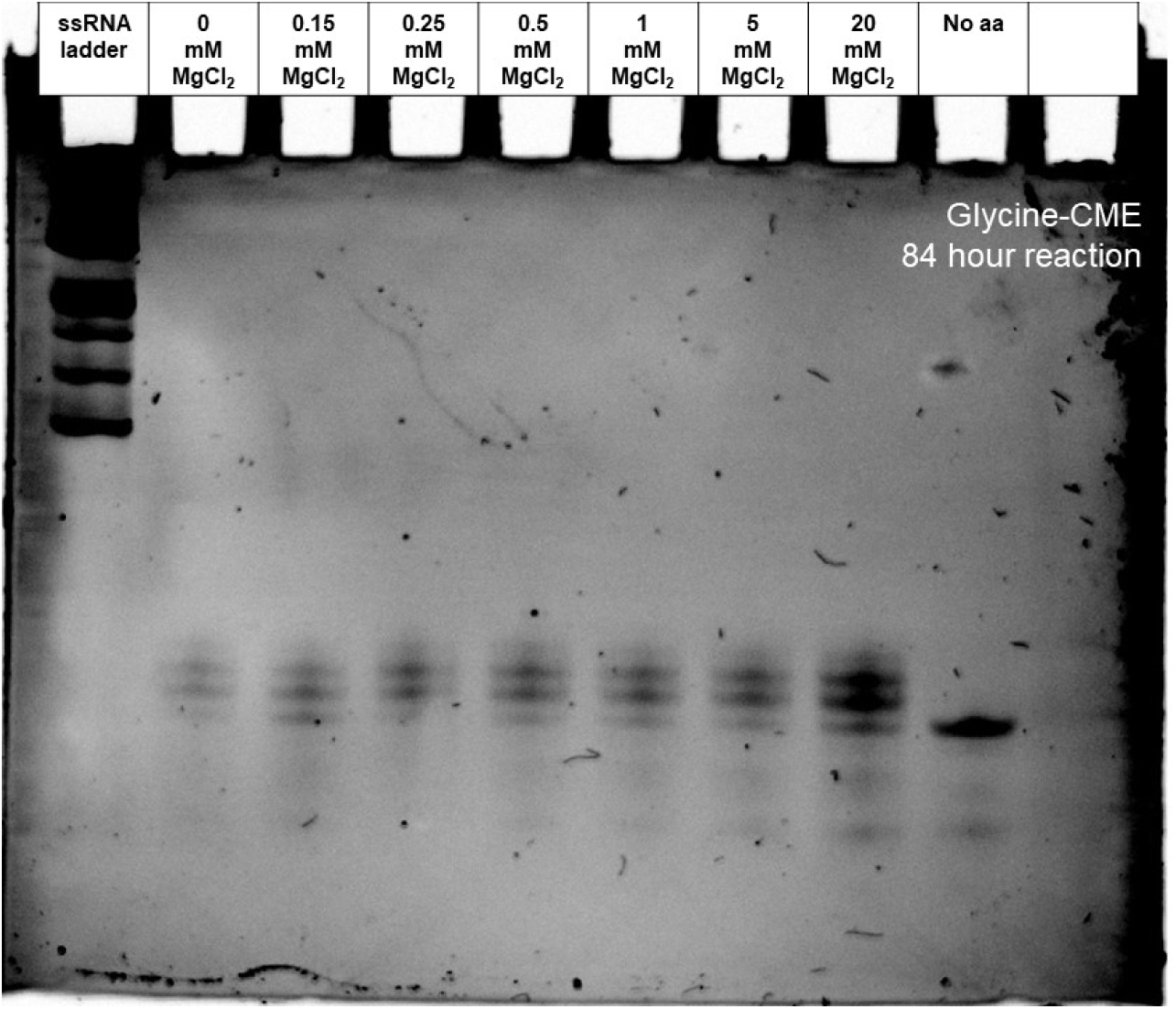
Gel of eutectic nonenzymatic acylation reactions with decreasing concentrations of MgCl_2_ without EDTA added. Glycine -CME was charged for 84 hours at pH 7.5.

**Figure S28.**
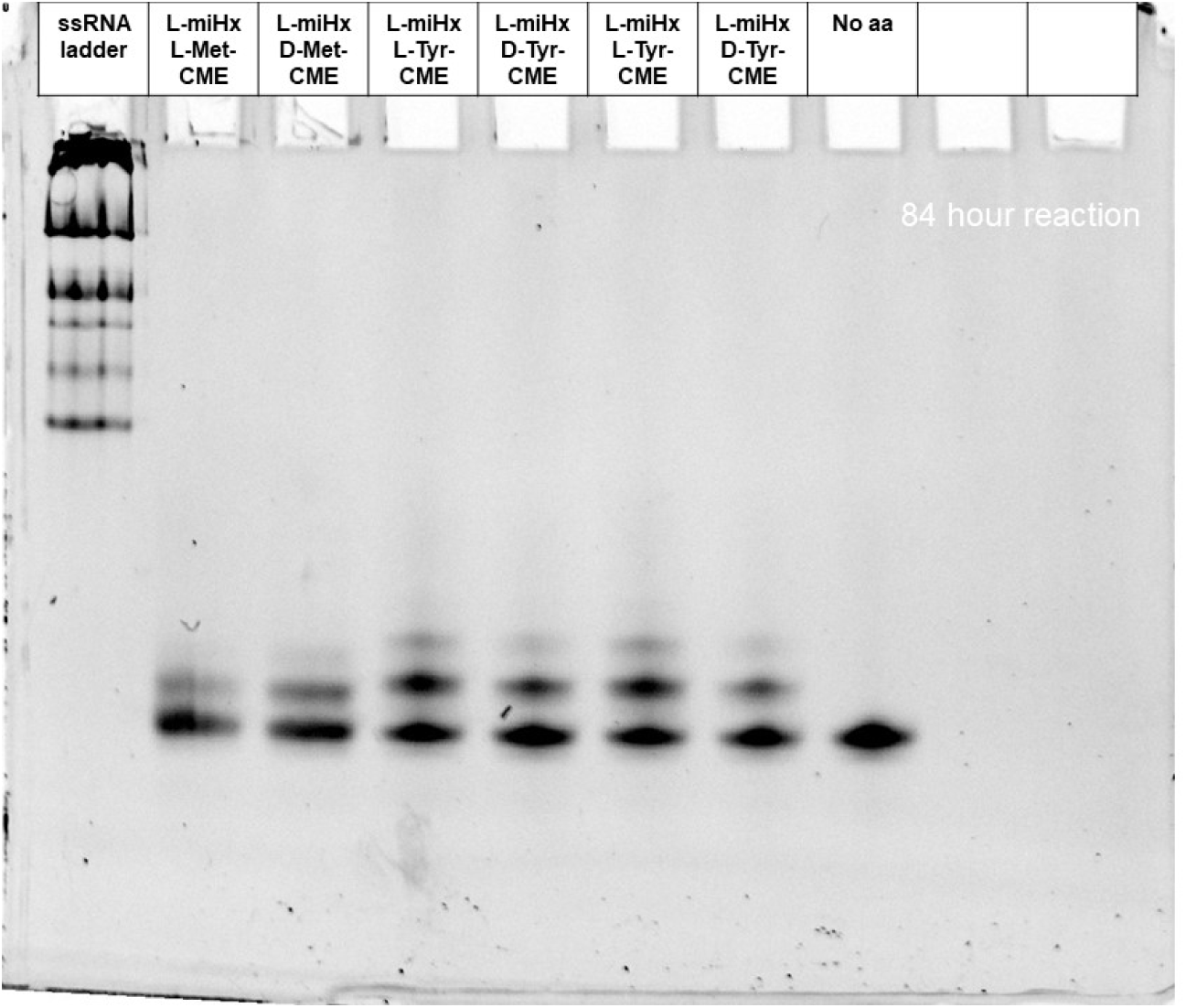
Gel of eutectic nonenzymatic acylation using L-miHx RNA to charge with L/D-Met-CME or L/D-Tyr-CME for 84 hours at pH 7.5.

**Figure S29.**
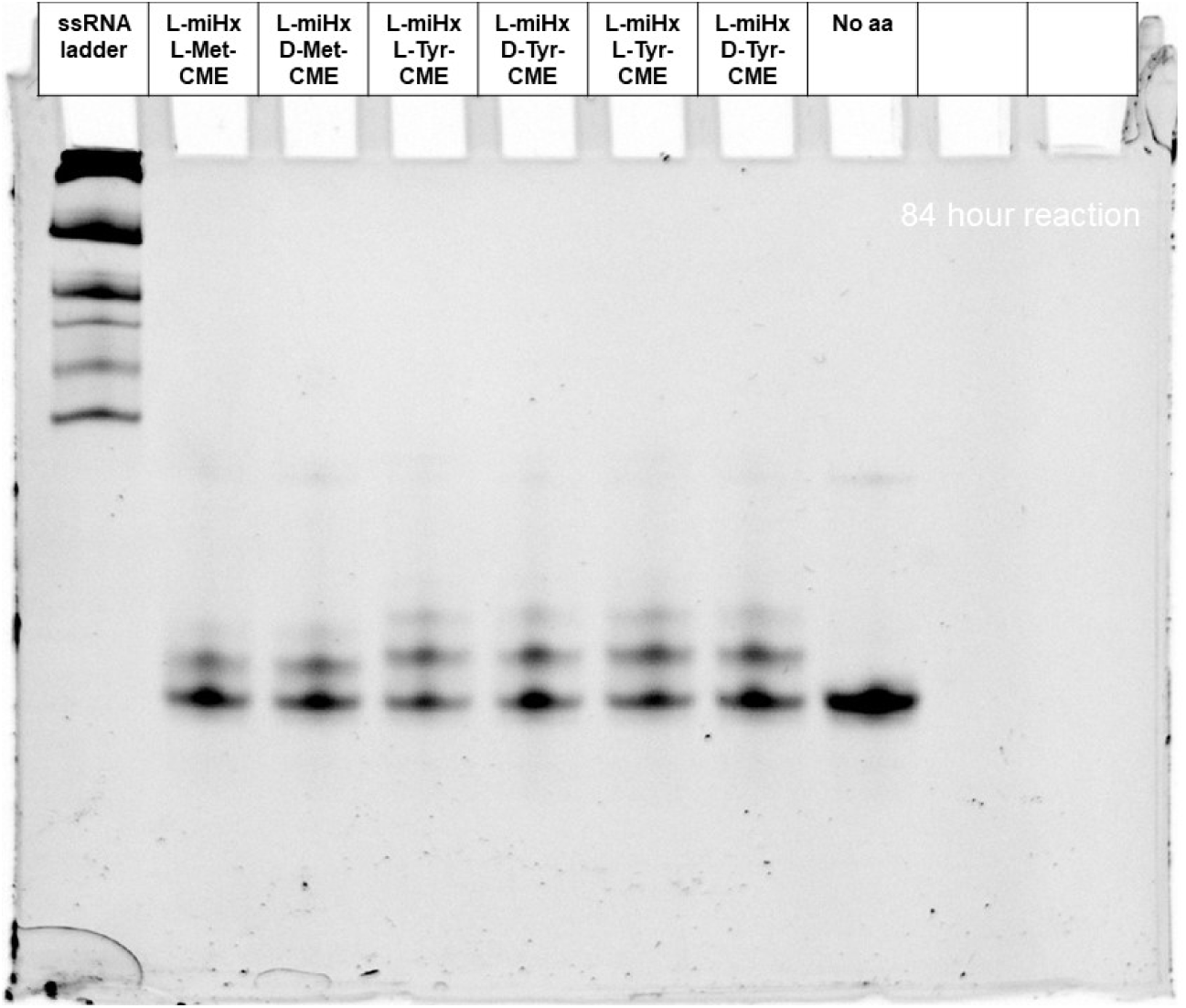
Gel of eutectic nonenzymatic acylation using L-miHx RNA to charge with L/D-Met-CME or L/D-Tyr-CME for 84 hours at pH 7.5.

**Figure S30.**
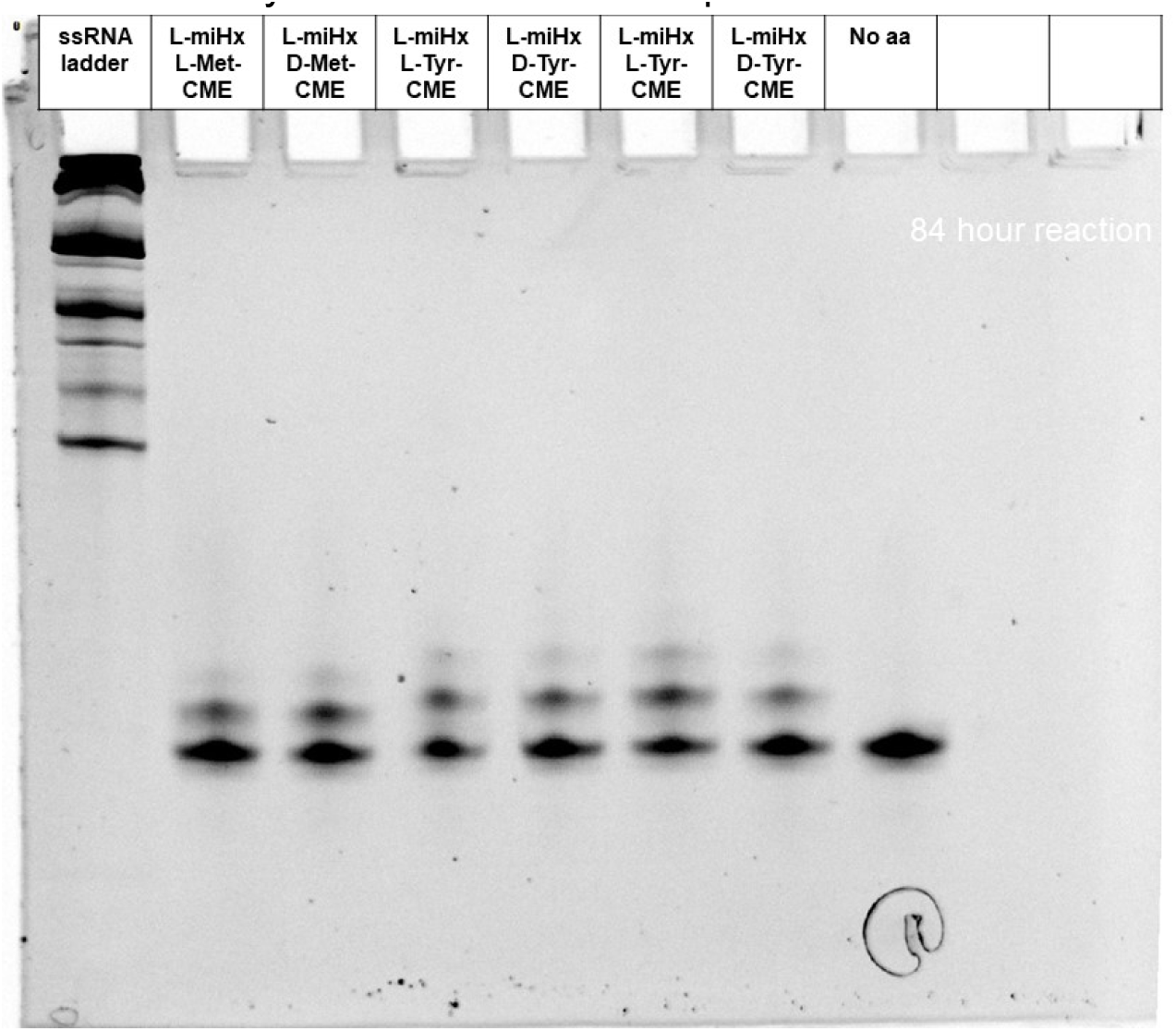
Gel of eutectic nonenzymatic acylation using L-miHx RNA to charge with L/D-Met-CME or L/D-Tyr-CME for 84 hours at pH 7.5.

**Figure S31.**
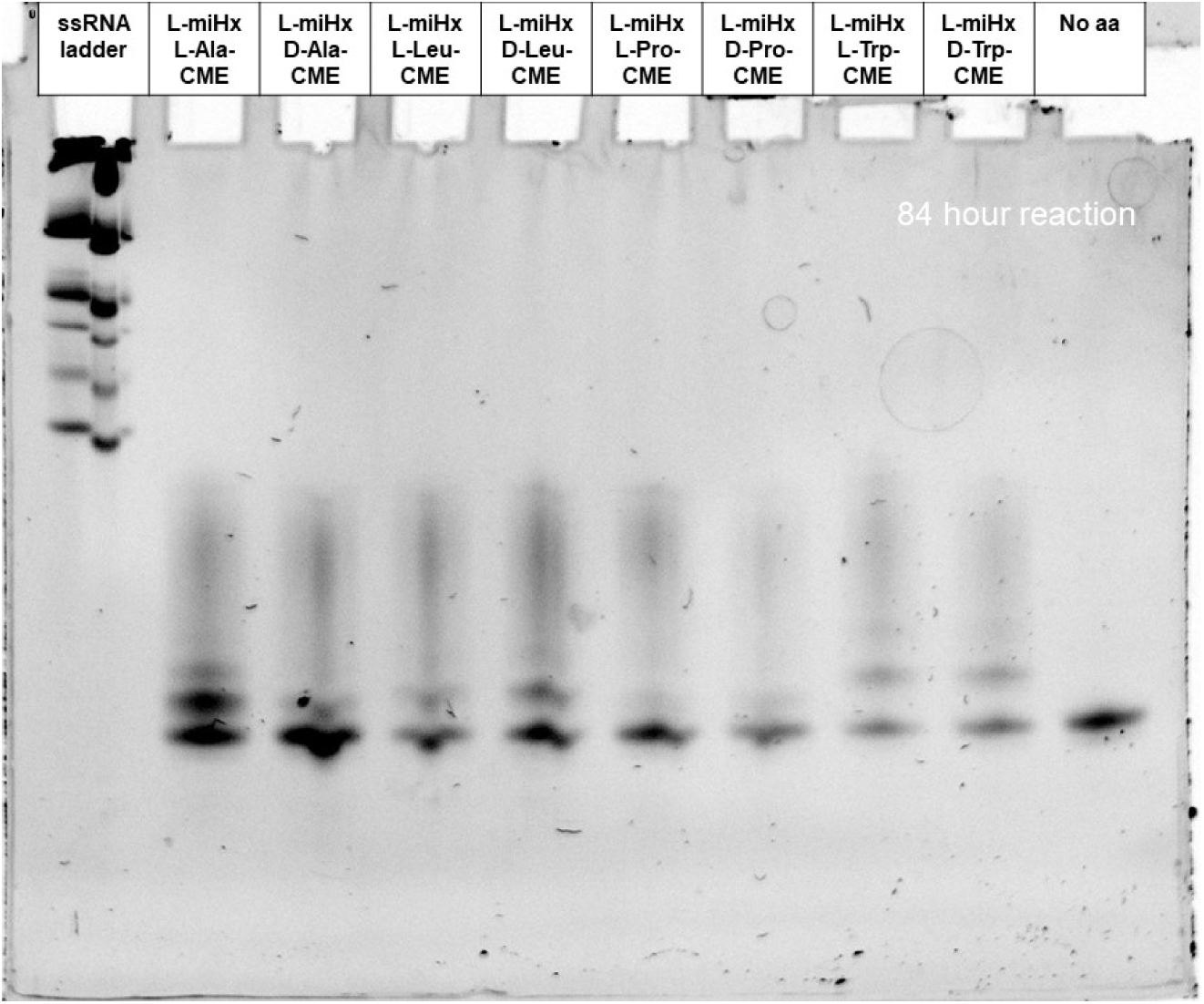
Gel of eutectic nonenzymatic acylation using L-miHx RNA to charge with L/D-Ala-CME or L/D-Leu-CME or L/D-Pro-CME or L/D-Trp-CME for 84 hours at pH 7.5.

**Figure S32.**
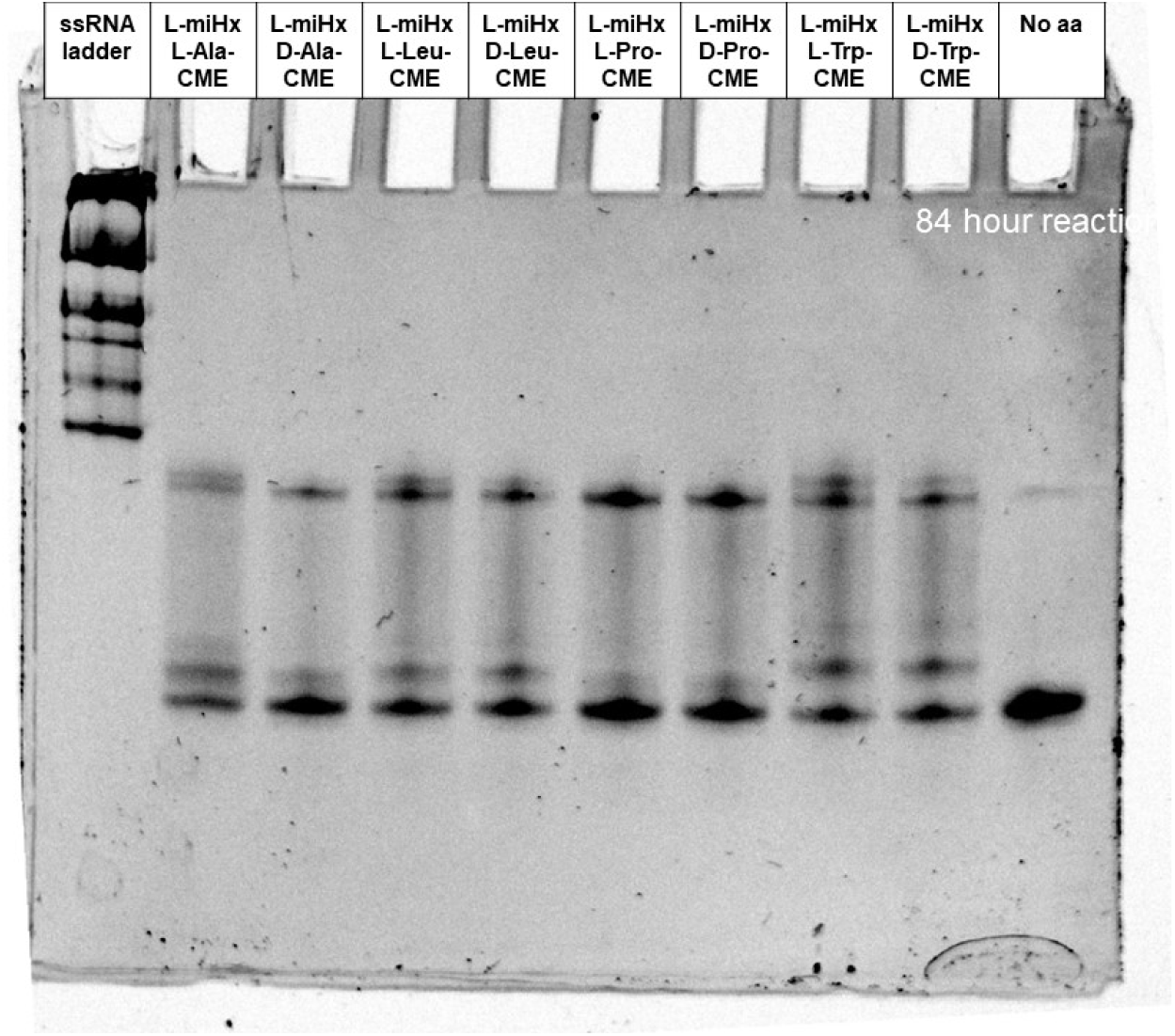
Gel of eutectic nonenzymatic acylation using L-miHx RNA to charge with L/D-Ala-CME or L/D-Leu-CME or L/D-Pro-CME or L/D-Trp-CME for 84 hours at pH 7.5.

**Figure S33.**
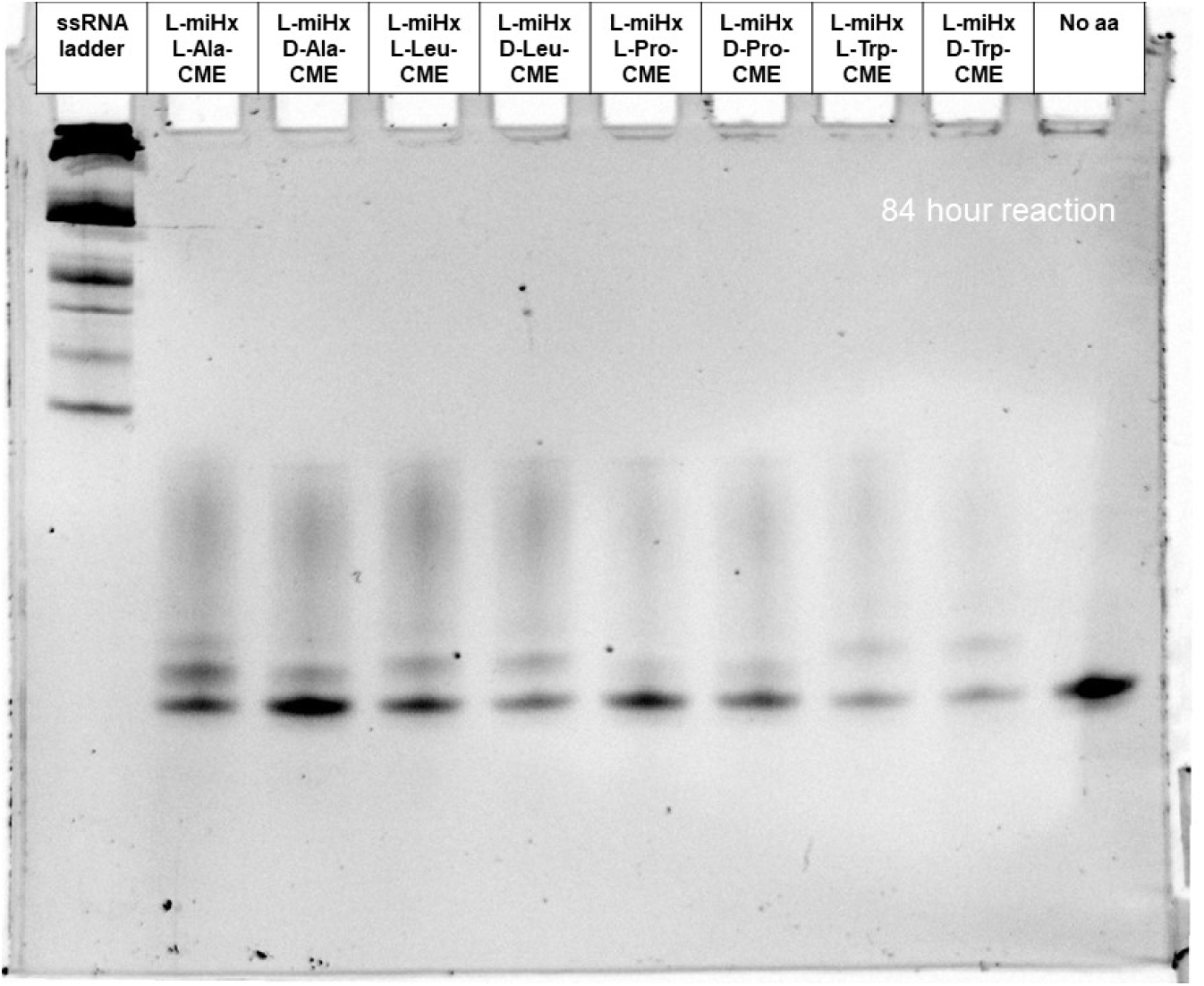
Gel of eutectic nonenzymatic acylation using L-miHx RNA to charge with L/D-Ala-CME or L/D-Leu-CME or L/D-Pro-CME or L/D-Trp-CME for 84 hours at pH 7.5.

**Figure S34.**
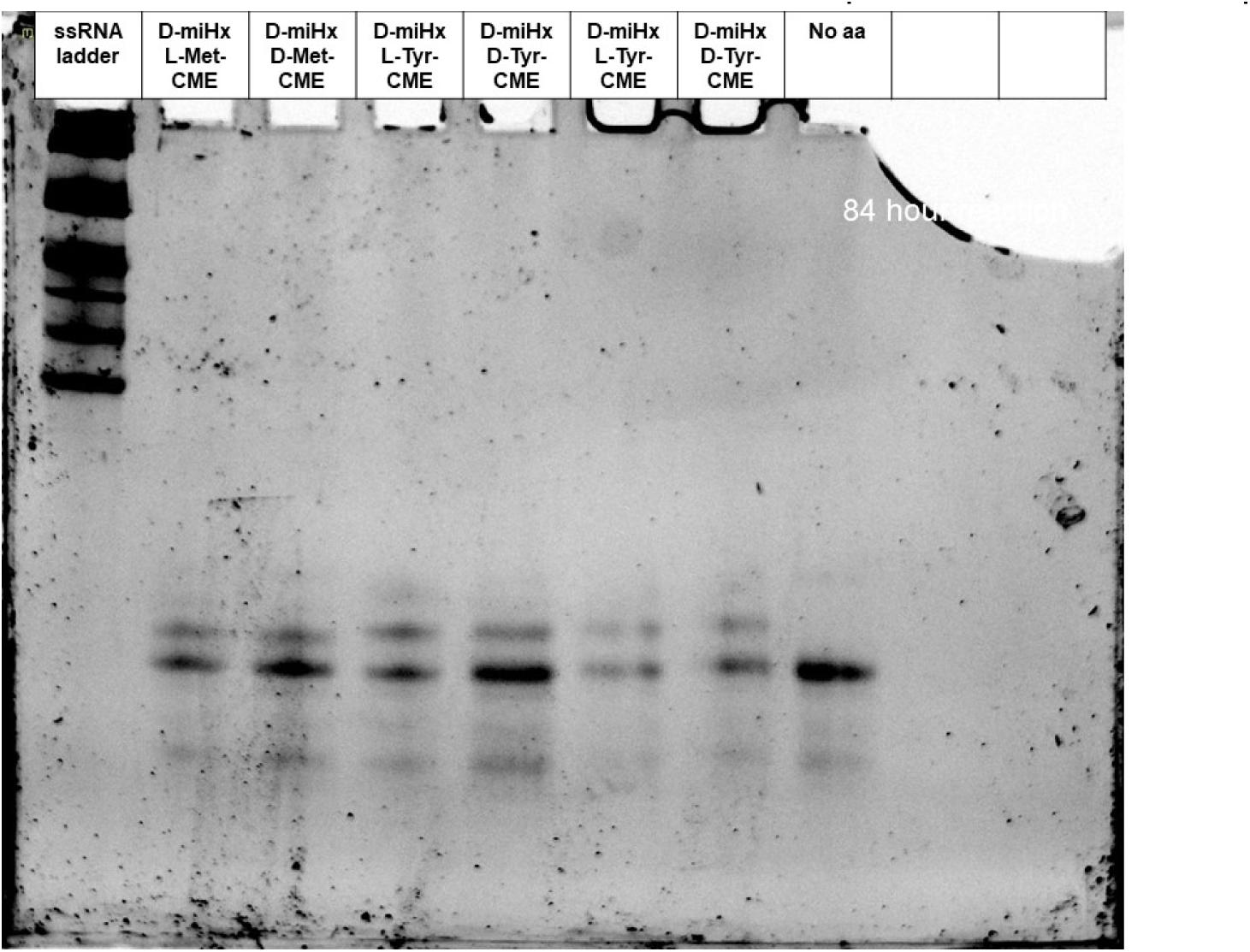
Gel of eutectic nonenzymatic acylation using D-miHx RNA to charge with L/D-Met-CME or L/D-Tyr-CME for 84 hours at pH 7.5.

**Figure S35.**
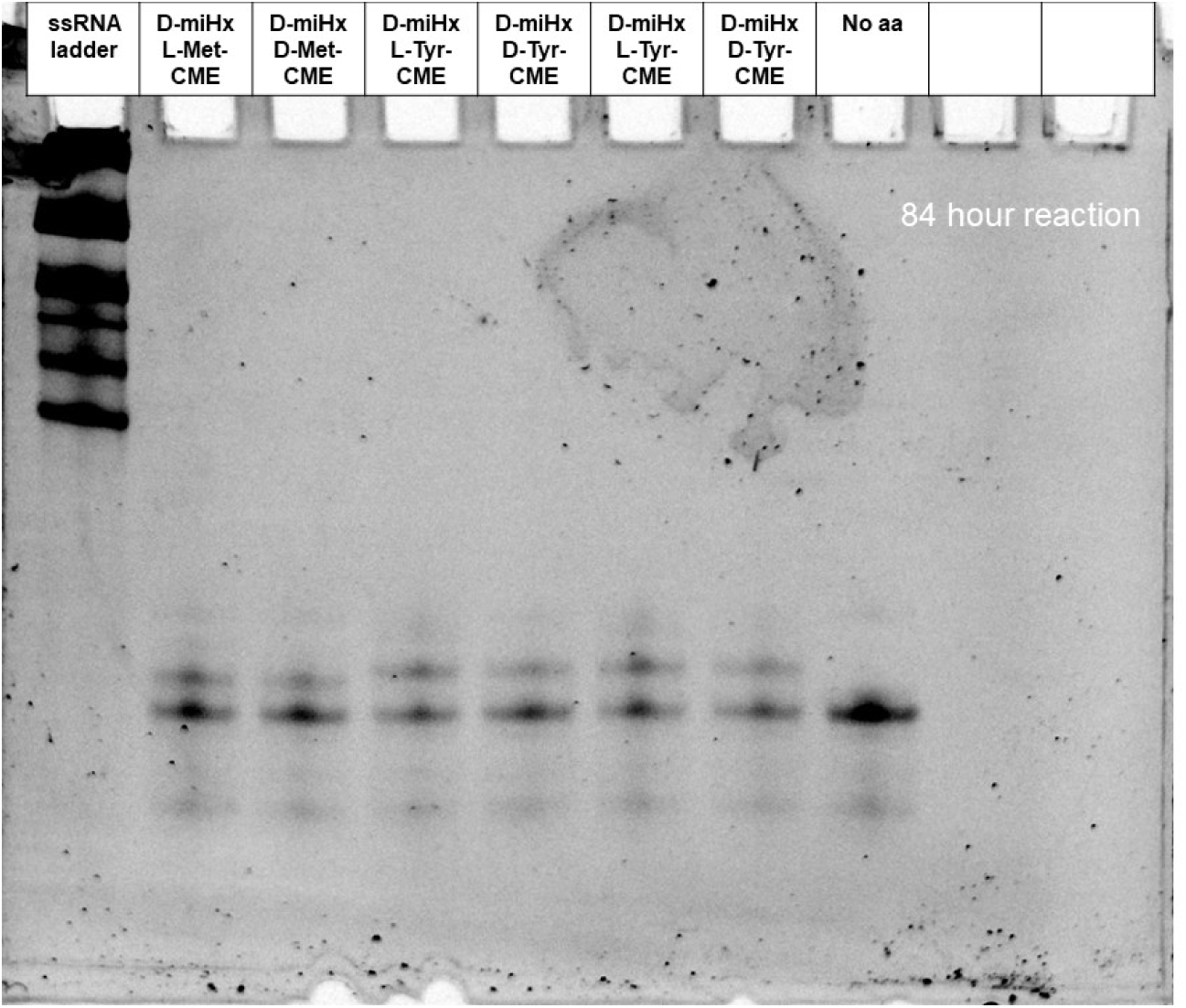
Gel of eutectic nonenzymatic acylation using D-miHx RNA to charge with L/D-Met-CME or L/D-Tyr-CME for 84 hours at pH 7.5.

**Figure S36.**
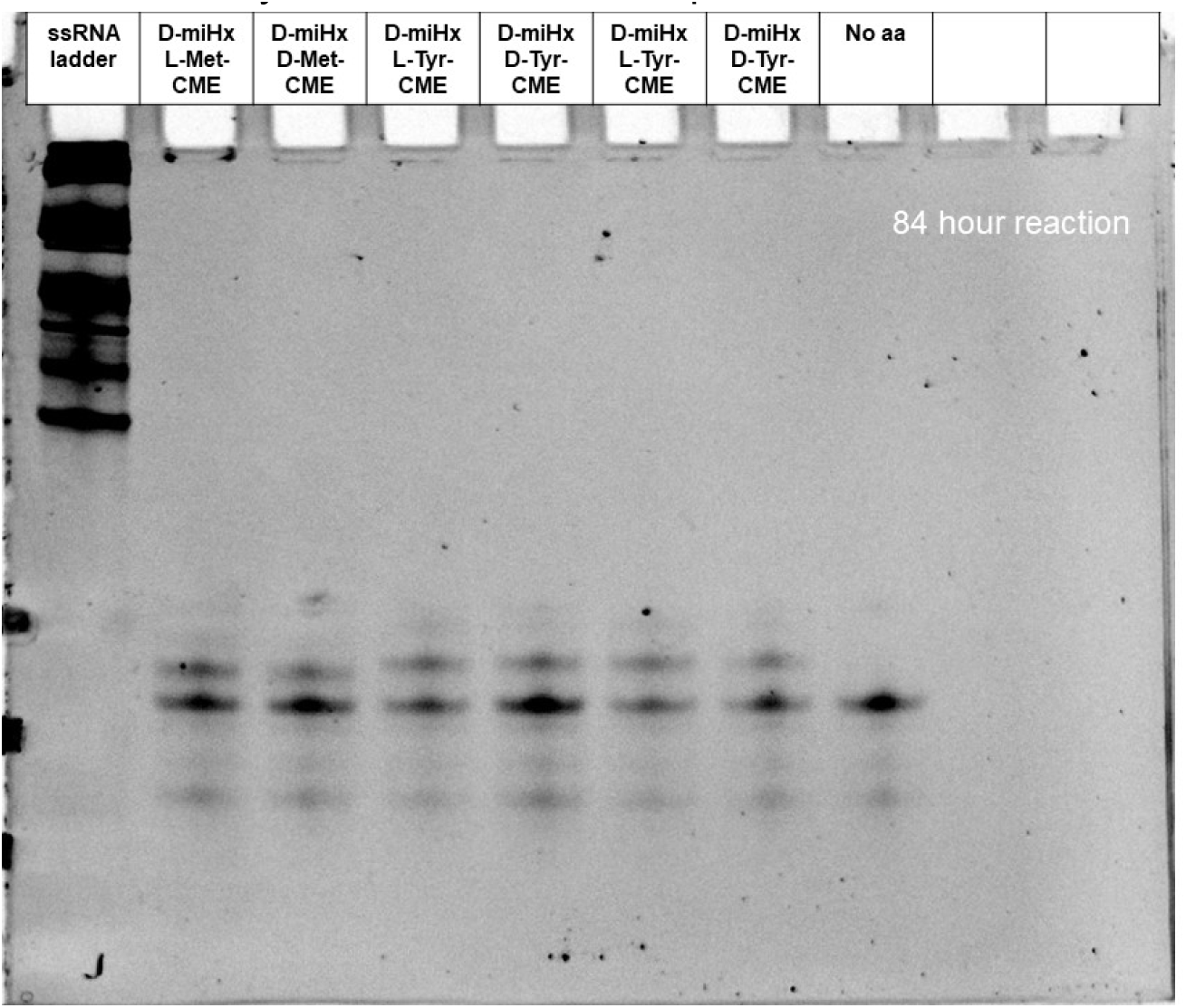
Gel of eutectic nonenzymatic acylation using D-miHx RNA to charge with L/D-Met-CME or L/D-Tyr-CME for 84 hours at pH 7.5.

**Figure S37.**
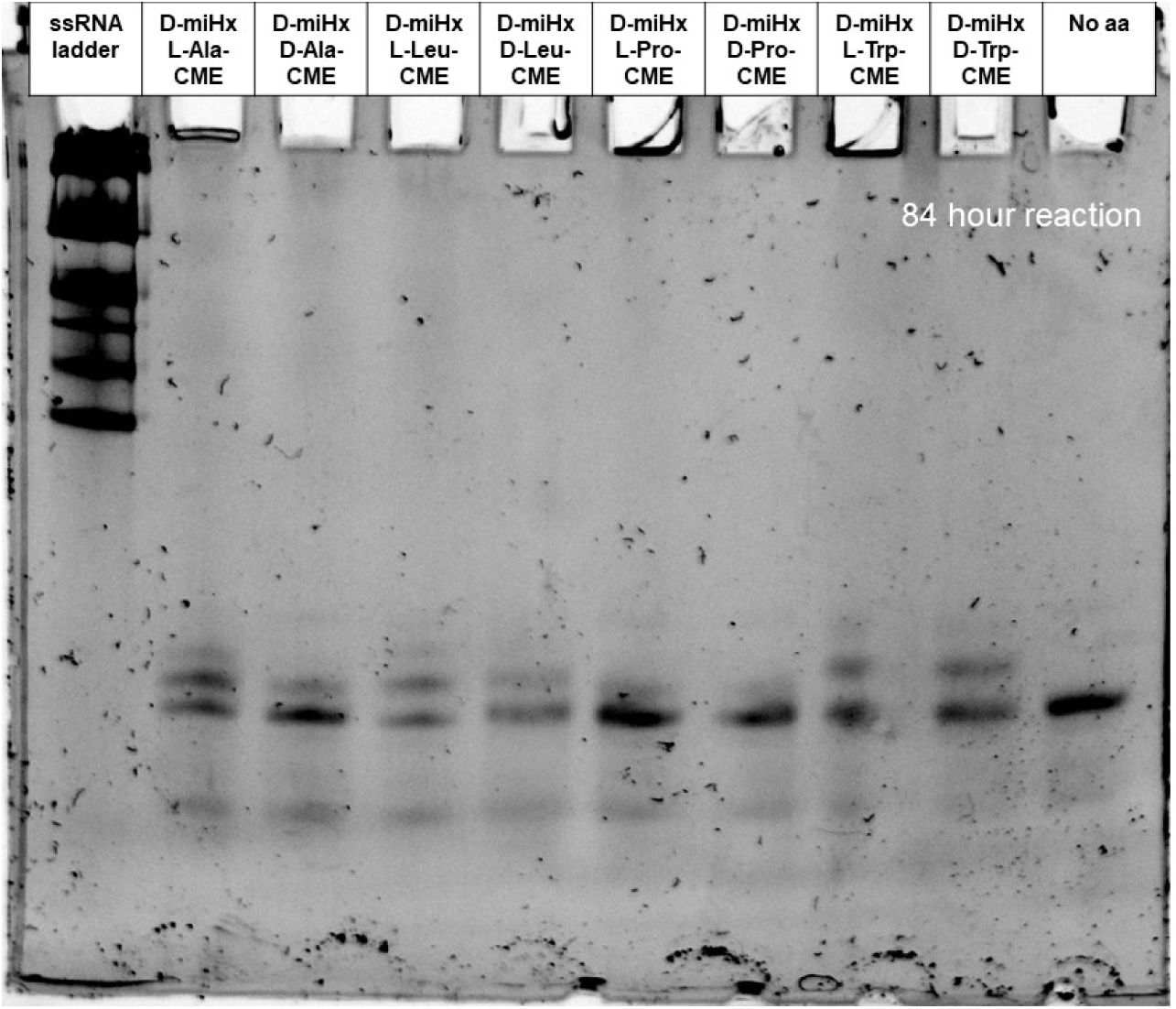
Gel of eutectic nonenzymatic acylation using D-miHx RNA to charge with L/D-Ala-CME or L/D-Leu-CME or L/D-Pro-CME or L/D-Trp-CME for 84 hours at pH 7.5.

**Figure S38.**
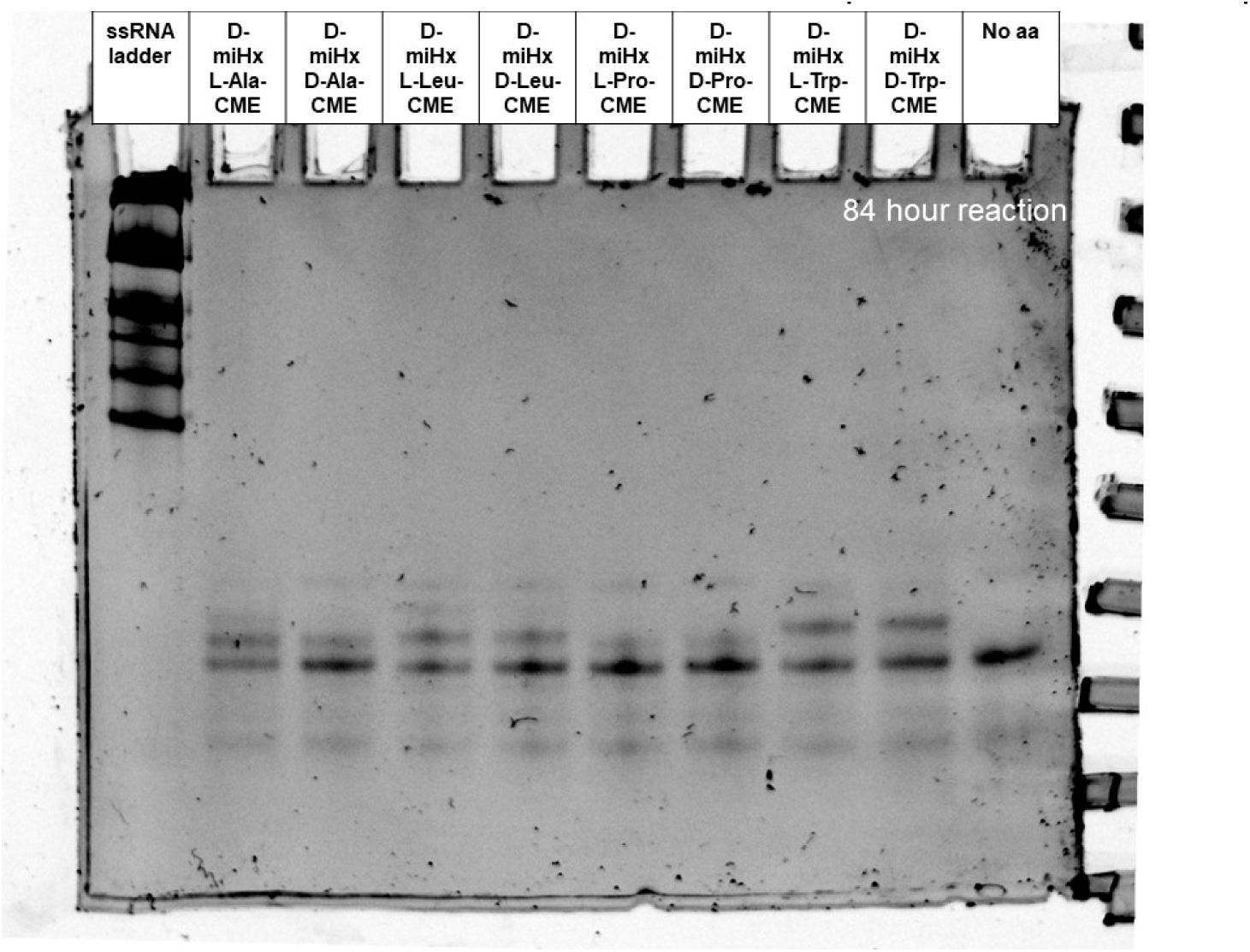
Gel of eutectic nonenzymatic acylation using D-miHx RNA to charge with L/D-Ala-CME or L/D-Leu-CME or L/D-Pro-CME or L/D-Trp-CME for 84 hours at pH 7.5.

**Figure S39.**
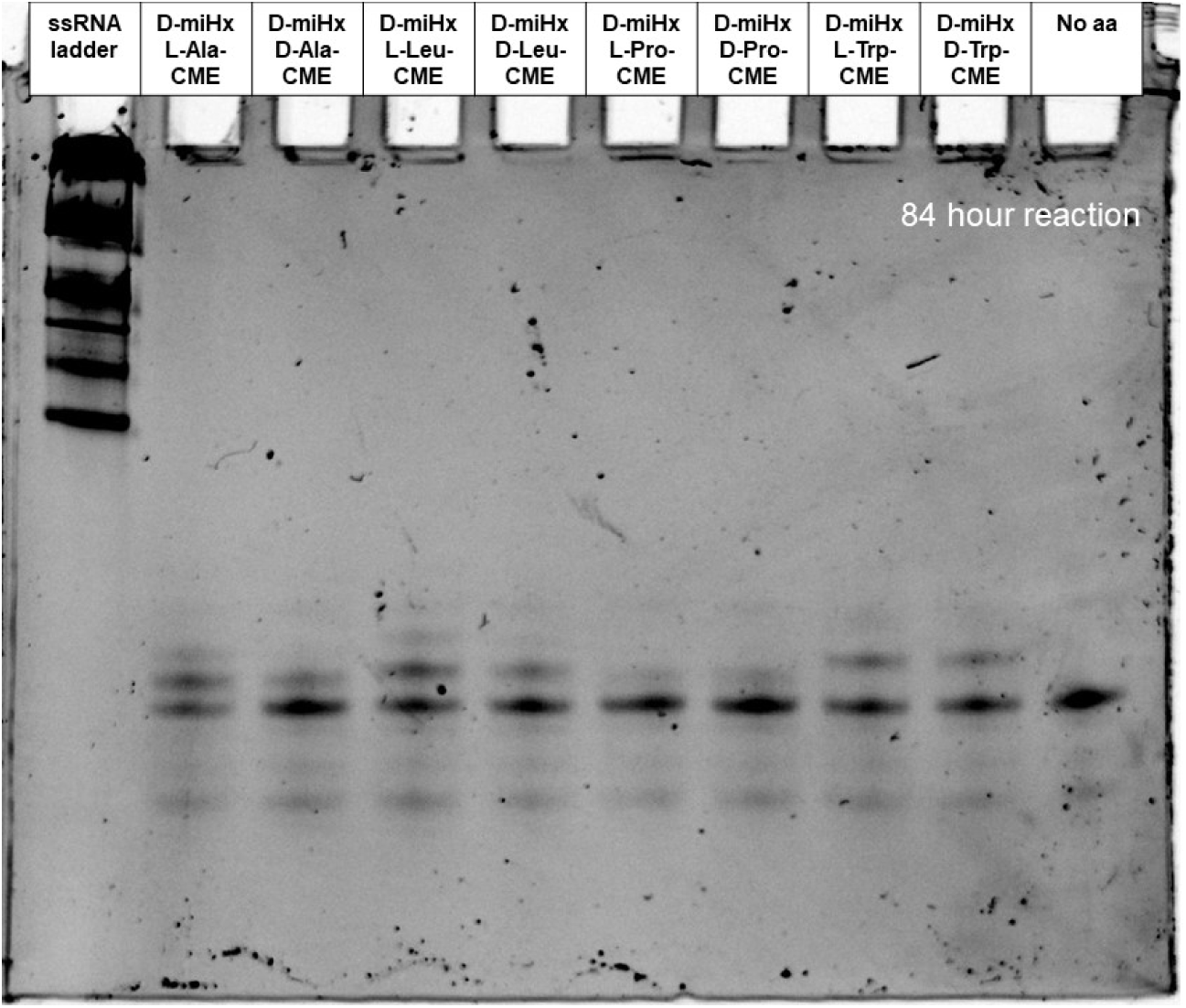
Gel of eutectic nonenzymatic acylation using D-miHx RNA to charge with L/D-Ala-CME or L/D-Leu-CME or L/D-Pro-CME or L/D-Trp-CME for 84 hours at pH 7.5.

**Figure S40.**
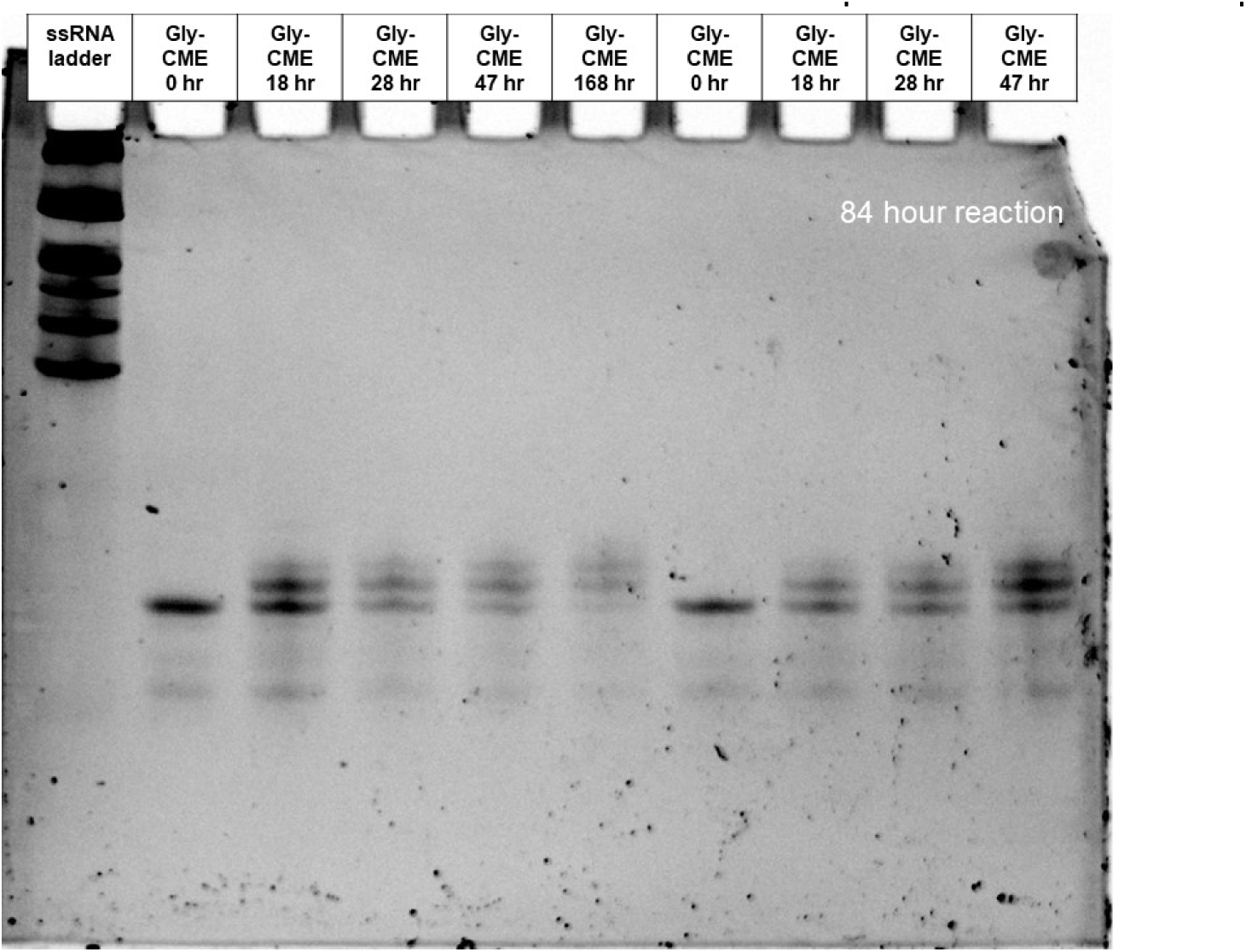
Gel of eutectic nonenzymatic charging reaction tracking acylation yield over time using glycine-CME from 0-168 hours at pH 7.5.

**Figure S41.**
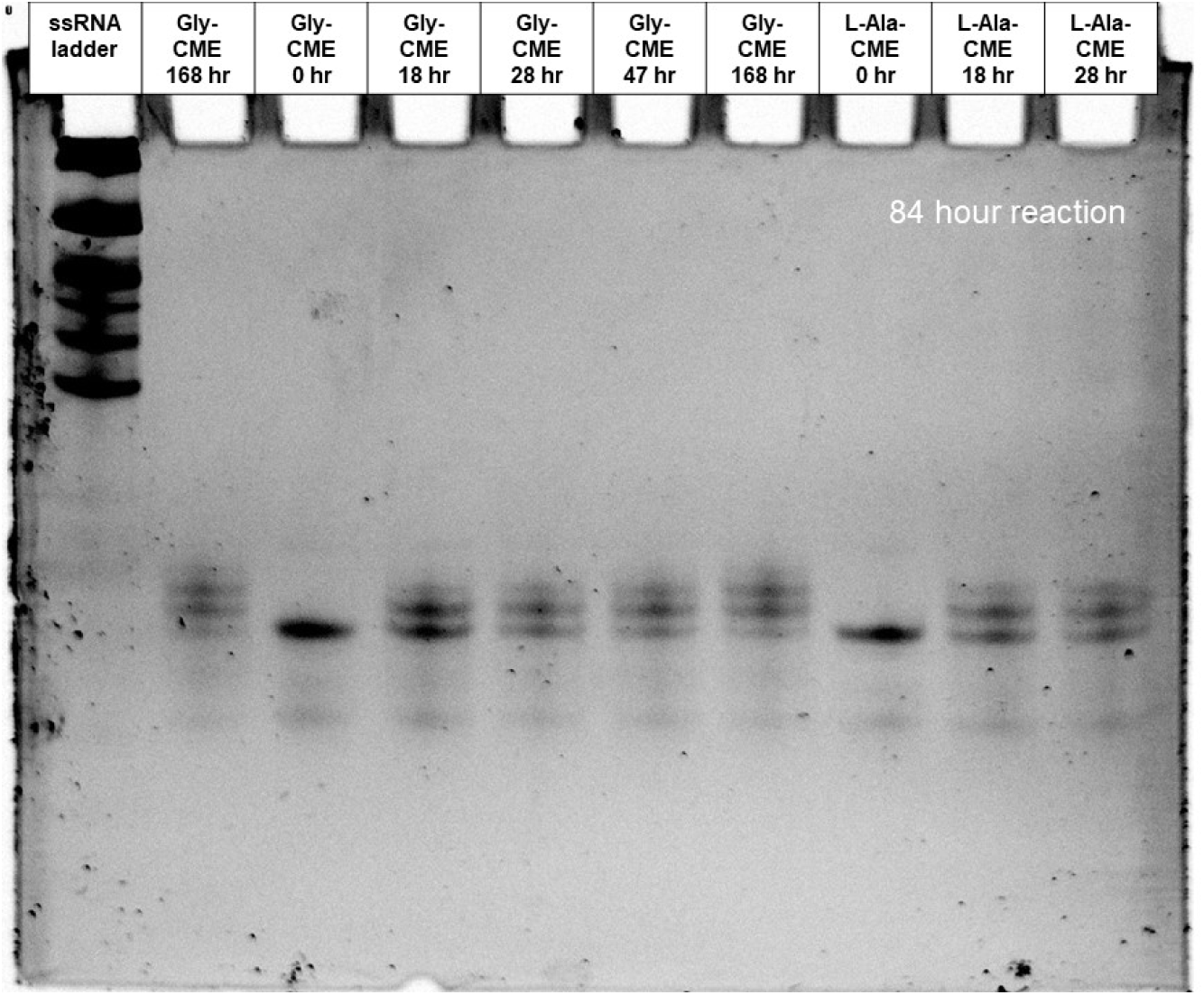
Gel of eutectic nonenzymatic charging reaction tracking acylation yield over time using glycine-CME or L-alanine-CME from 0-168 hours at pH 7.5.

**Figure S42.**
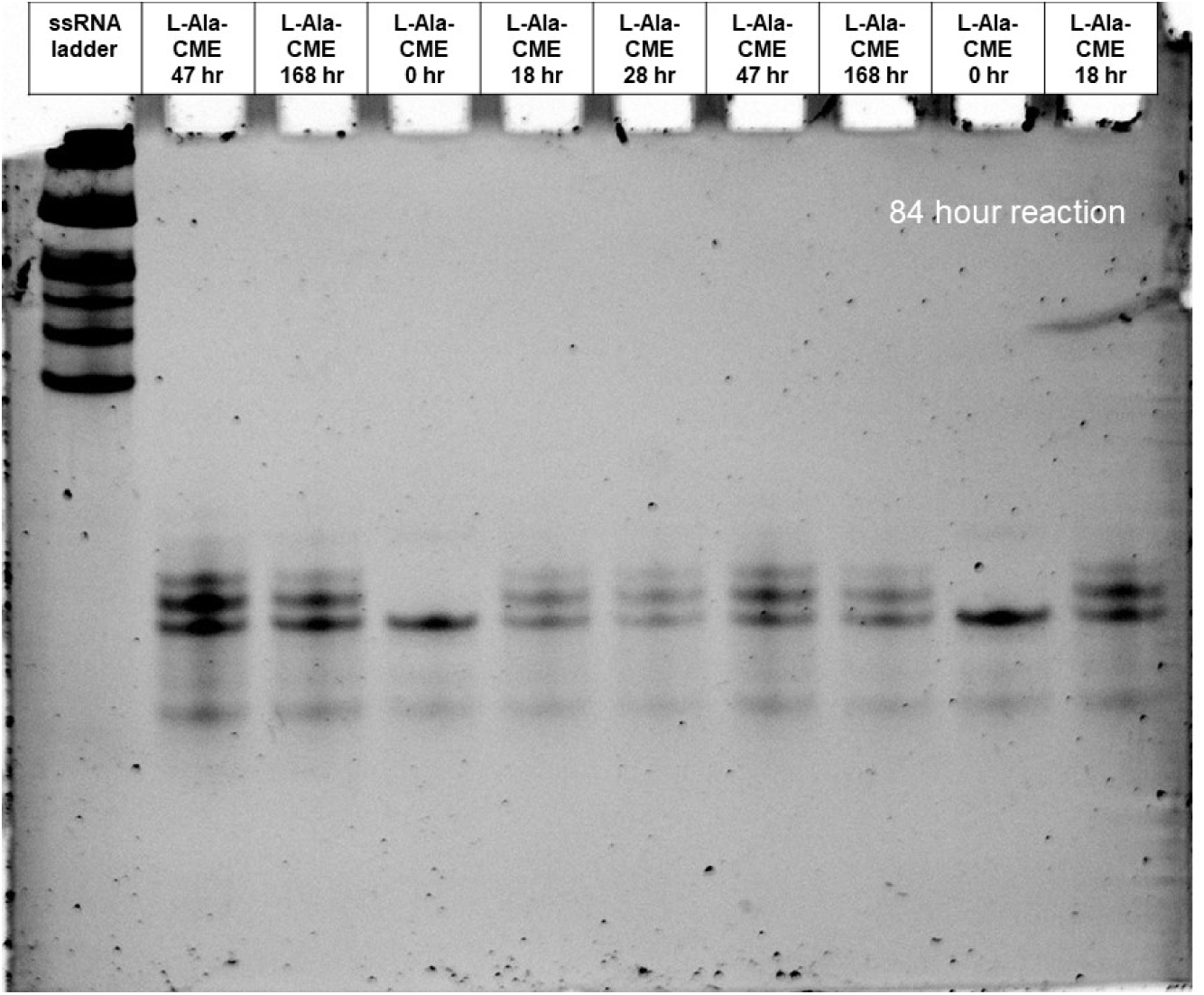
Gel of eutectic nonenzymatic charging reaction tracking acylation yield over time using L-alanine-CME from 0-168 hours at pH 7.5.

**Figure S43.**
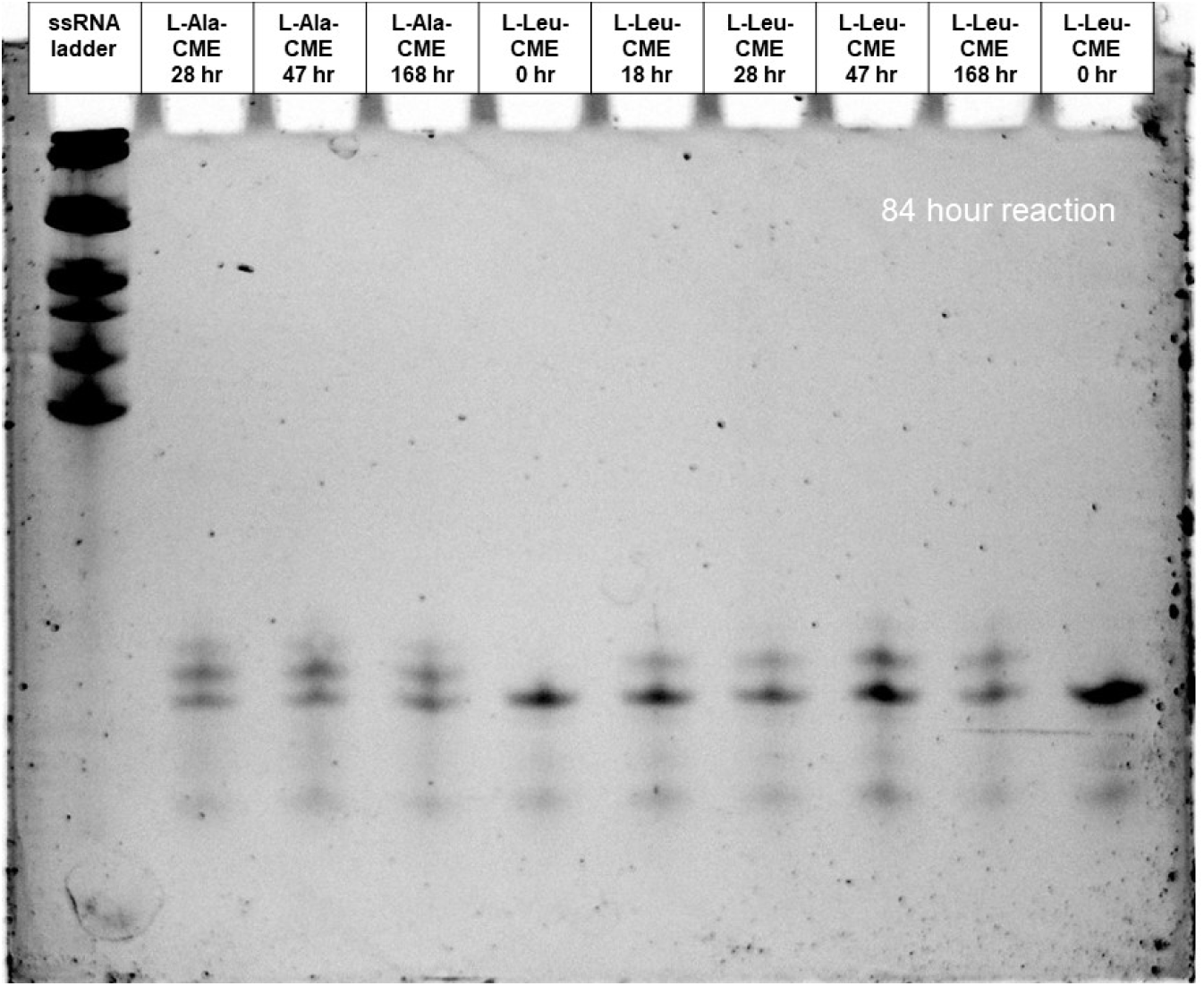
Gel of eutectic nonenzymatic charging reaction tracking acylation yield over time using L-alanine-CME or L-leucine-CME from 0-168 hours at pH 7.5.

**Figure S44.**
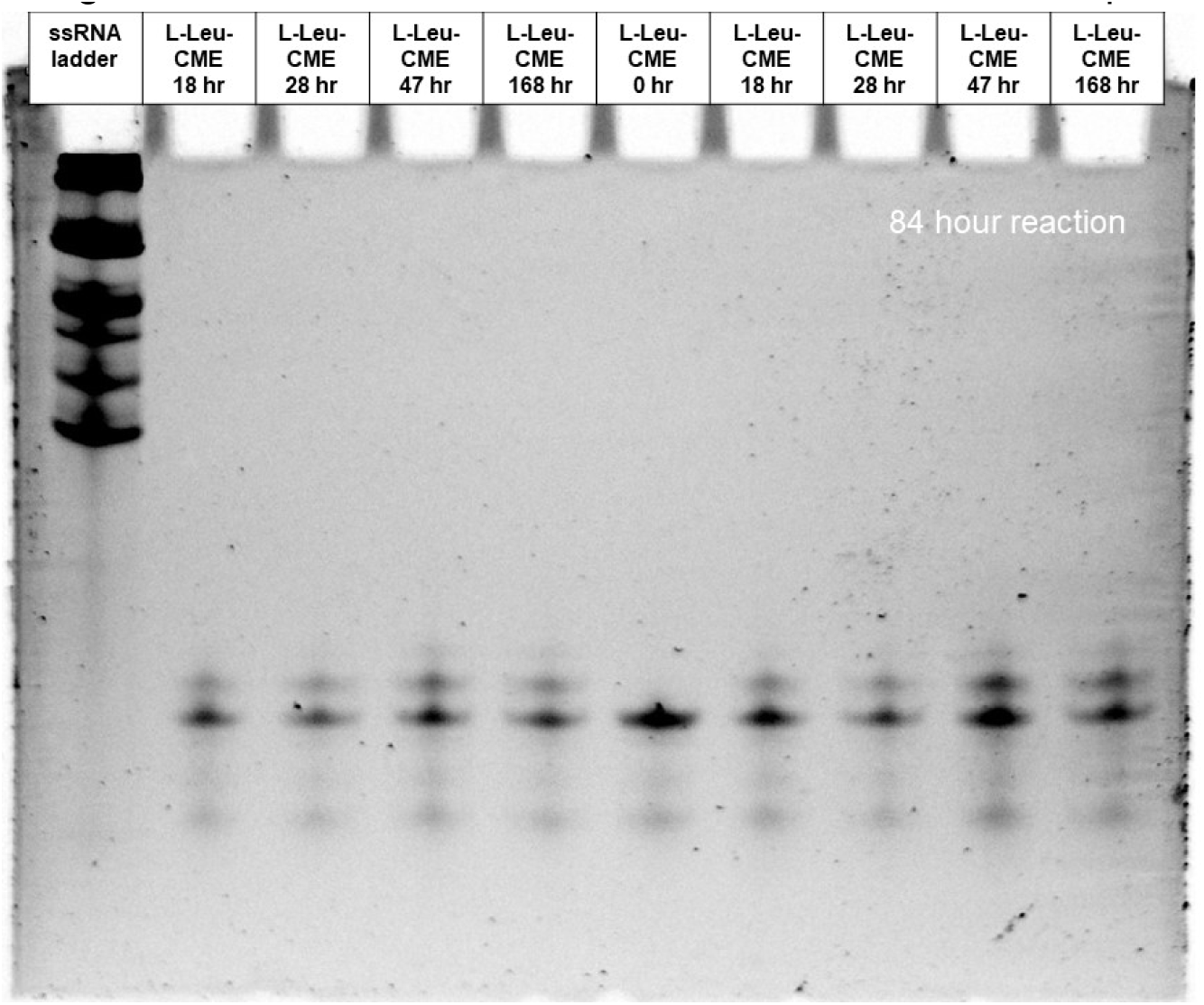
Gel of eutectic nonenzymatic charging reaction tracking acylation yield over time using L-leucine-CME from 0-168 hours at pH 7.5.

**Figure S45.**
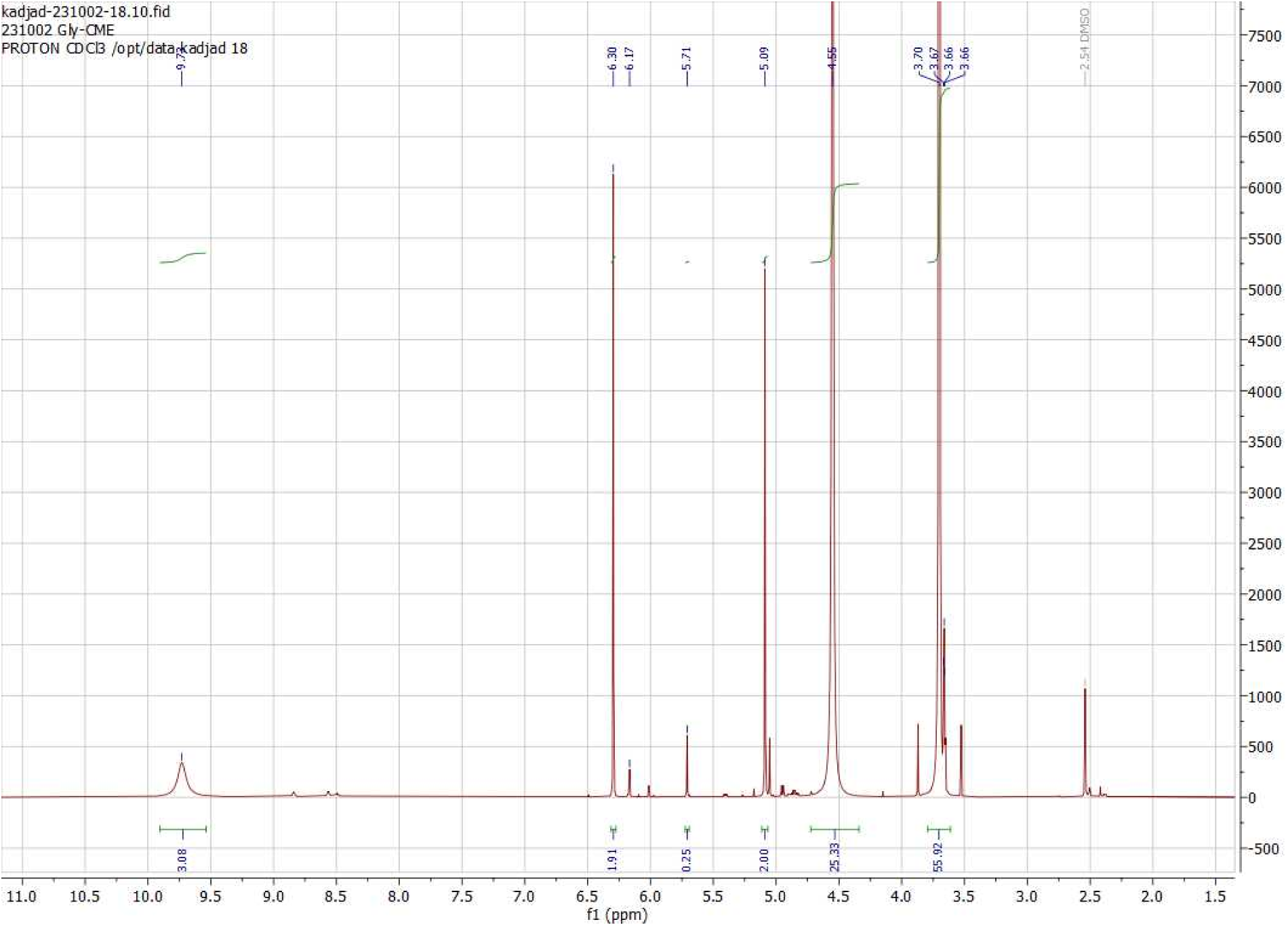
^1^H NMR of glycine-cyanomethylester

**Figure S46.**
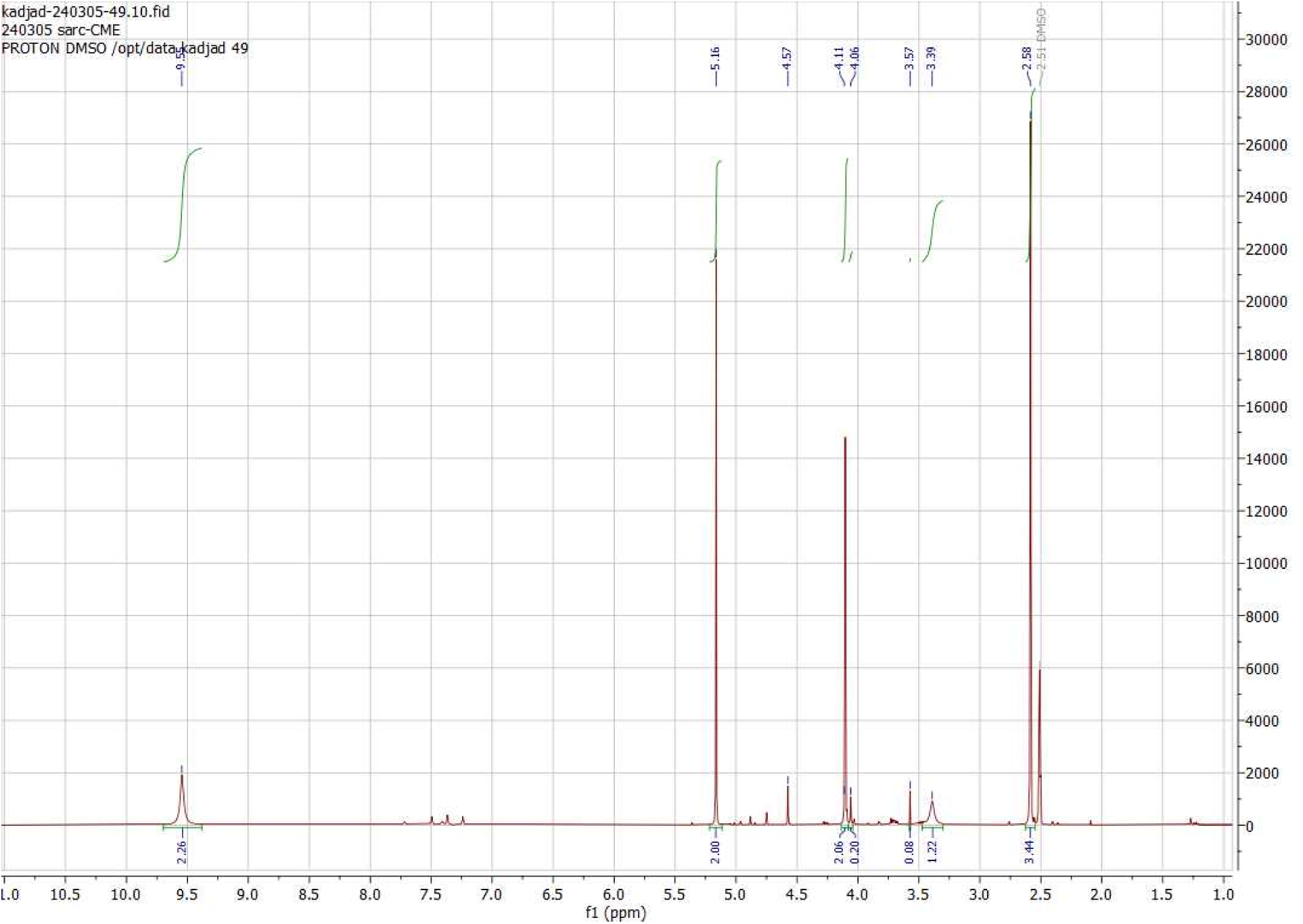
^1^H NMR of sarcosine-cyanomethylester

**Figure S47.**
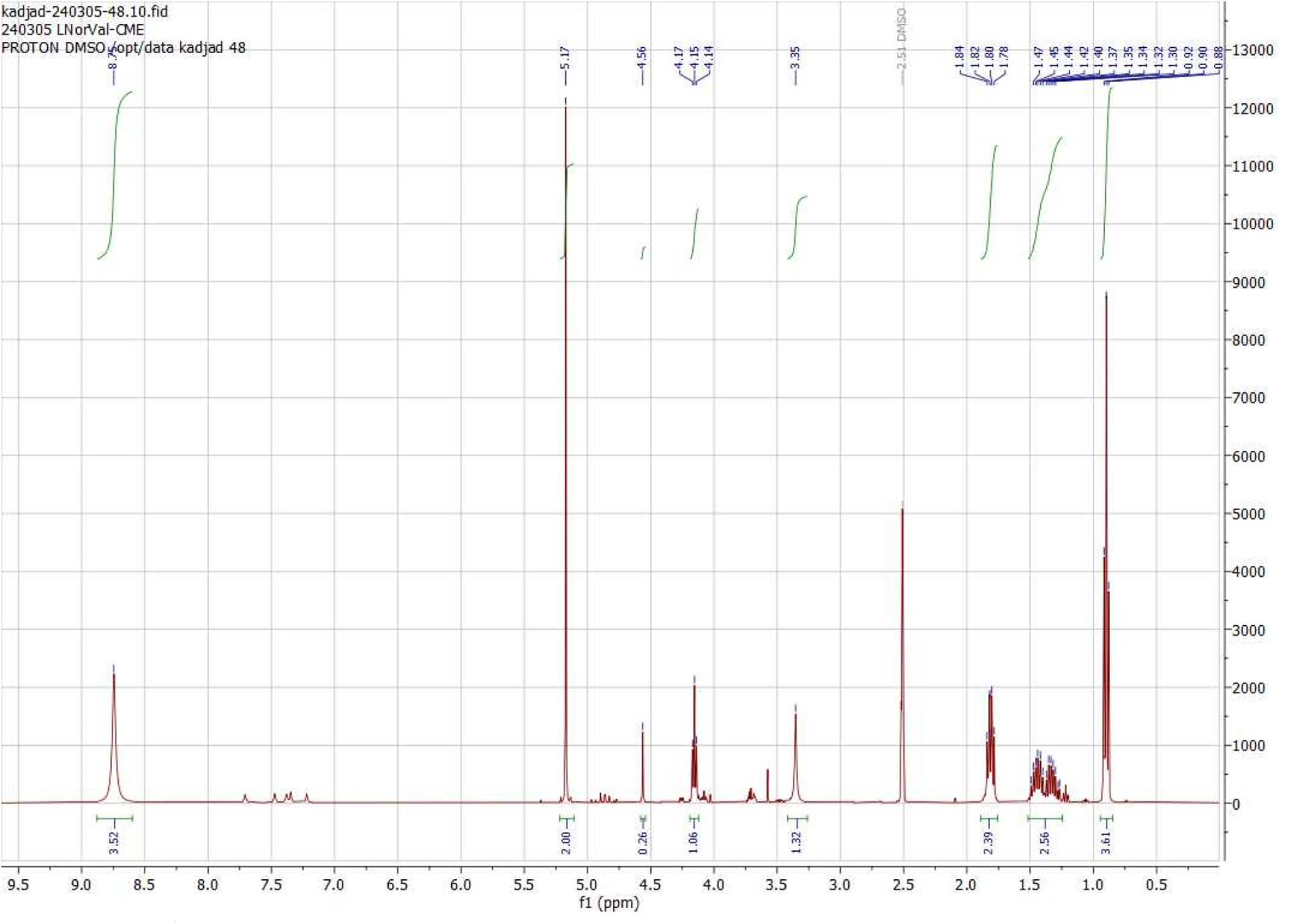
^1^H NMR of L-norvaline-cyanomethylester

**Figure S48.**
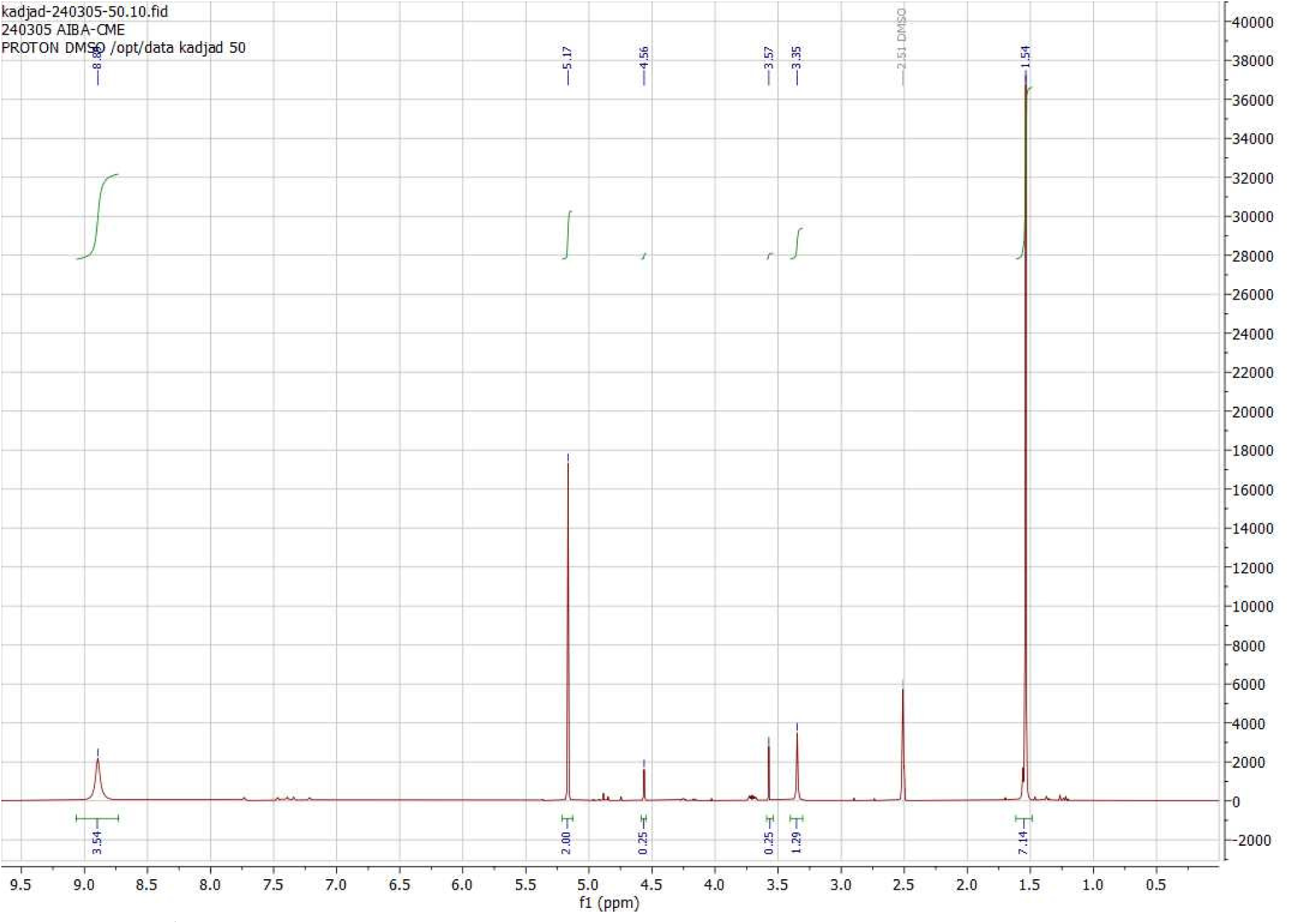
^1^H NMR of aminoisobutyric acid-cyanomethylester

**Figure S49.**
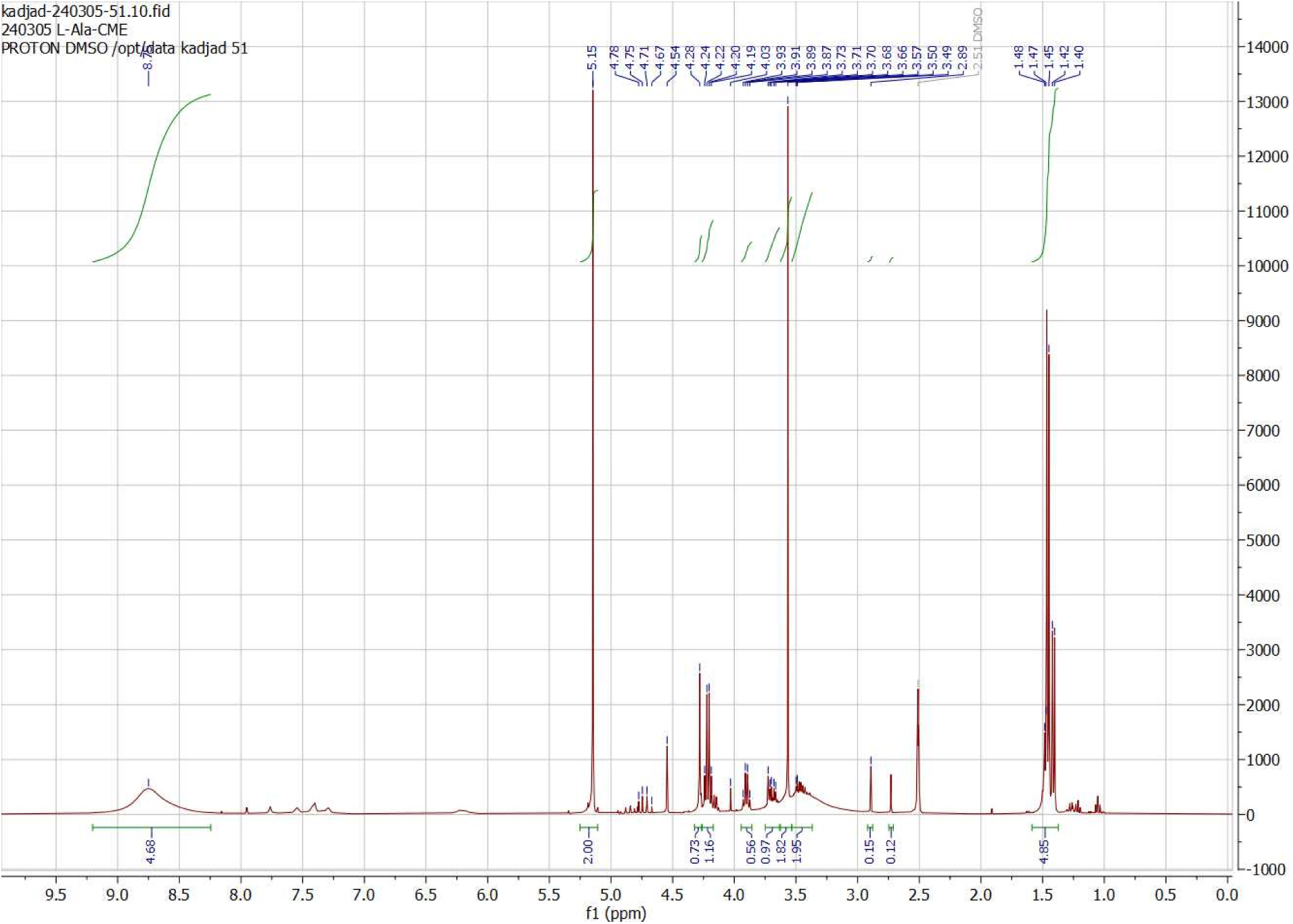
^1^H NMR of L-alanine-cyanomethylester

**Figure S50.**
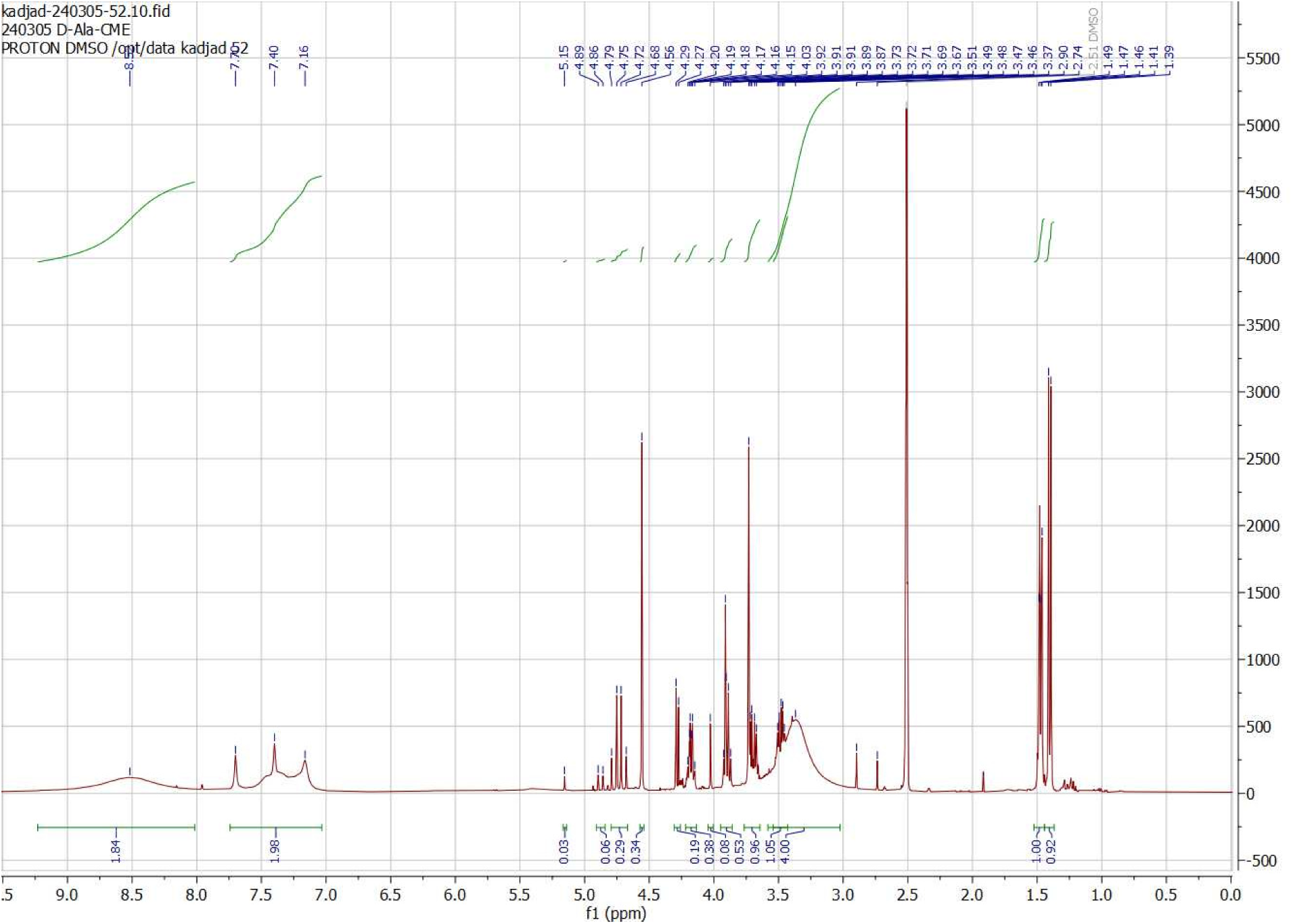
^1^H NMR of D-alanine-cyanomethylester

**Figure S51.**
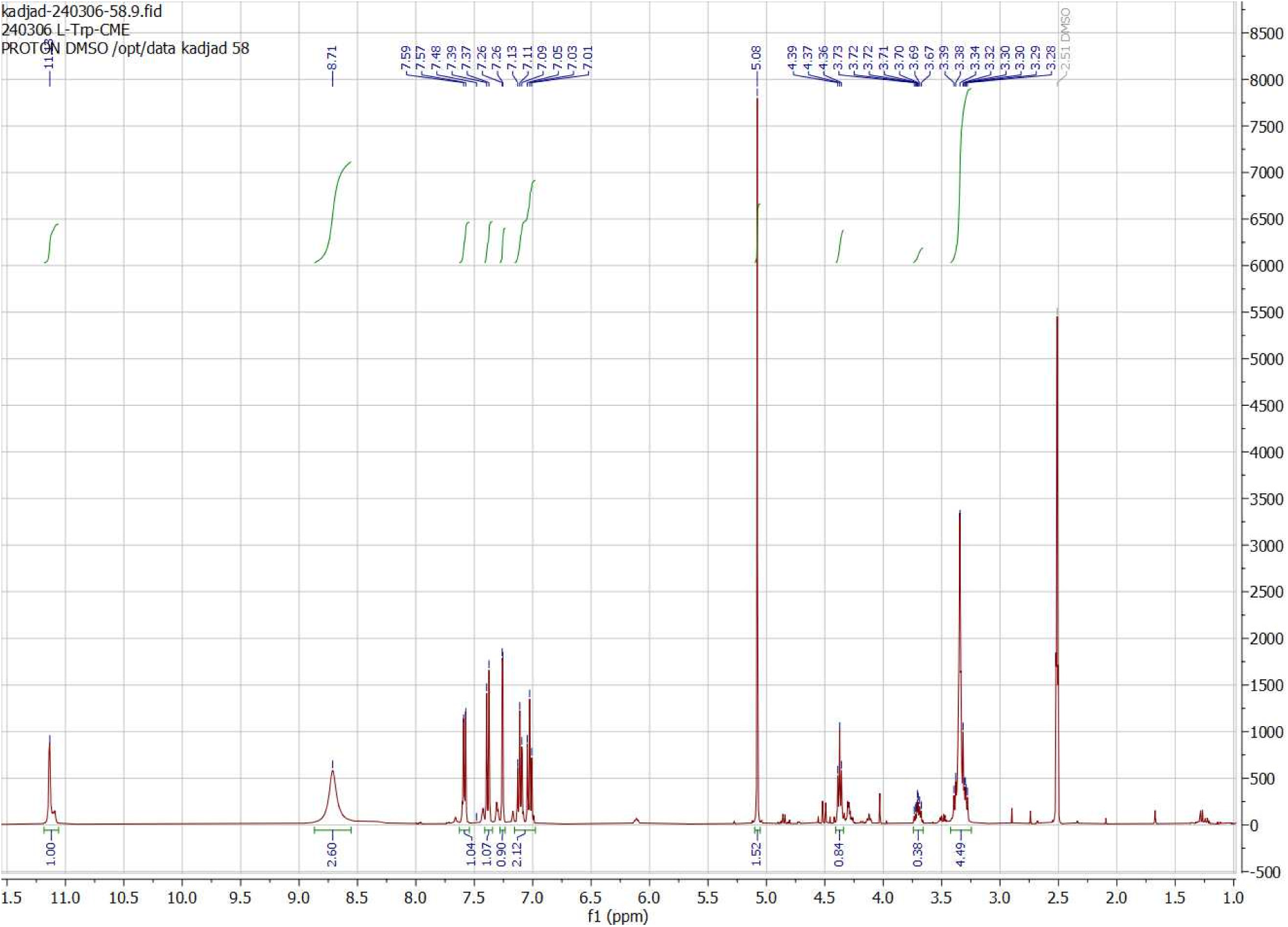
^1^H NMR of L-tryptophan-cyanomethylester

**Figure S52.**
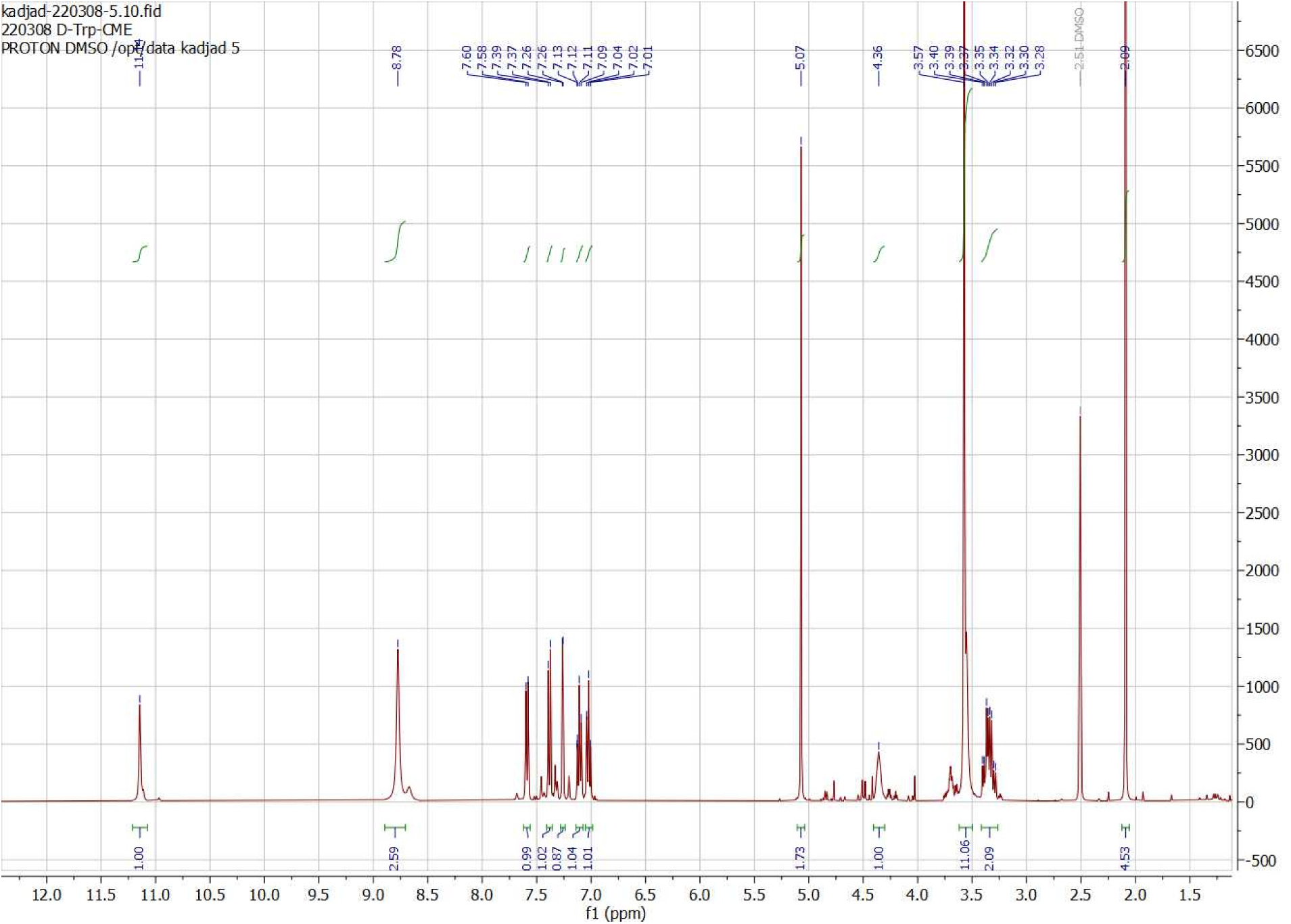
^1^H NMR of D-tryptophan-cyanomethylester

**Figure S53.**
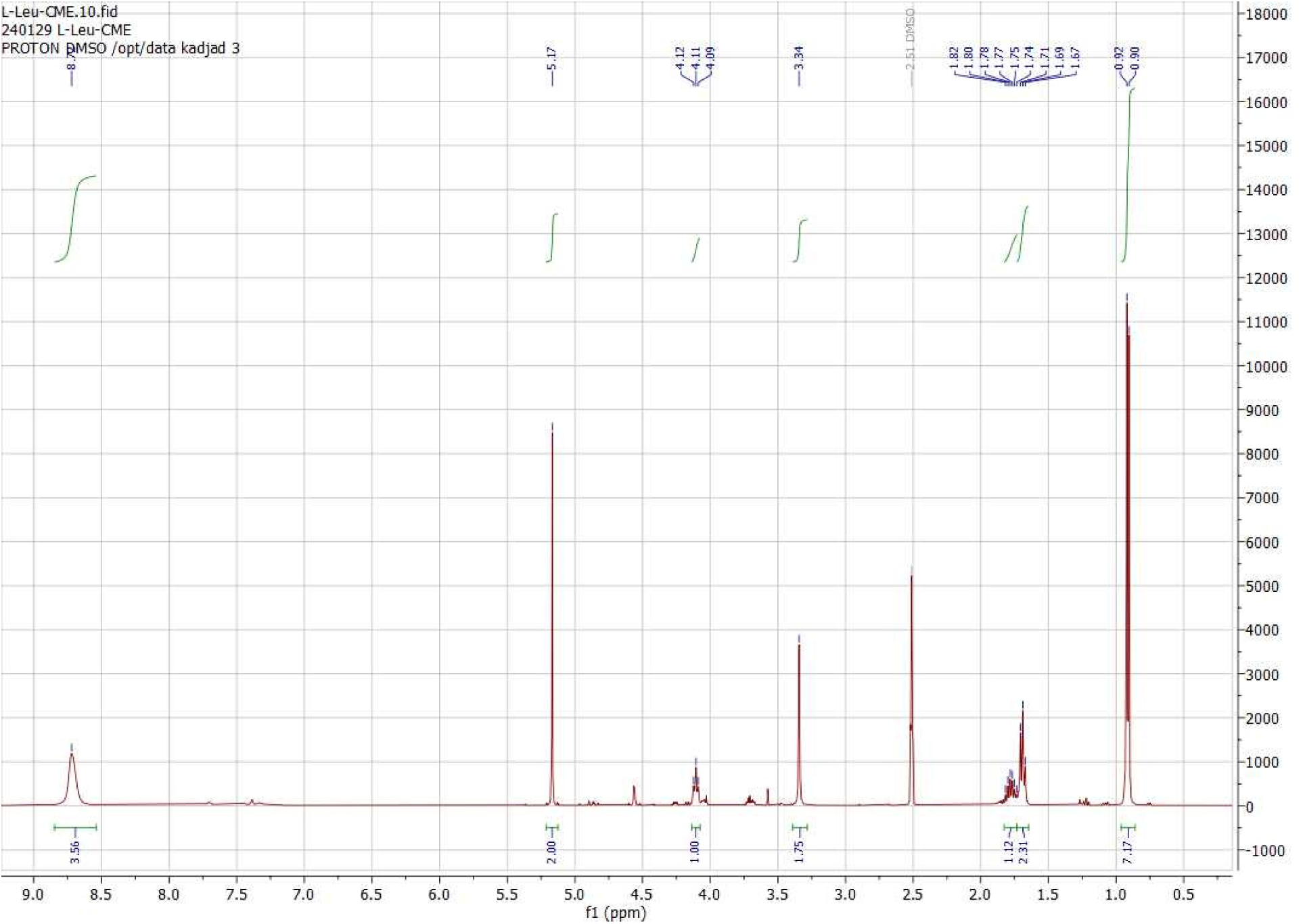
^1^H NMR of L-leucine-cyanomethylester

**Figure S54.**
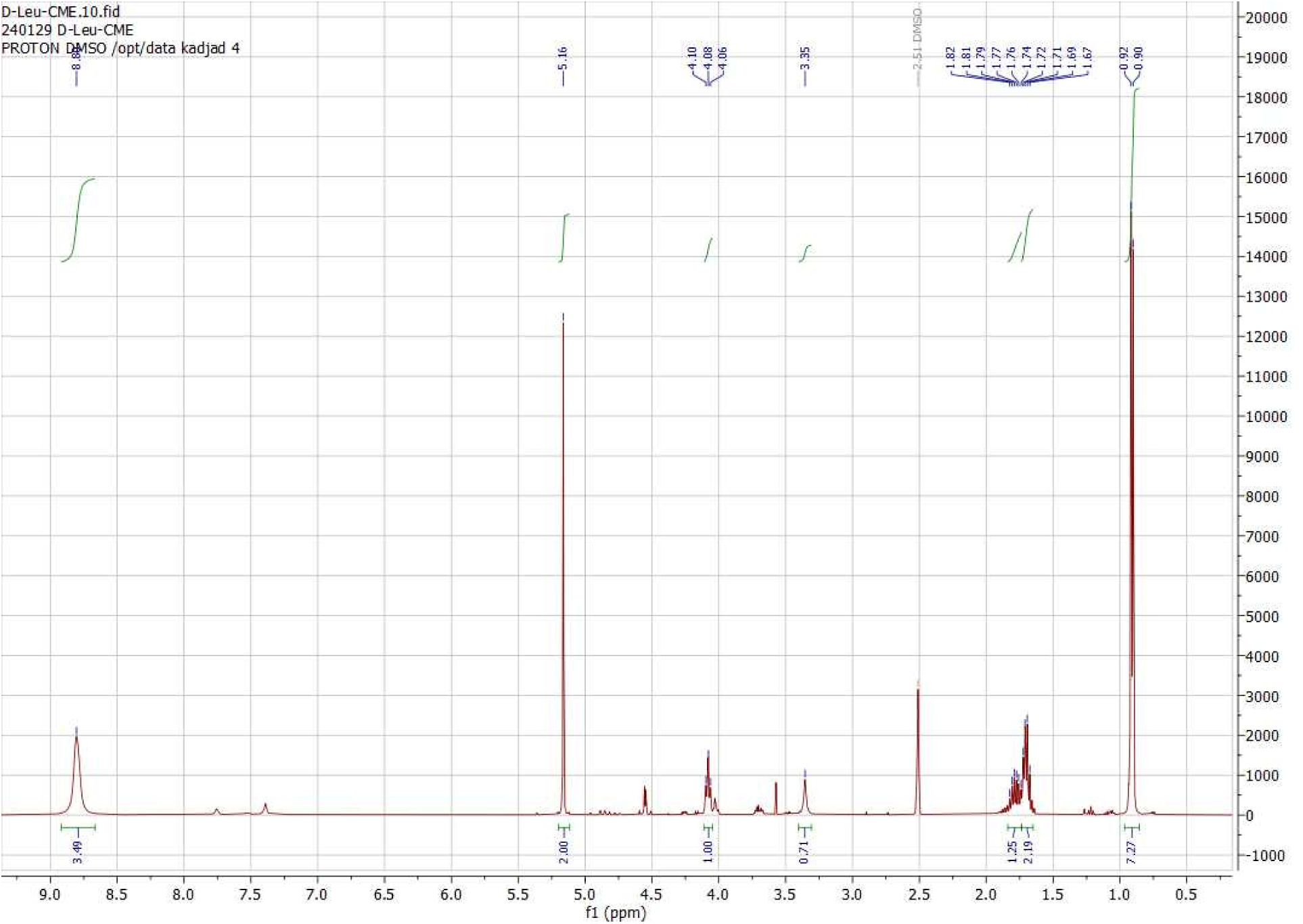
^1^H NMR of D-leucine-cyanomethylester

**Figure S55.**
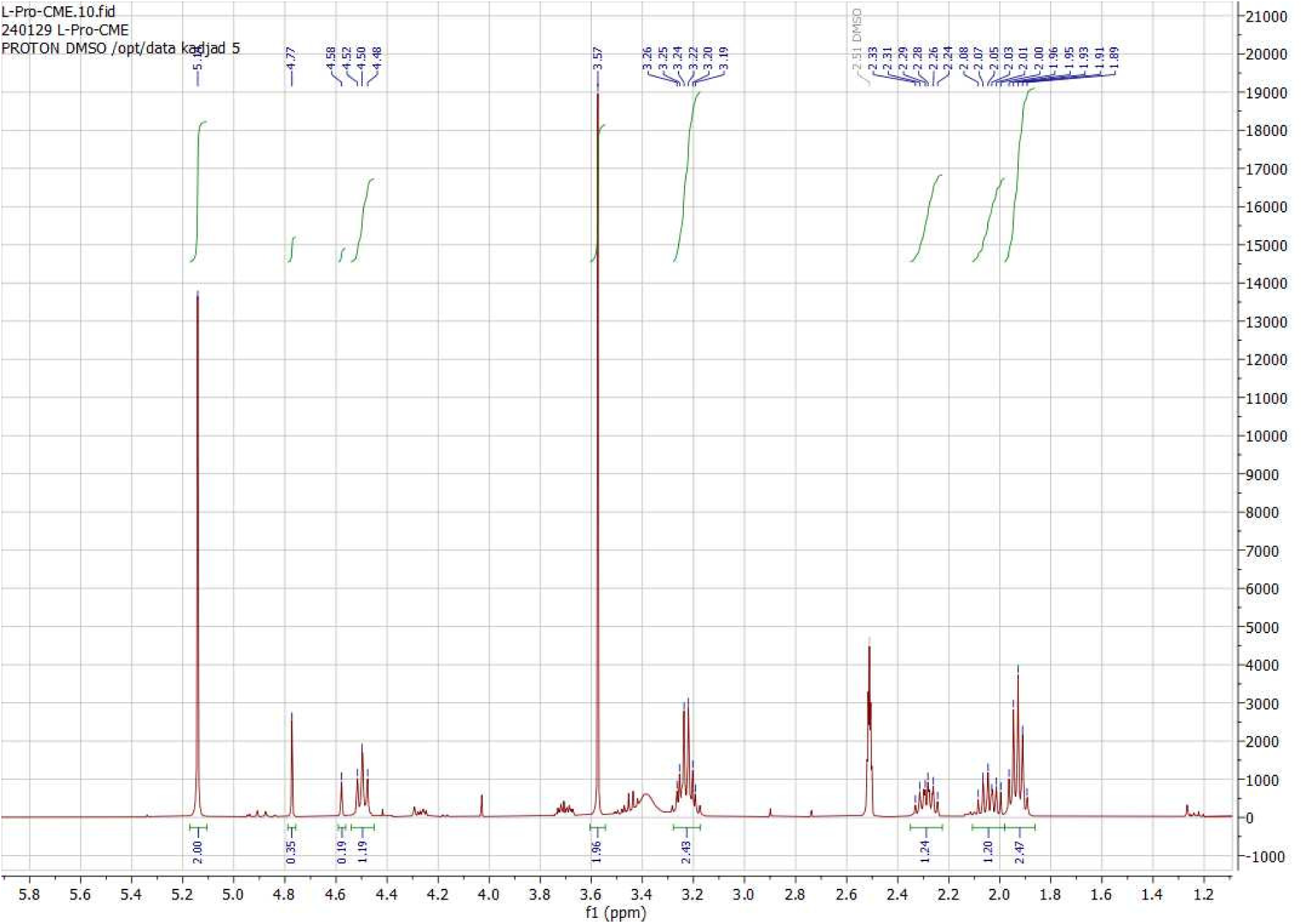
^1^H NMR of L-proline-cyanomethylester

**Figure S56.**
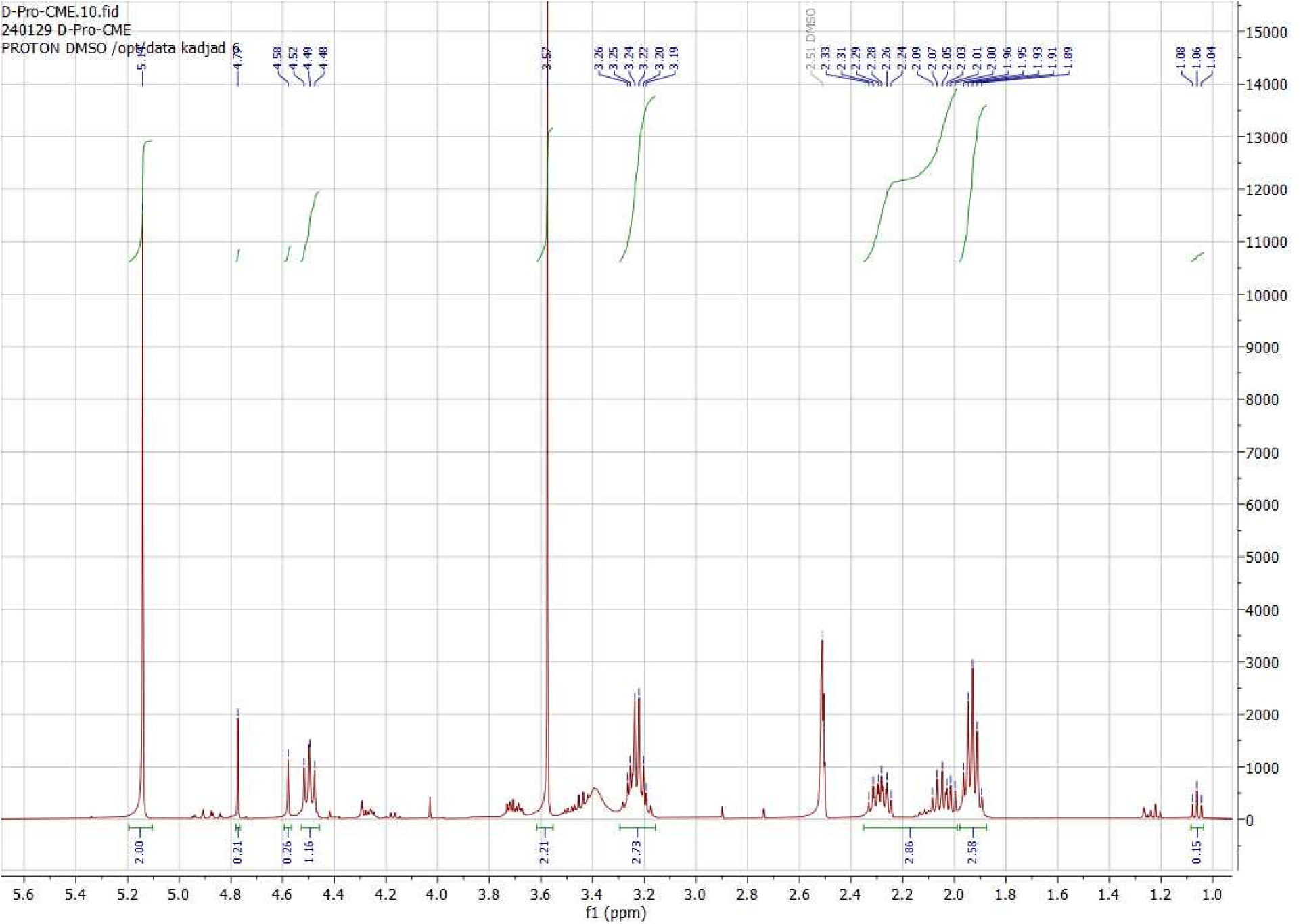
^1^H NMR of D-proline-cyanomethylester

**Figure S57.**
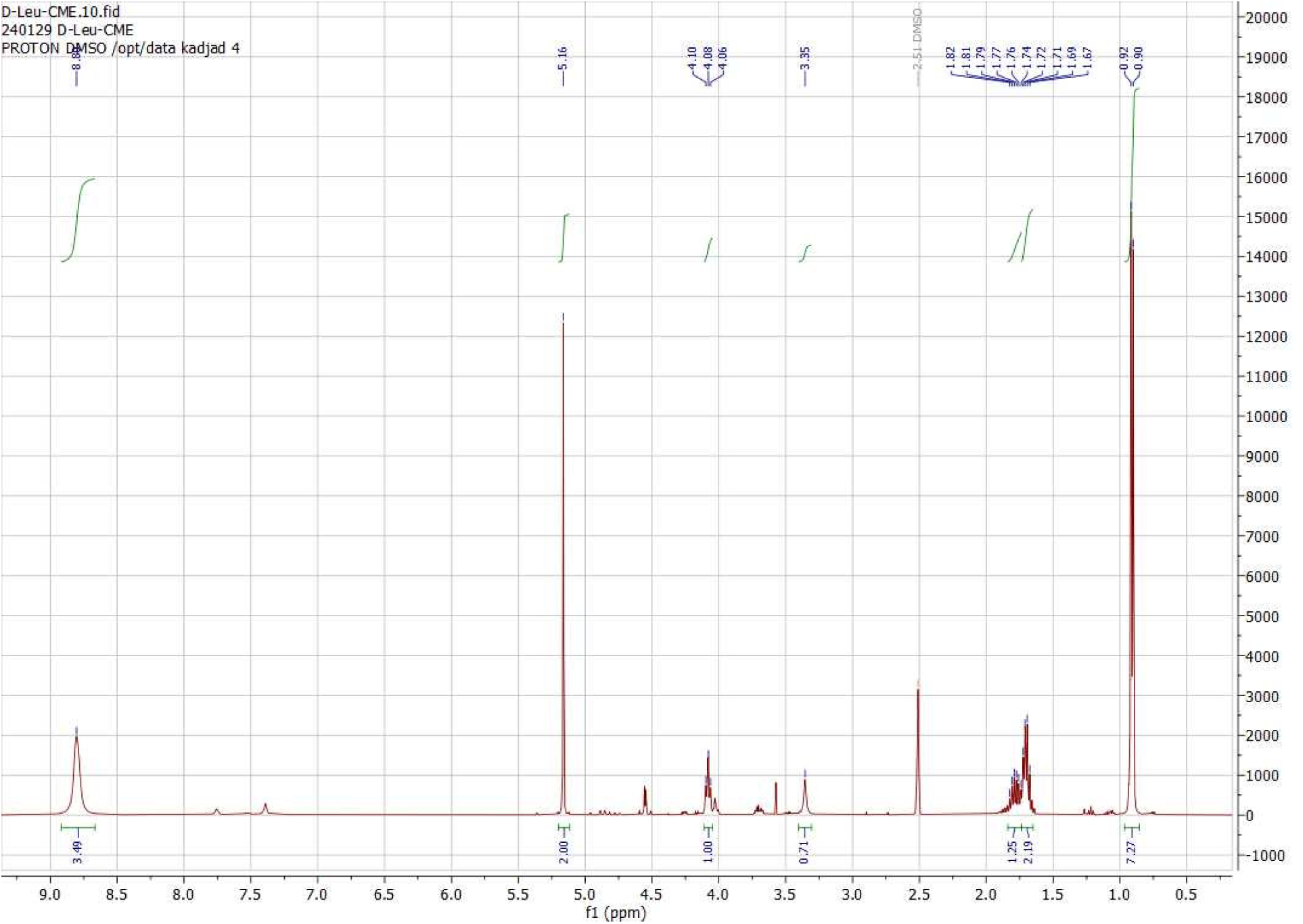
^1^H NMR of L-methionine-cyanomethylester

**Figure S58.**
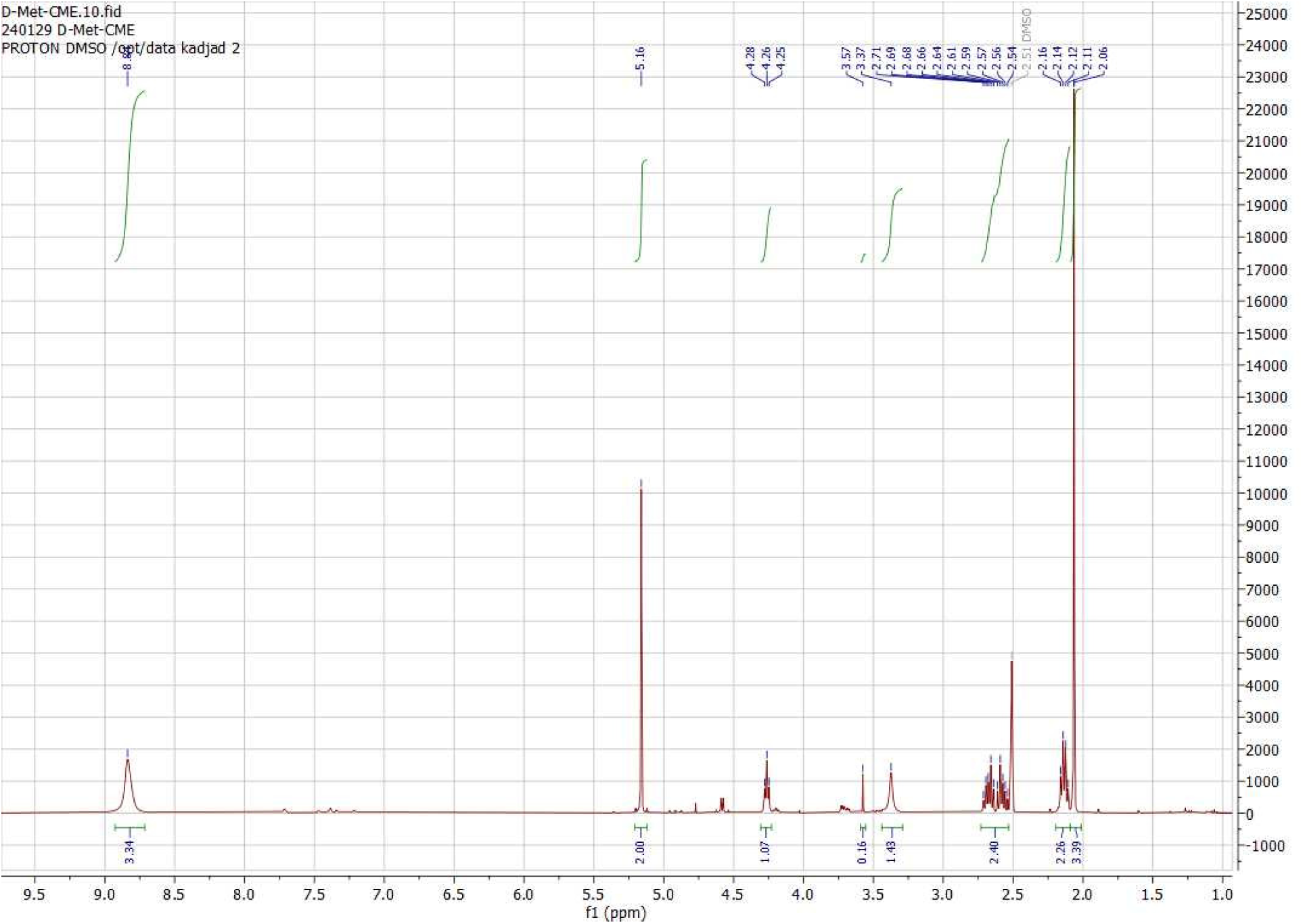
^1^H NMR of D-methionine-cyanomethylester

**Figure S59.**
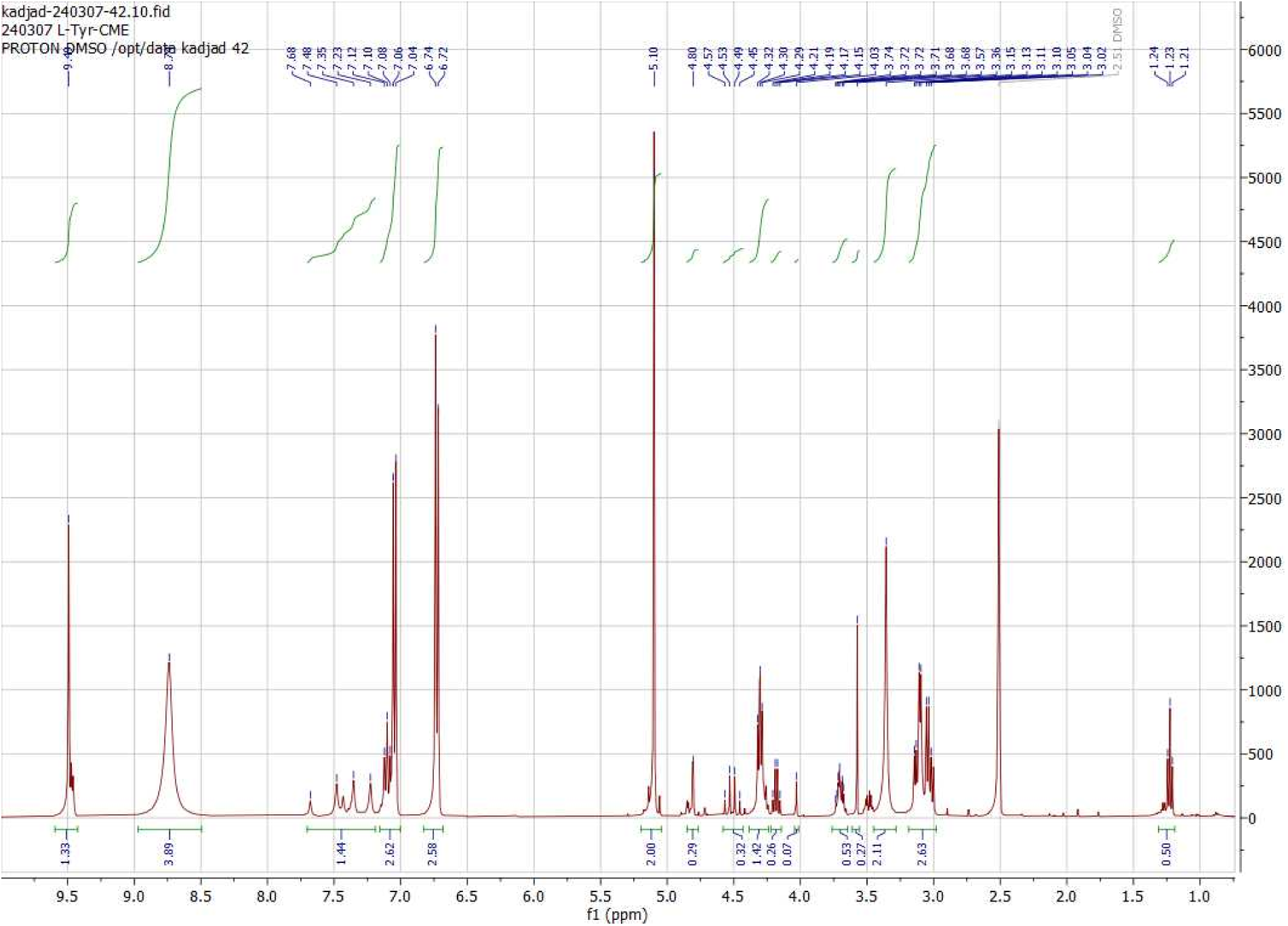
^1^H NMR of L-tyrosine-cyanomethylester

**Figure S60.**
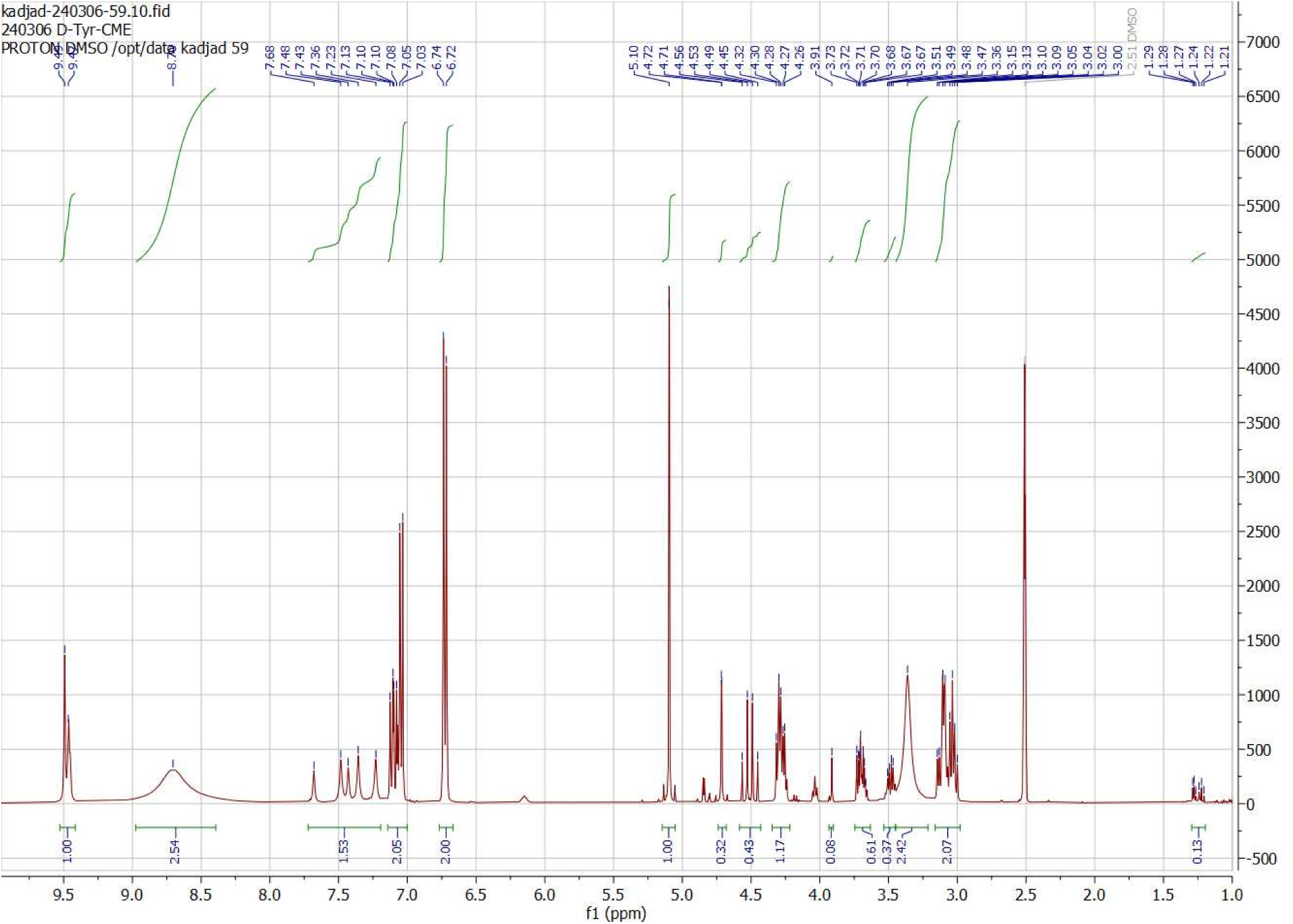
^1^H NMR of D-tyrosine-cyanomethylester

**Figure S61.**
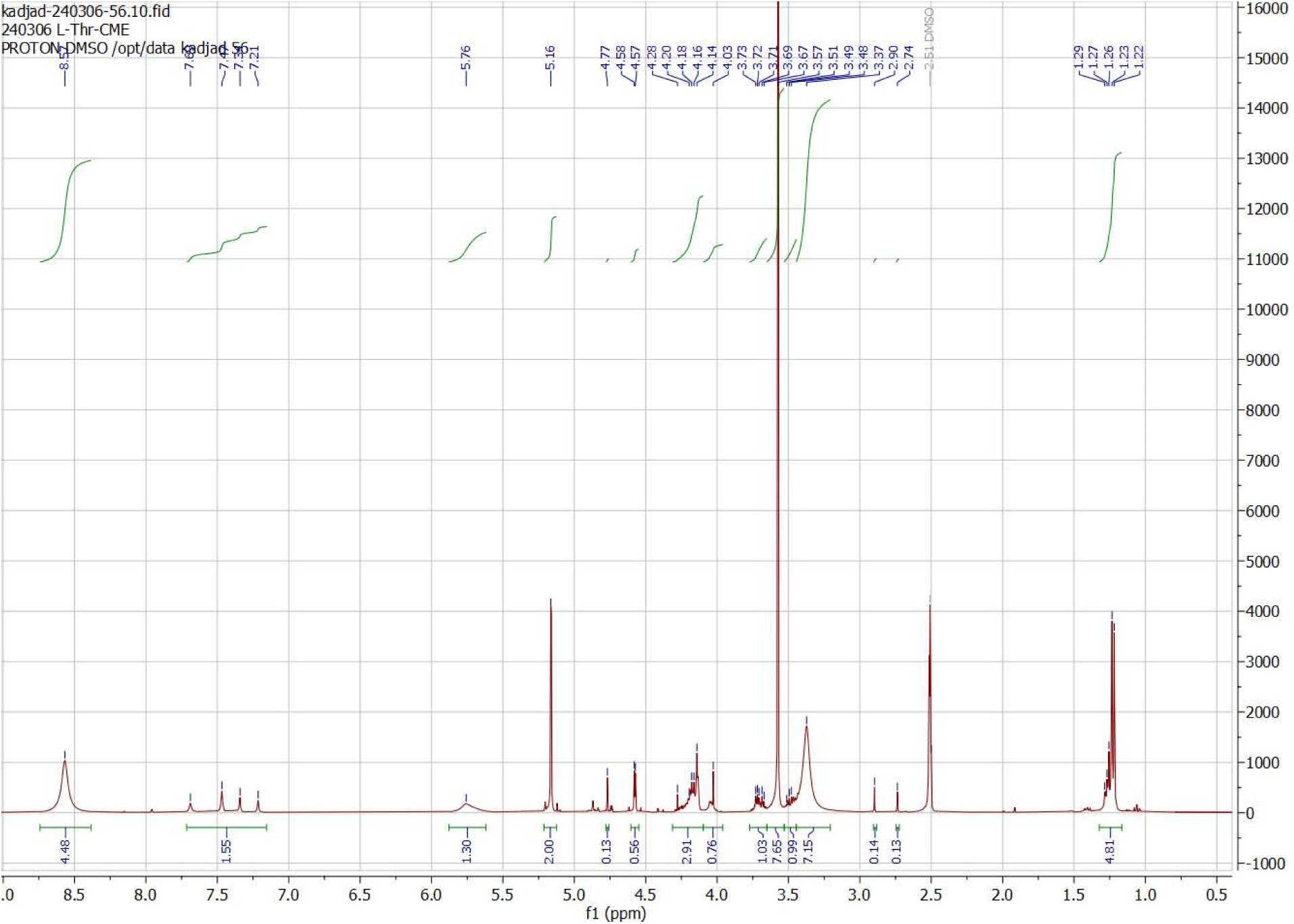
^1^H NMR of L-threonine-cyanomethylester

**Figure S62.**
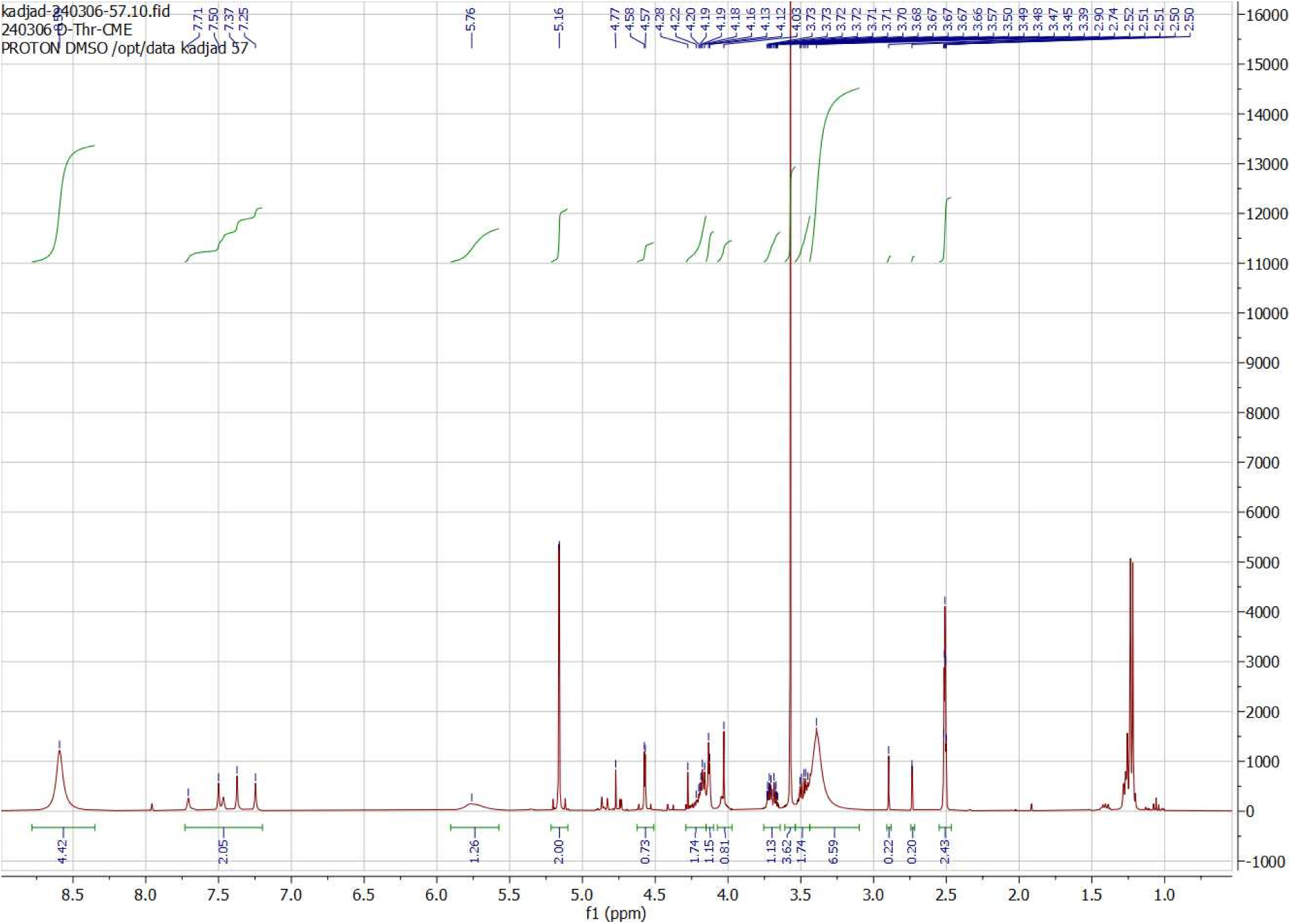
^1^H NMR of D-threonine-cyanomethylester

**Figure S63.**
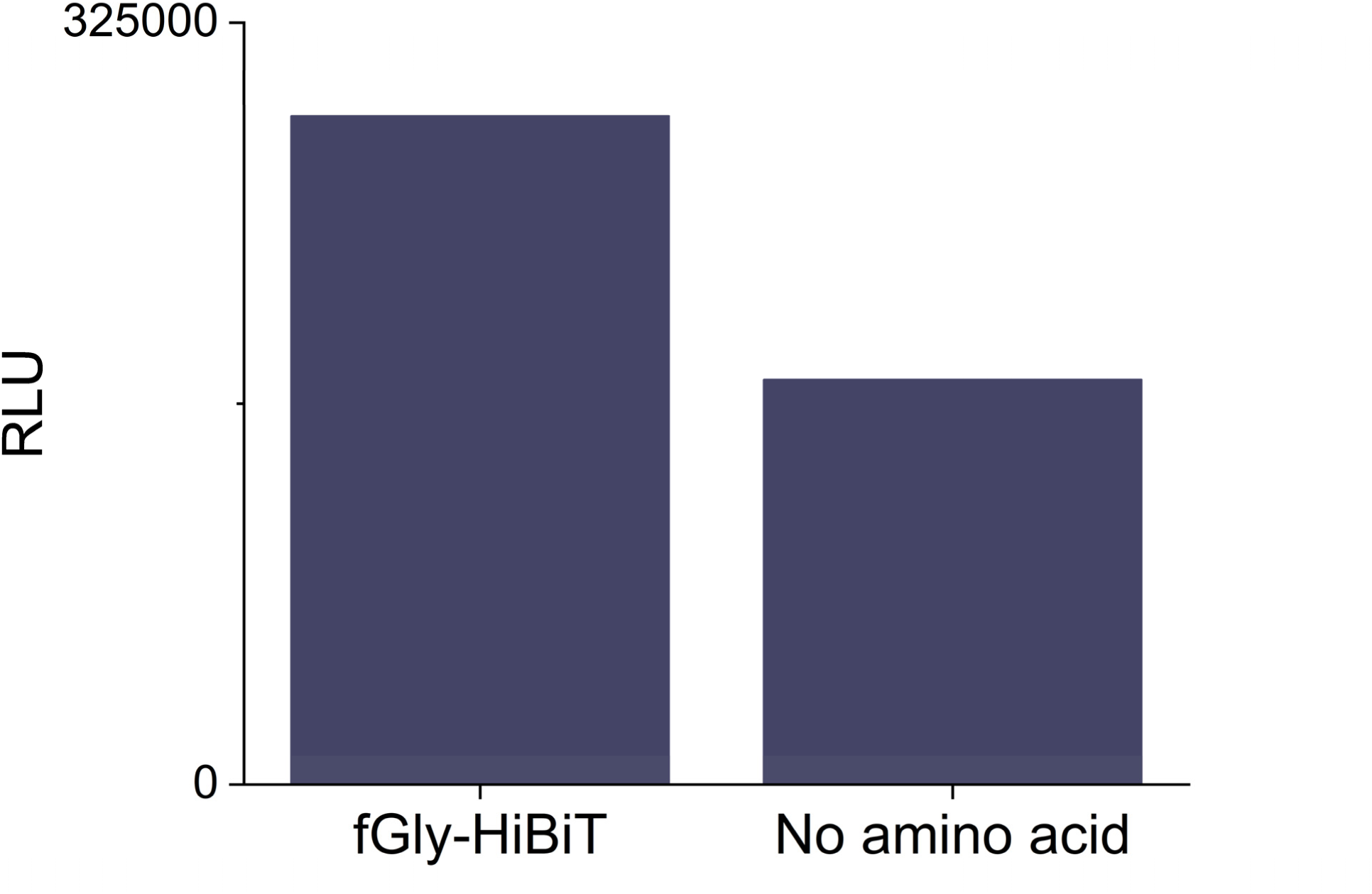
HiBiT luminescence from an in vitro translation reaction using tRNA^fMet^ aminoacylated with glycine-CME

**Figure S64.**
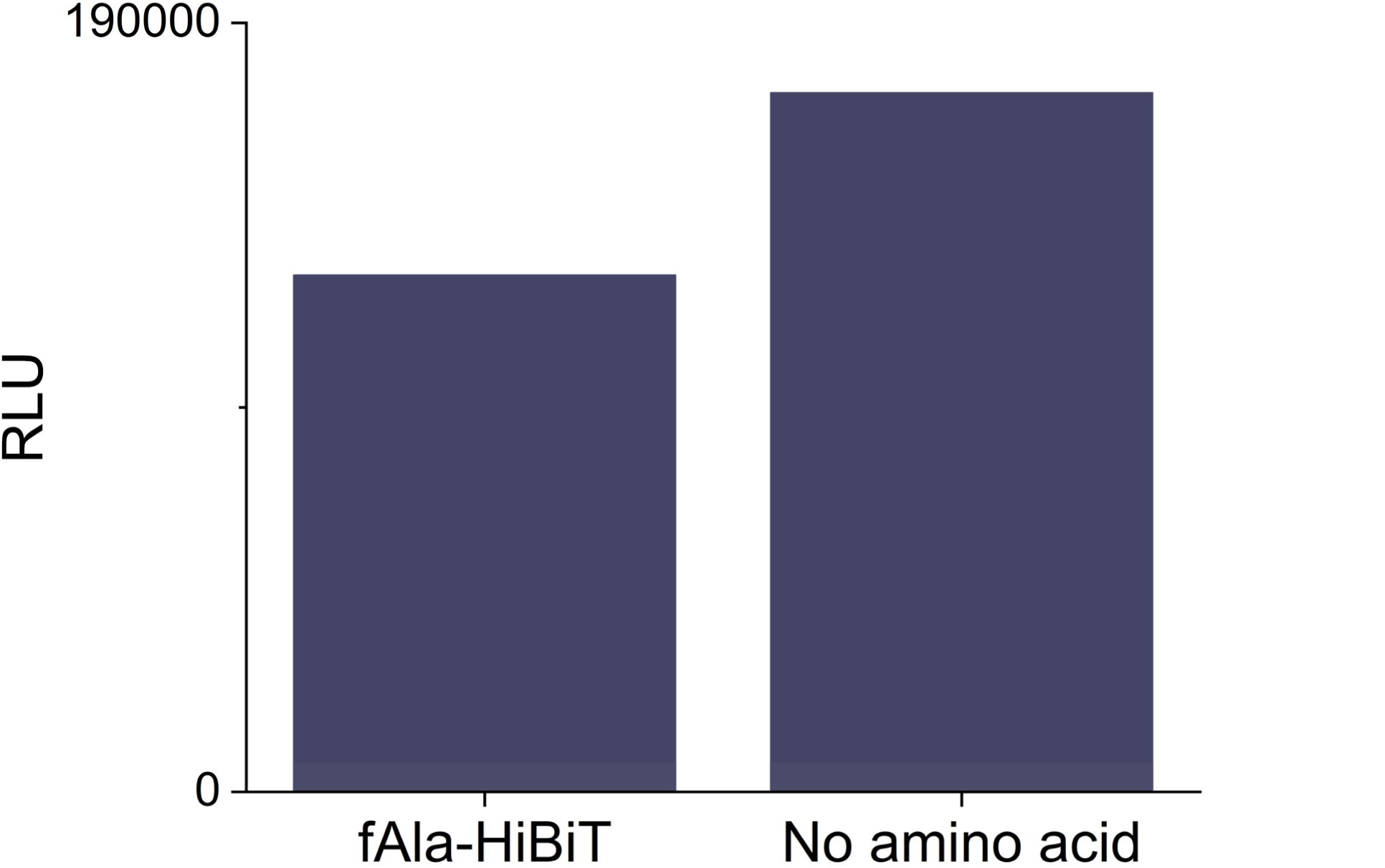
HiBiT luminescence from an in vitro translation reaction using tRNA^fMet^ aminoacylated with alanine-CME

**Figure S65.**
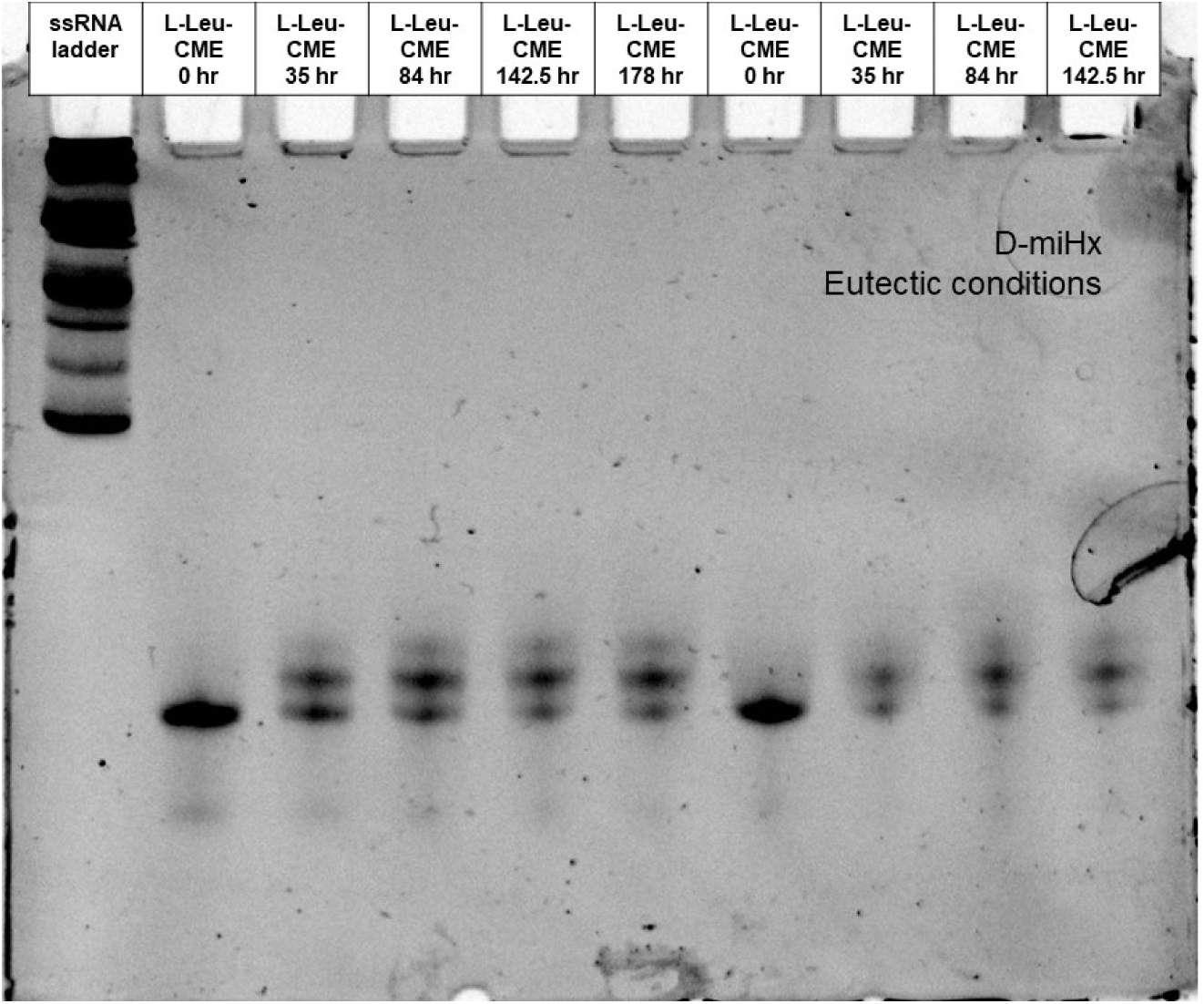
Gel of eutectic nonenzymatic acylation time course using D-miHx RNA to charge with L-Leu-CME at pH 7.5.

**Figure S66.**
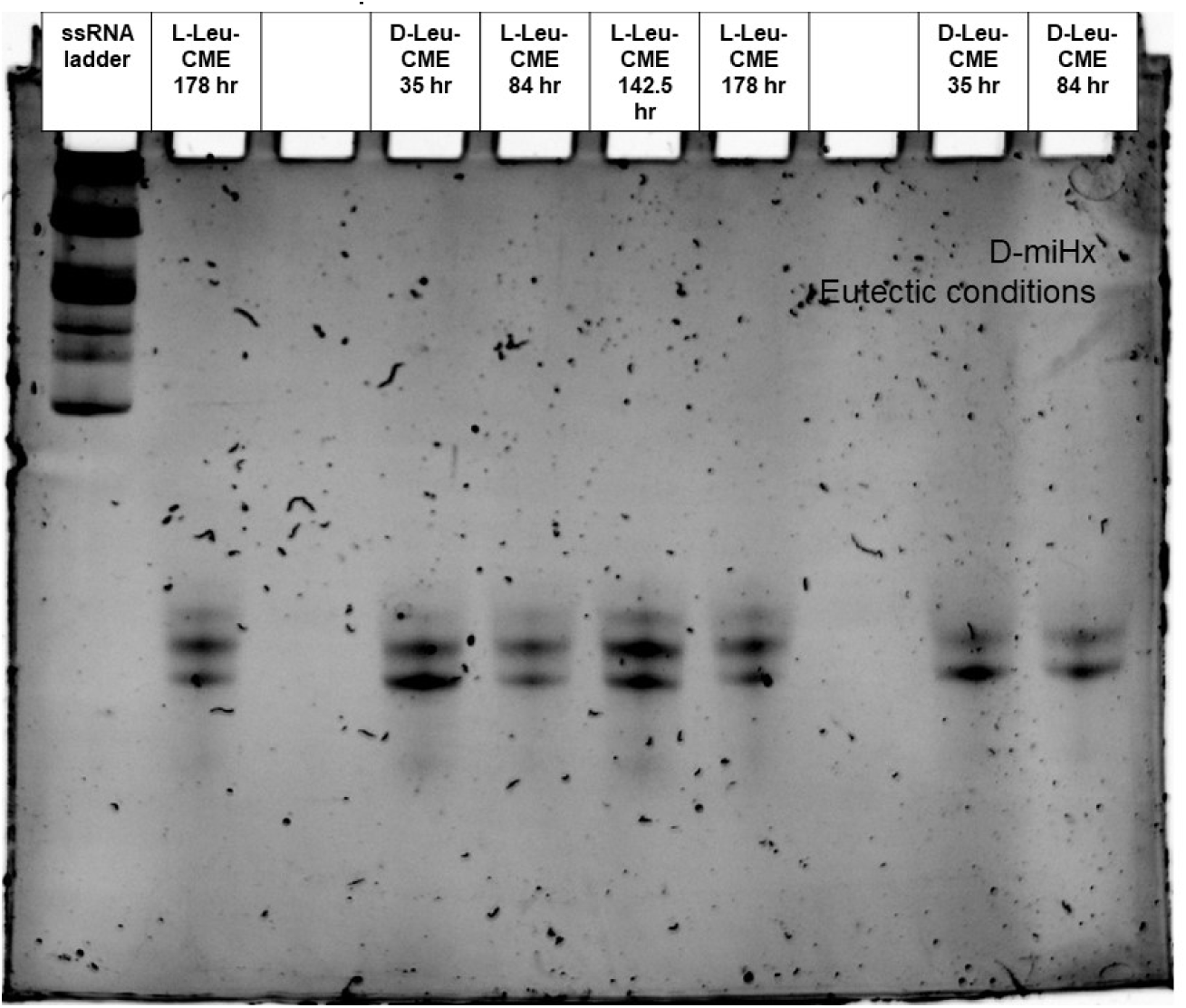
Gel of eutectic nonenzymatic acylation time course using D-miHx RNA to charge with L or D-Leu-CME at pH 7.5.

**Figure S67.**
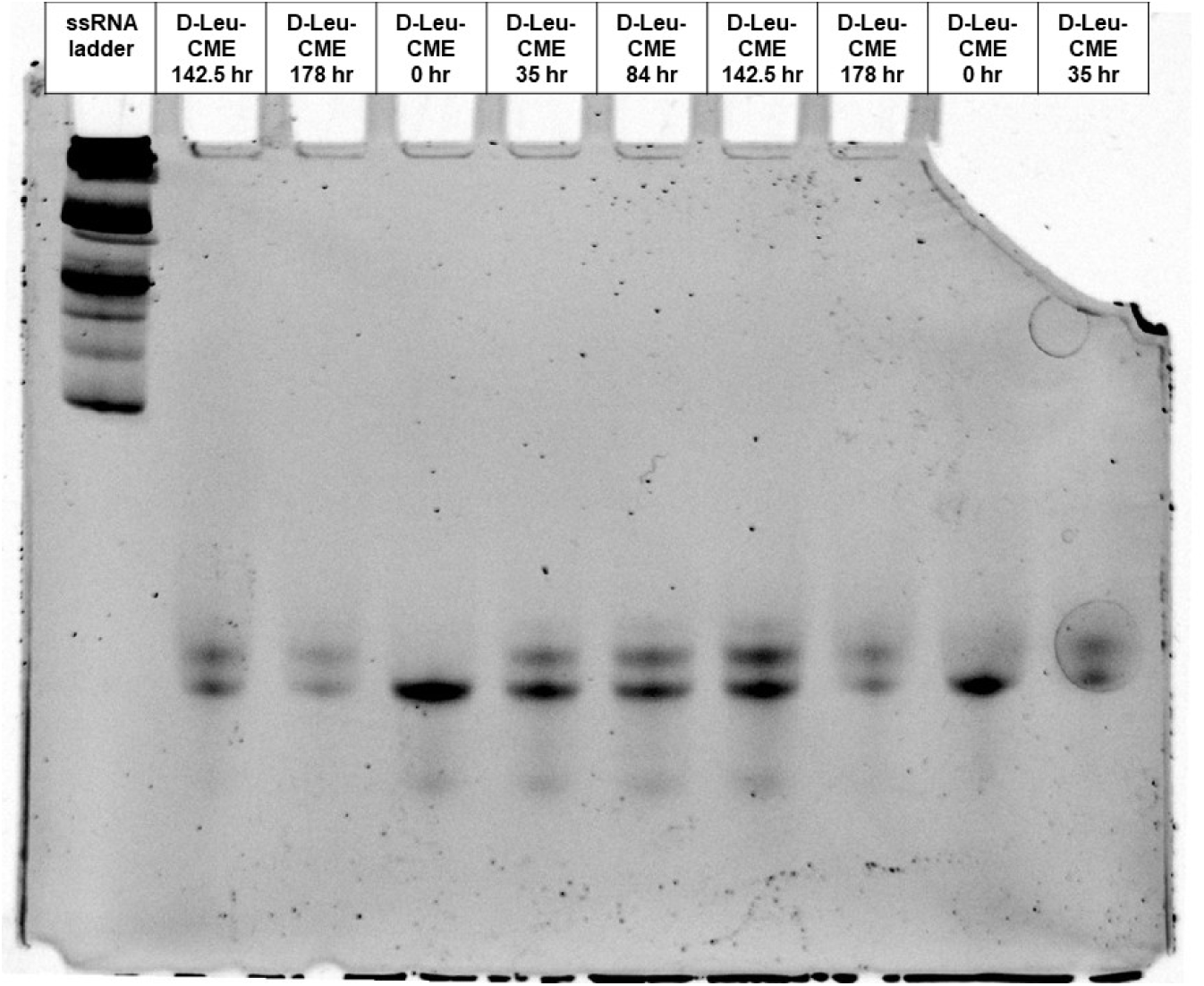
Gel of eutectic nonenzymatic acylation time course using D-miHx RNA to charge with D-Leu-CME at pH 7.5.

**Figure S68.**
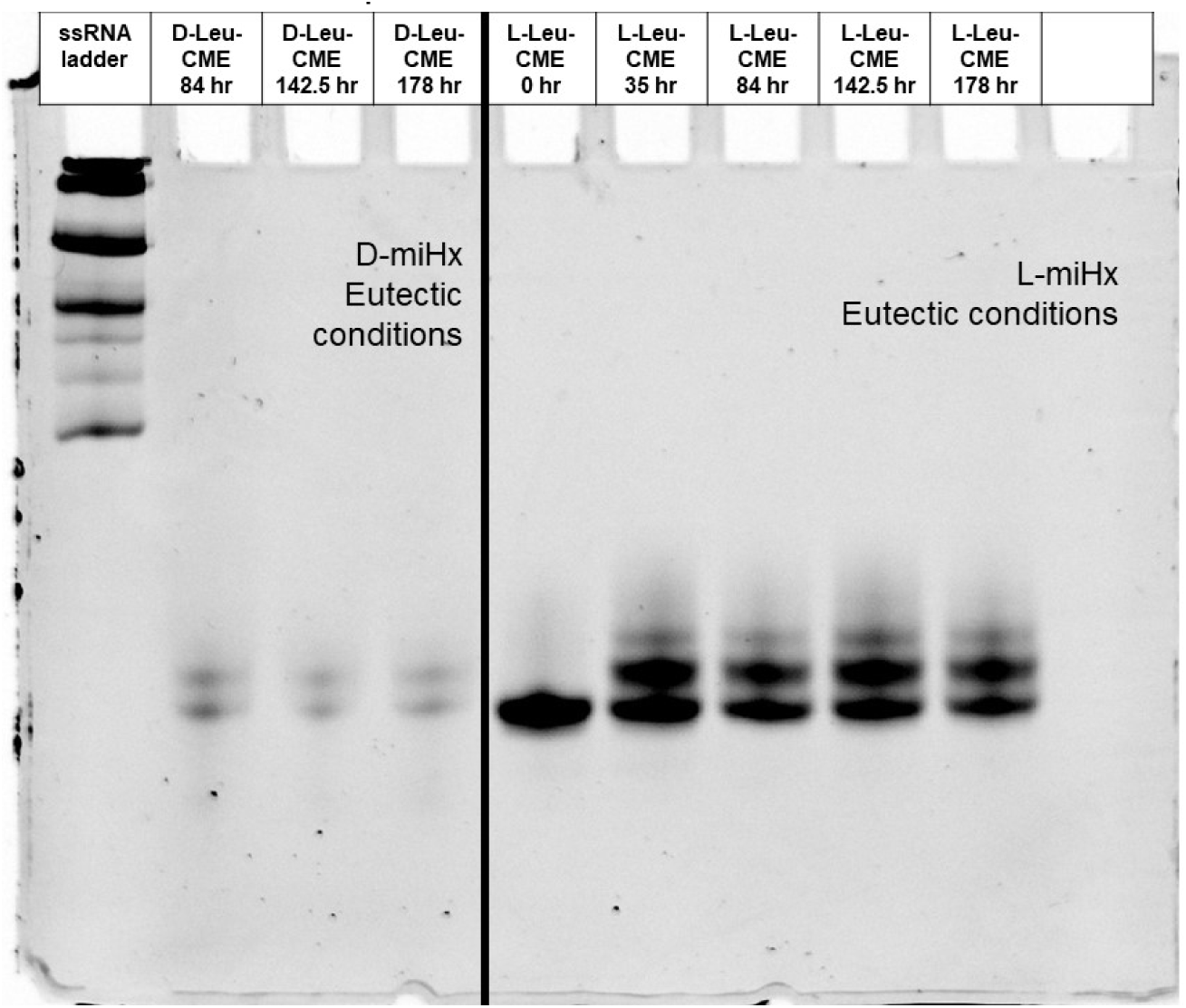
Gel of eutectic nonenzymatic acylation time course using D or L-miHx RNA to charge with L or D-Leu-CME at pH 7.5.

**Figure S69.**
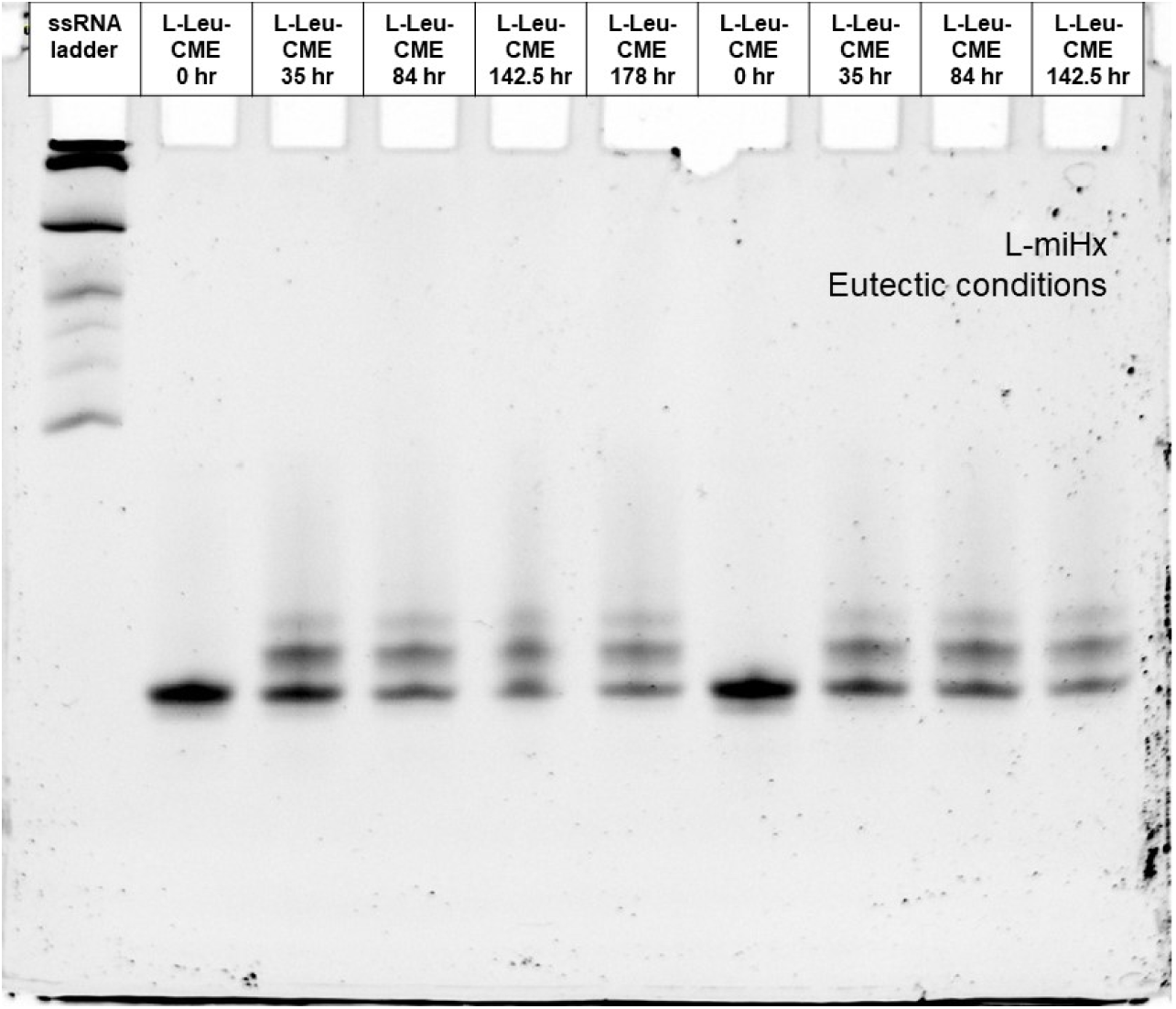
Gel of eutectic nonenzymatic acylation time course using L-miHx RNA to charge with L-Leu-CME at pH 7.5.

**Figure S70.**
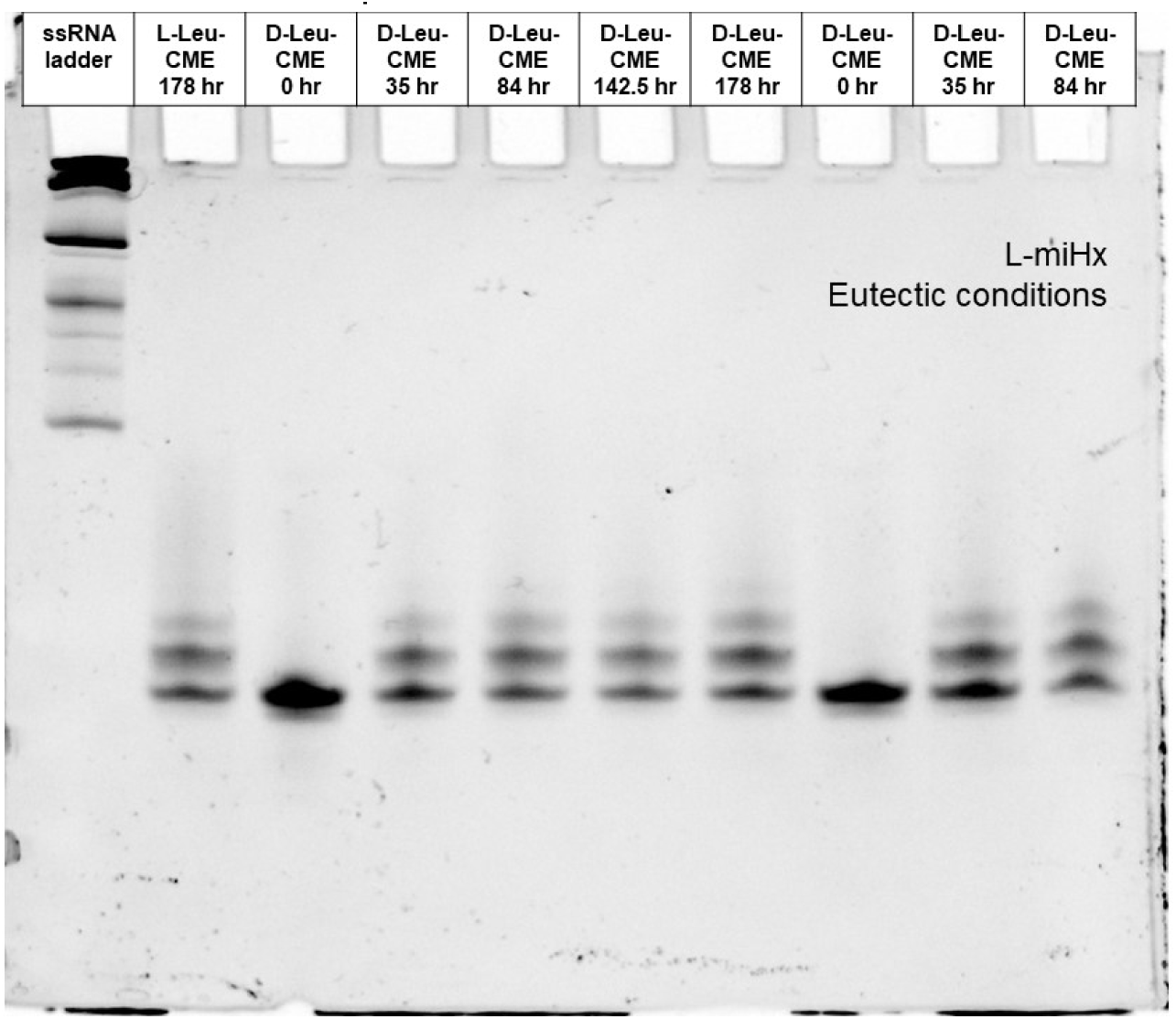
Gel of eutectic nonenzymatic acylation time course using L-miHx RNA to charge with L or D-Leu-CME at pH 7.5.

**Figure S71.**
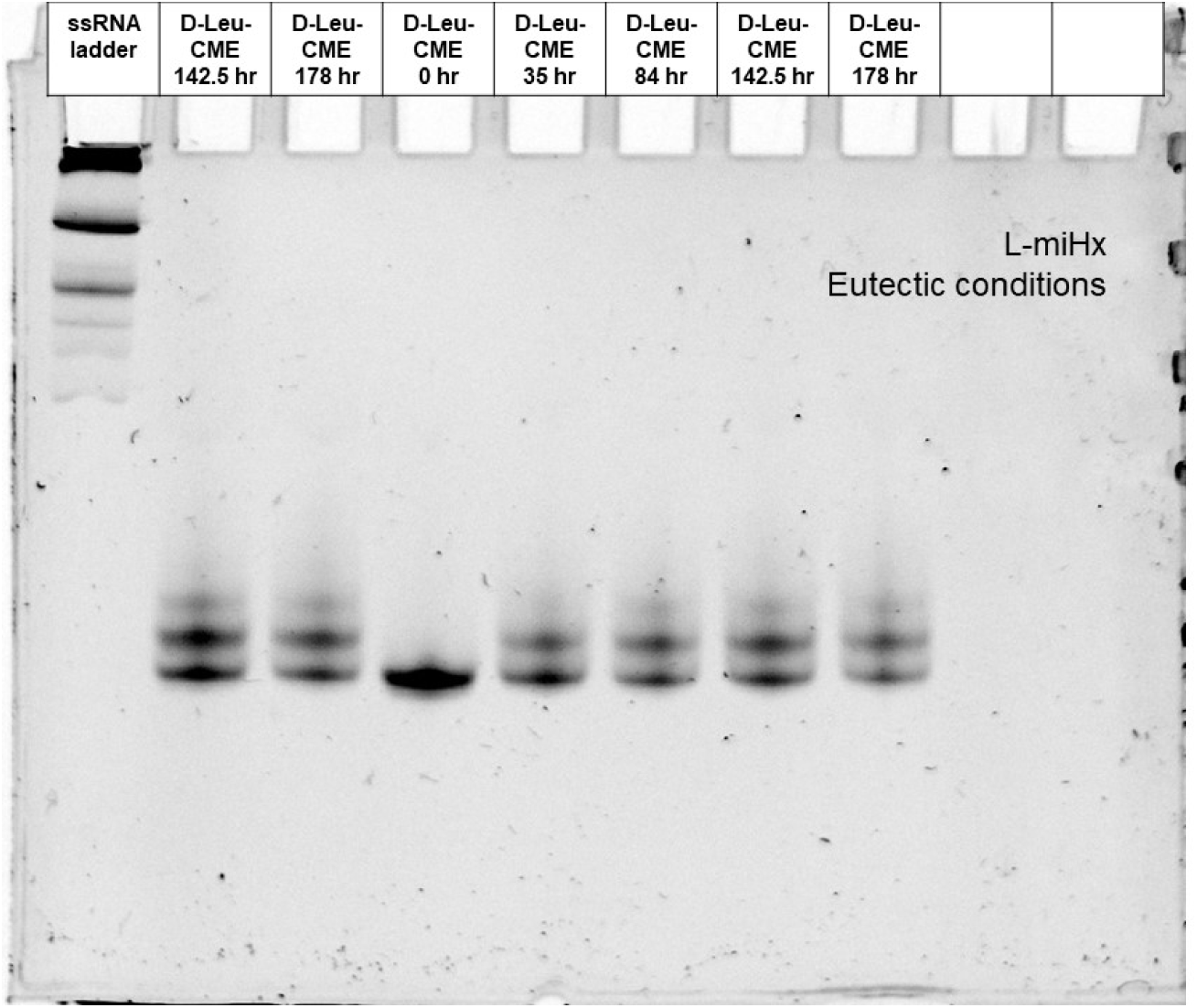
Gel of eutectic nonenzymatic acylation time course using L-miHx RNA to charge with D-Leu-CME at pH 7.5.

**Figure S72.**
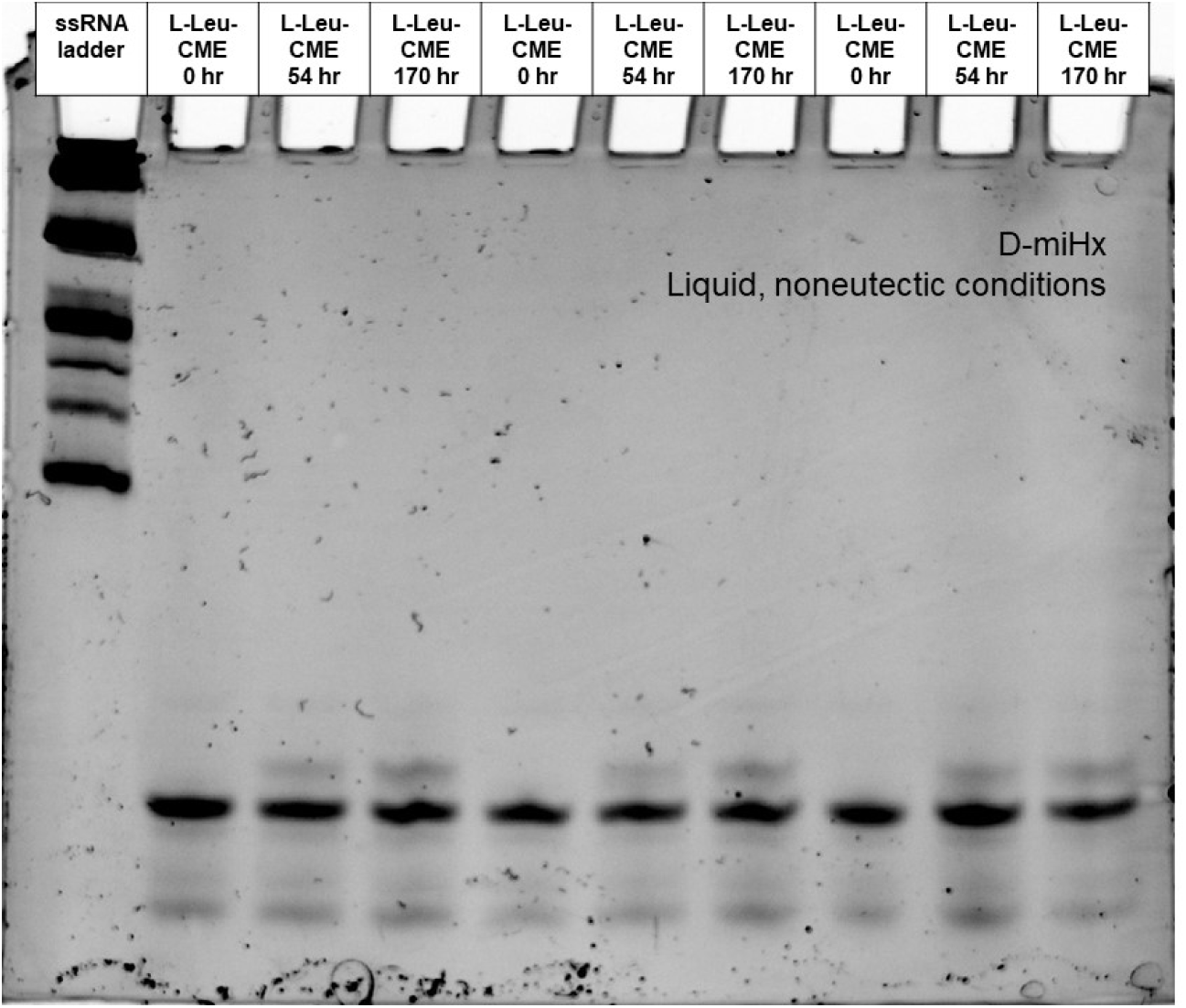
Gel of liquid, noneutectic nonenzymatic acylation time course using D-miHx RNA to charge with L-Leu-CME at pH 7.5.

**Figure S73.**
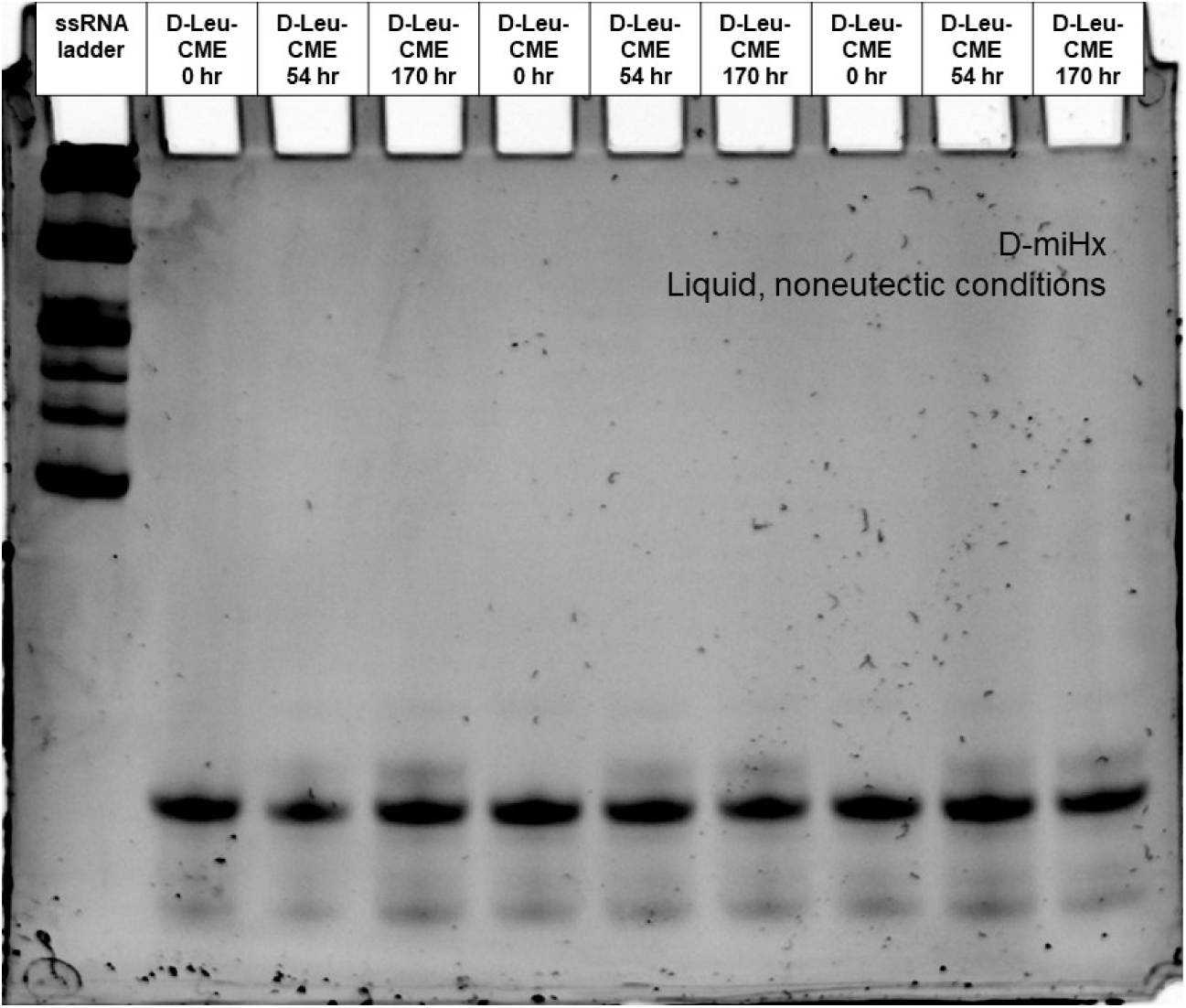
Gel of liquid, noneutectic nonenzymatic acylation time course using D-miHx RNA to charge with D-Leu-CME from at pH 7.5.

**Figure S74.**
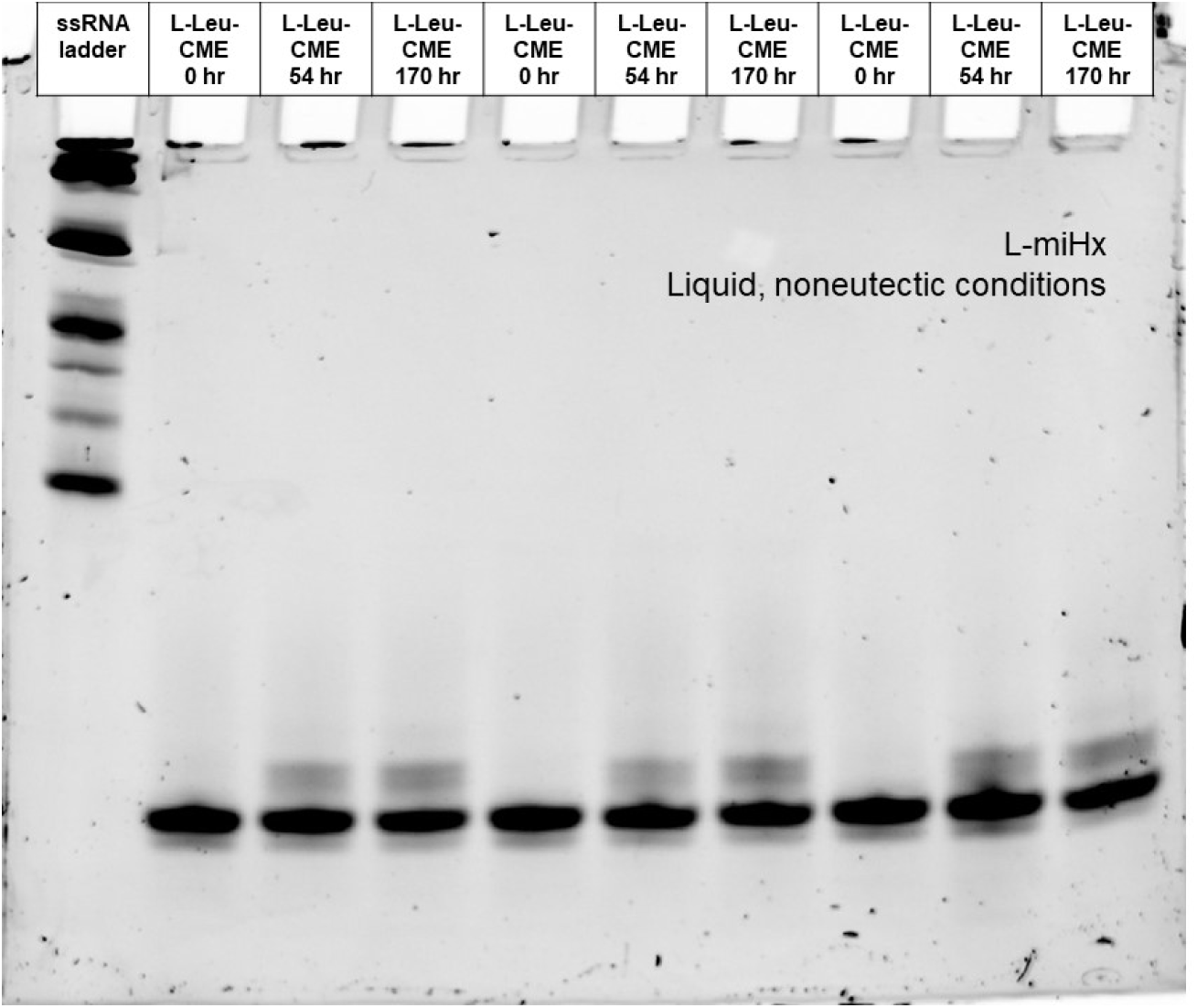
Gel of liquid, noneutectic nonenzymatic acylation time course using L-miHx RNA to charge with L-Leu-CME from at pH 7.5.

**Figure S75.**
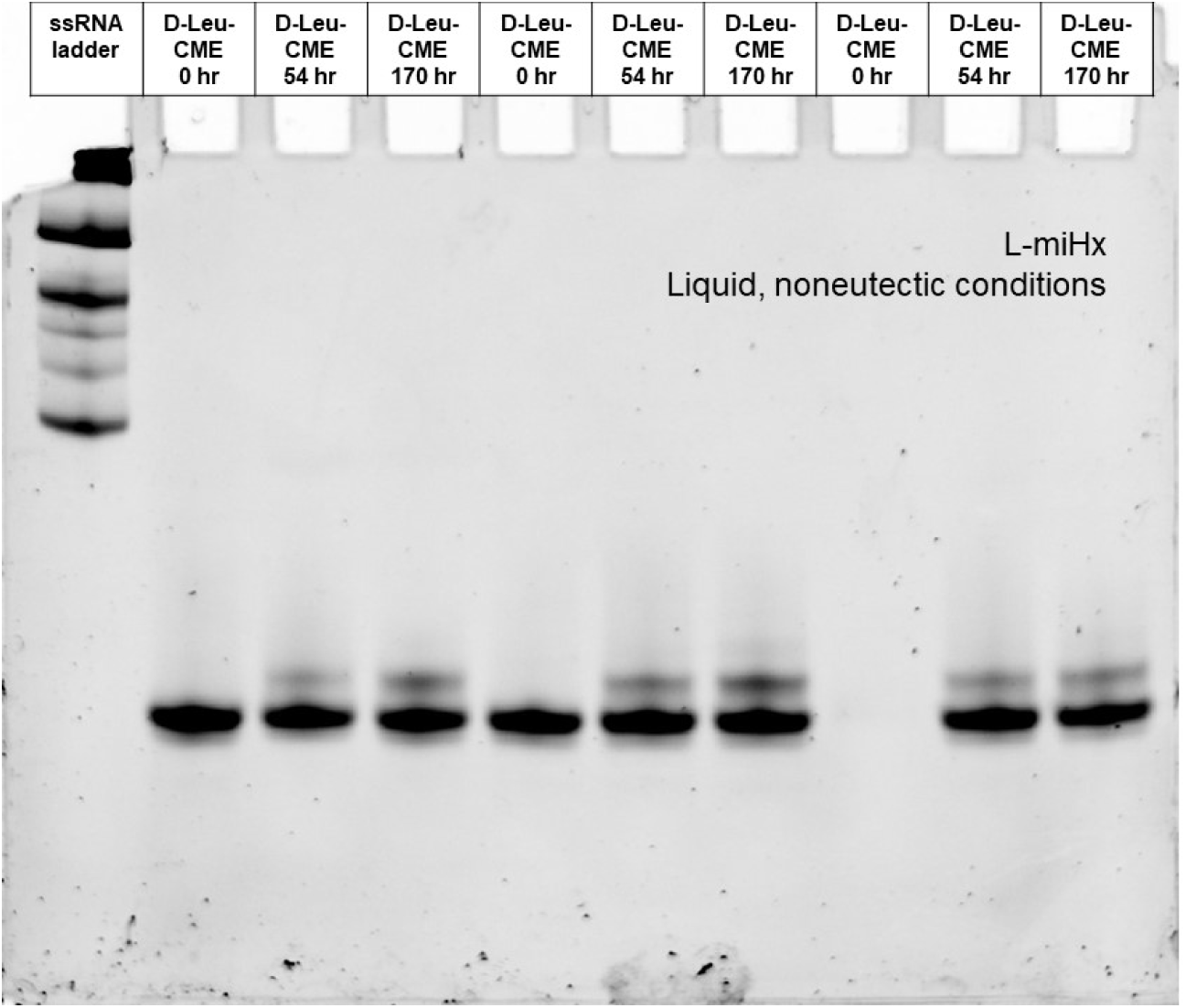
Gel of liquid, noneutectic nonenzymatic acylation time course using L-miHx RNA to charge with D-Leu-CME at pH 7.5.

**Figure S76.**
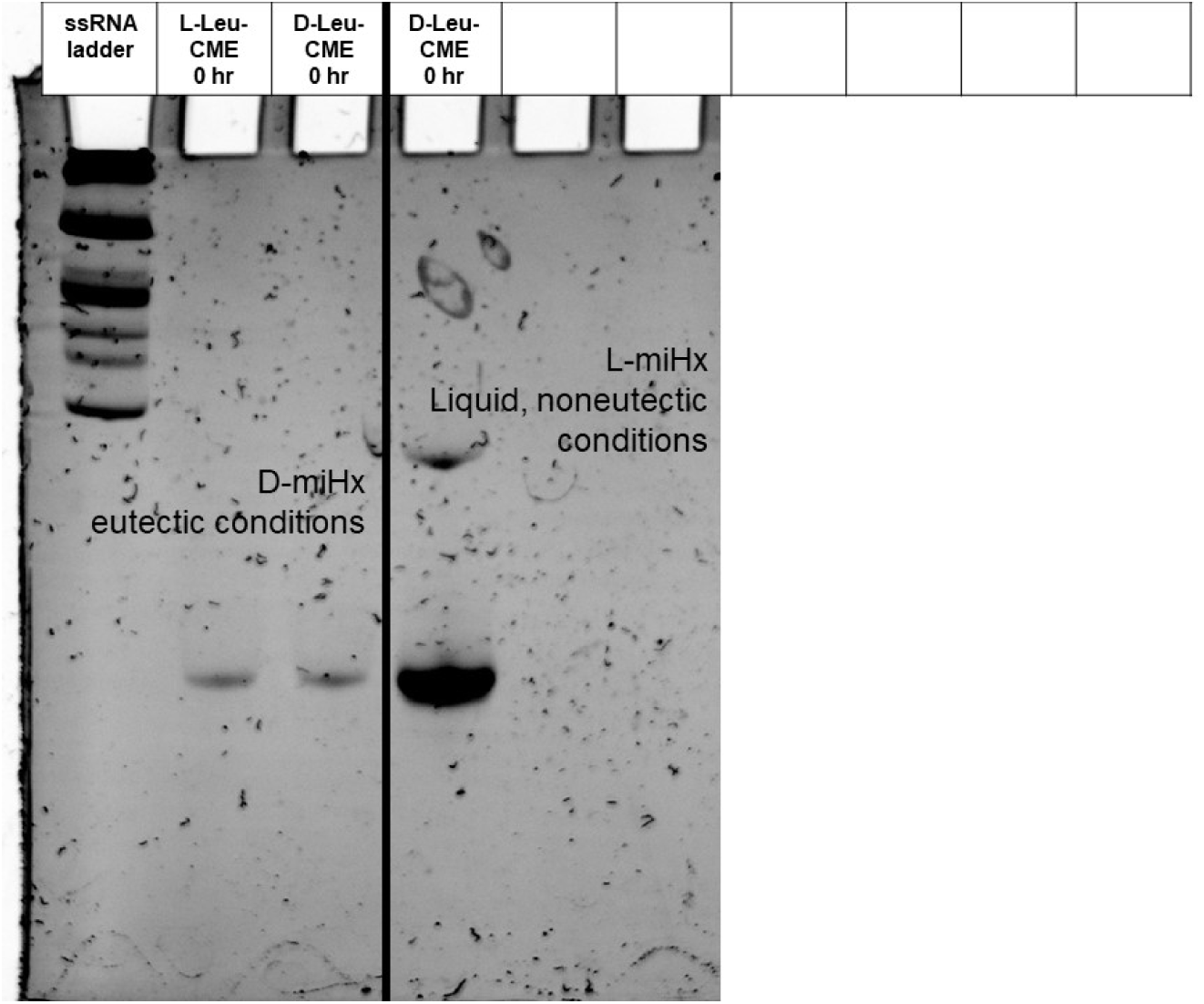
Gel of 0 hr time points for both L and D-Leucine in both eutectic conditions and in liquid, noneutectic nonenzymatic conditions at pH 7.5.

**Figure S77.**
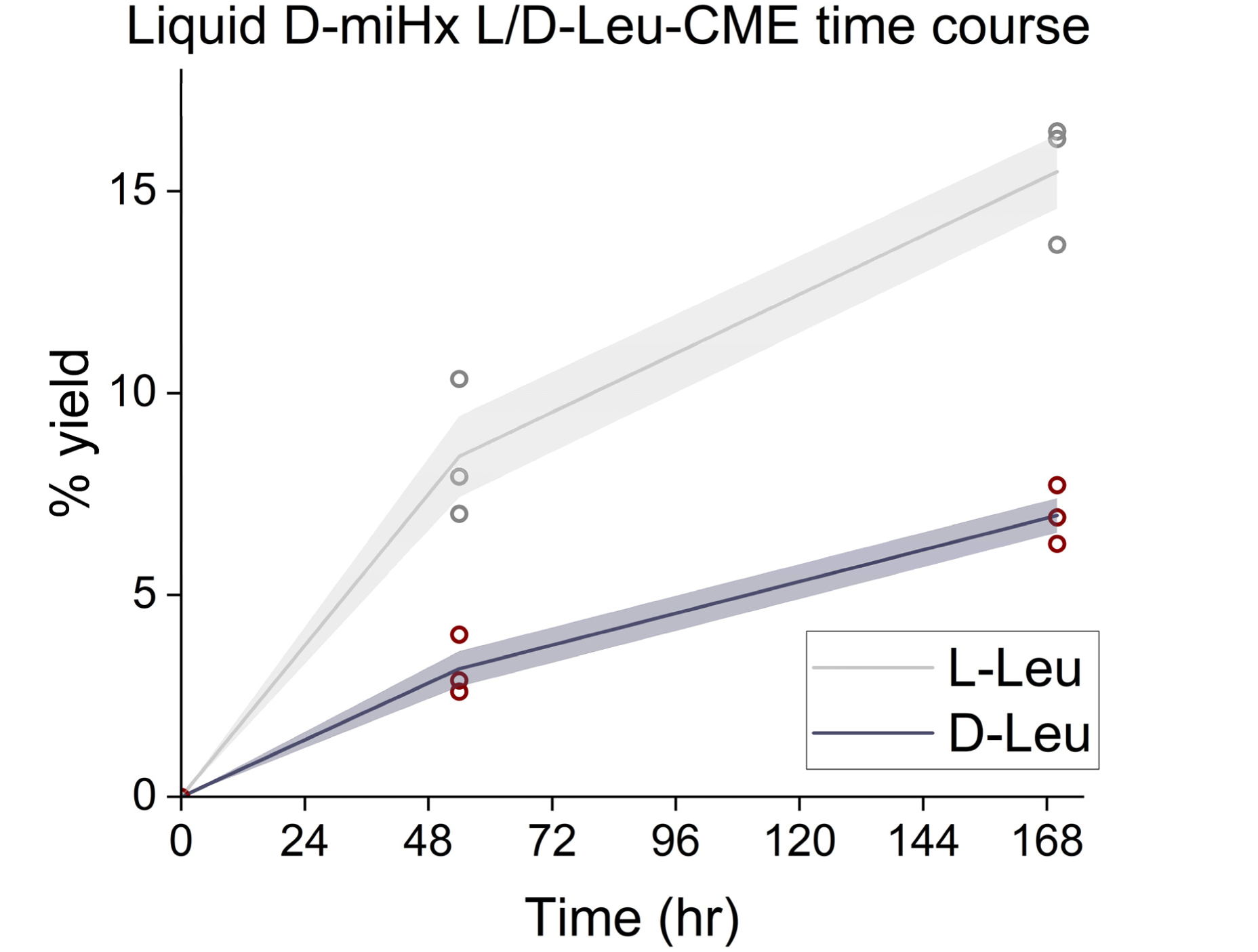
Aminoacylation time course for liquid, noneutectic nonenzymatic acylation reaction using D-miHx RNA to charge with L or D-Leu-CME from 0 to 178 hours at pH 7.5. Gel analysis on supplementary figures S73-74.

**Figure S78.**
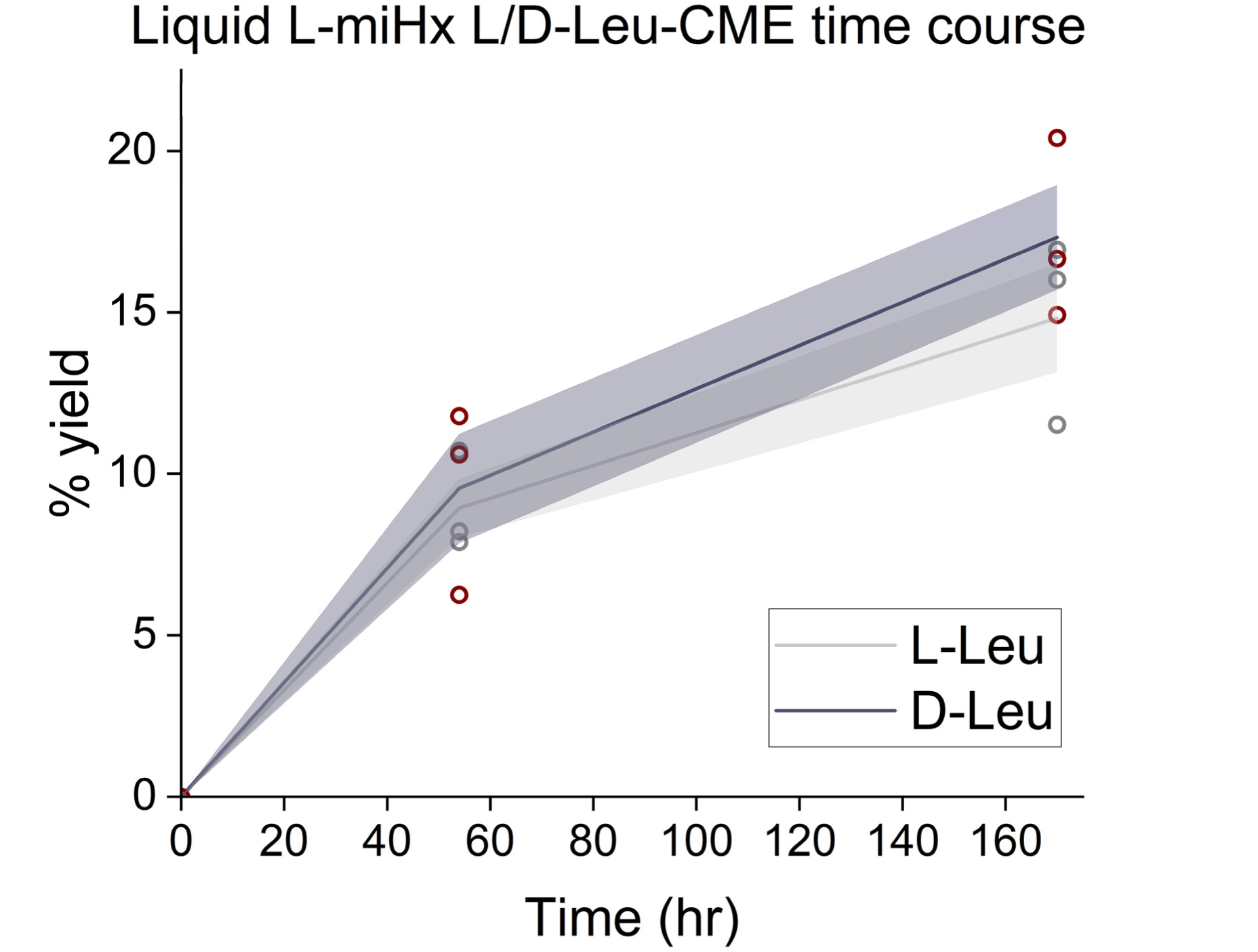
Aminoacylation time course for liquid, noneutectic nonenzymatic acylation reaction using L-miHx RNA to charge with L or D-Leu-CME from 0 to 178 hours at pH 7.5. Gel analysis on supplementary figures S75-76, 80.

**Table S1.**
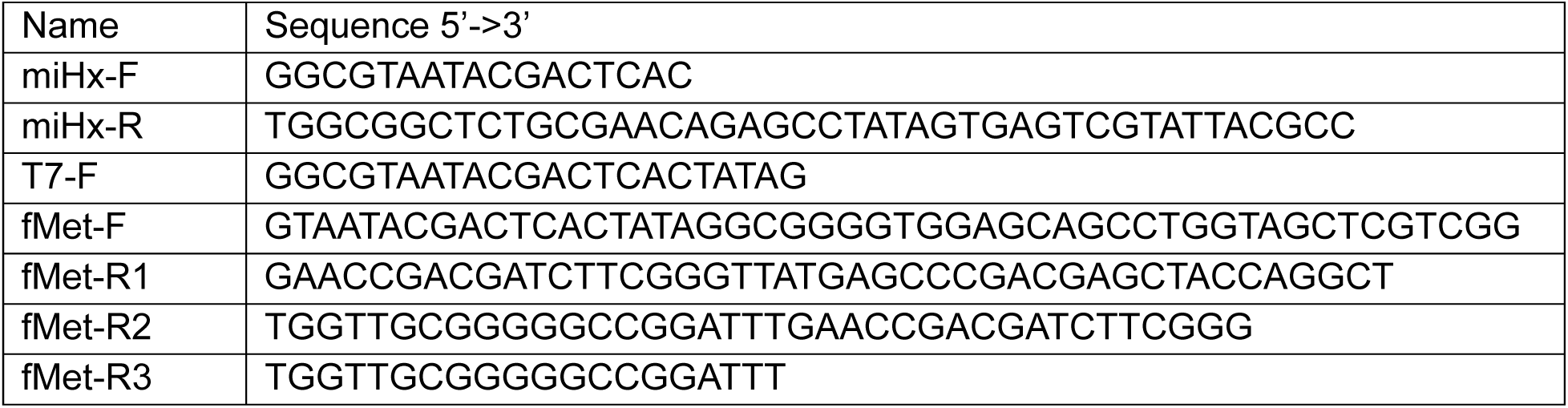
Sequences for primers used to generate DNA templates for *in vitro* transcriptions.

